# Asynchrony between virus diversity and antibody selection limits influenza virus evolution

**DOI:** 10.1101/2020.04.27.064915

**Authors:** Dylan H. Morris, Velislava N. Petrova, Fernando W. Rossine, Edyth Parker, Bryan T. Grenfell, Richard A. Neher, Simon A. Levin, Colin A. Russell

**Affiliations:** Department of Ecology & Evolutionary Biology, Princeton University, Princeton, NJ 08544, USA; Department of Human Genetics, Wellcome Trust Sanger Institute, Cambridge, UK; Department of Veterinary Medicine, University of Cambridge, Cambridge, UK; Department of Medical Microbiology, Academic Medical Center, University of Amsterdam, Amsterdam, NL; Fogarty International Center, National Institutes of Health, Bethesda, MD 20892, USA; Biozentrum, University of Basel, Basel, CH

## Abstract

Seasonal influenza viruses create a persistent global disease burden by evolving to escape immunity induced by prior infections and vaccinations. New antigenic variants have a substantial selective advantage at the population level, but these variants are rarely selected within-host, even in previously immune individuals. Using a mathematical model, we show that the temporal asynchrony between within-host virus exponential growth and antibody-mediated selection could limit within-host antigenic evolution. If selection for new antigenic variants acts principally at the point of initial virus inoculation, where small virus populations encounter well-matched mucosal antibodies in previously infected individuals, there can exist protection against reinfection that does not regularly produce observable new antigenic variants within individual infected hosts. Our results provide a theoretical explanation for how virus antigenic evolution can be highly selective at the global level but nearly neutral within host. They also suggest new avenues for improving influenza control.

## Introduction

Antibody-mediated immunity exerts evolutionary selection pressure on the antigenic phenotype of seasonal influenza viruses (Archetti & Horsfall, 1950; Hensley et al., 2009). Influenza virus infections and vaccinations induce neutralizing antibodies that can prevent re-infection with previously encountered virus antigenic variants, but such re-infections nonetheless occur (Clements et al., 1986; Javaid et al., 2020; Memoli et al., 2019). At the human population level, accumulation of antibody-mediated immunity creates selection pressure favoring antigenic novelty. Circulating antigenic variants typically go rapidly extinct following the population-level emergence of a new antigenic variant, at least for A/H3N2 viruses (Smith et al., 2004).

New antigenic variants like those that result in antigenic cluster transitions (Smith et al., 2004) and warrant updating the composition of seasonal influenza virus vaccines are likely to be produced in every infected host. Seasonal influenza viruses have high polymerase error rates (on the order of 10^−5^ mutations/nucleotide/replication (Nobusawa & Sato, 2006)), reach large within-host virus population sizes (as many as 10^10^ virions (Perelson et al., 2012)), and can be altered antigenically by single amino acid substitutions in the hemagglutinin (HA) protein (Koel et al., 2013; Linderman et al., 2014).

In the absence of antibody-mediated selection pressure, *de novo* generated antigenic variants should constitute a tiny minority of the total within-host virus population. Such minority variants are unlikely to be transmitted onward or detected with current next-generation sequencing (NGS) methods. But selection pressure imposed by the antibody-mediated immune response in previously exposed individuals could promote these variants to sufficiently high frequencies to make them easily transmissible and NGS detectable. The potential for antibody-mediated antigenic selection can be readily observed in infections of vaccinated mice (Hensley et al., 2009) and in virus passage in eggs in the presence of immune sera (Davis et al., 2018).

Surprisingly, new antigenic variants are rarely observed in human seasonal influenza virus infections, even in recently infected or vaccinated hosts (Debbink et al., 2017; Dinis et al., 2016; Han et al., 2019; Javaid et al., 2020; Leonard et al., 2016; McCrone et al., 2018; Valesano et al., 2019) (Fig. 1A,B). These observations contradict existing models of within-host influenza virus evolution (Luo et al., 2012; Volkov et al., 2010) and pathogen immune escape generally (Kennedy & Read, 2017), which model strong within-host antibody selection from the beginning of infection and therefore predict that new antigenic variants will be at consensus or fixation in detectable reinfections of previously immune hosts. This raises a fundamental dilemma. If within-host antibody selection is strong, why do new antigenic variants appear so rarely? If this selection is weak, how can there be protection against reinfection and resulting strong population-level selection?

**Fig. 1.**
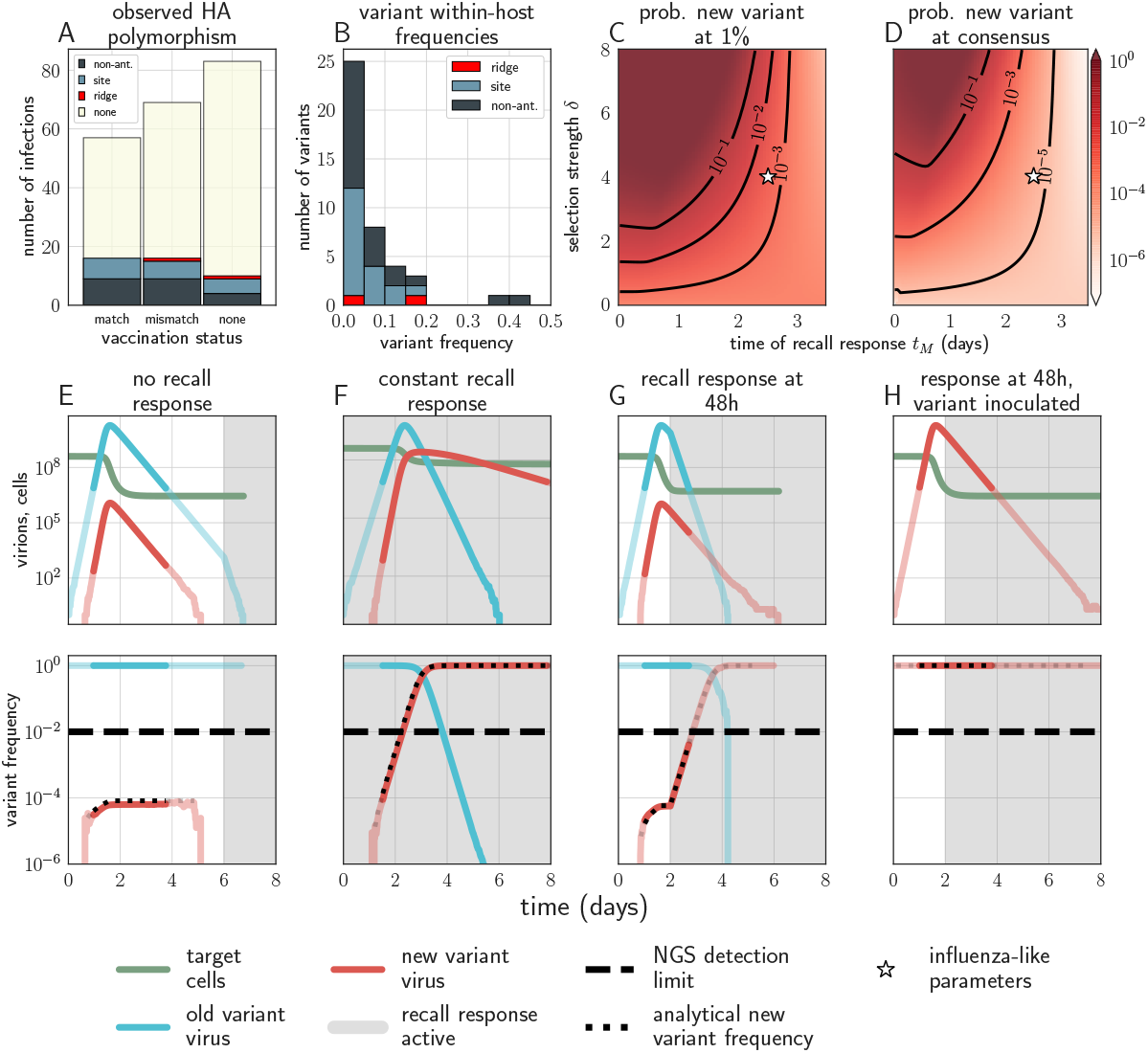
Empirical within-host influenza virus variant frequencies and model within-host evolutionary dynamics. (**A, B**) meta-analysis of A/H3N2 viruses from next-generation sequencing studies of naturally infected individuals (Debbink et al., 2017; McCrone et al., 2018). (**A**) Fraction of infections with one or more observed amino acid polymorphisms in the hemagglutinin (HA) protein, stratified by likelihood of affecting antigenicity: infections with a substitution in the “antigenic ridge” of 7 key amino acid positions found by (Koel et al., 2013) in red, infections with a substitution in a classically defined “antigenic site”, (Wiley et al., 1981) in blue, infections with HA substitutions only in non-antigenic regions in grey, infections with no HA substitutions in cream. Infections grouped by whether individuals had been (left) vaccinated in a year that the vaccine matched the circulating strain, (center) vaccinated in a year that the vaccine did not match the circulating strain, or (right) not vaccinated. (**B**) Distribution of plotted polymorphic sites from (**A**) by within-host frequency of the minor variant. (**C, D**) heatmaps showing model probability new antigenic variant selection to the NGS detection threshold of 1% (**C**) and to 50% (**D**) by 3 days post infection given the strength of immune selection *δ*, the antibody response time *t_M_* and an old variant founding population; calculated from equation 27 in the Methods. Calculated with *c_w_* = 1, *c_m_* = 0, but for *t_M_* > 1, replication selection probabilities are approximately equal for all *c_w_*, *c_m_*, *k* trios that yield a given *δ* (see Methods). Star denotes a plausible influenza-like parameter regime: 25% escape from sterilizing-strength immunity (*c_w_* = 1, *c_m_* = 0.75, *k* = 20) with a recall response at 2.5 days post infection. Black lines are probability contours. (**E–H**) example model trajectories. Upper row: absolute counts of virions and target cells. Lower row: variant frequencies for old antigenic variant (blue) and new variant (red). Dashed line shows 1% frequency, the detection limit of NGS. Dotted line shows an analytical prediction for new variant frequency according to equations 15 and 16 (see Methods). Model scenarios: (**E**) naive; (**F**) experienced with *t_M_* = 0; (**G**) experienced with *t_M_* = 2; (**H**) experienced with *t_M_* = 2 and new antigenic variant virion incoulated. Lines faded when infection is below 5% transmission probability—approximately 10^7^ virions with default parameters. All parameters as in Table 1 unless otherwise stated.

We hypothesized that influenza virus antigenic evolution is limited by the asynchrony between virus diversity and antibody-mediated selection pressure. Antibody immunity at the point of transmission in previously infected or vaccinated individuals should reduce the initial probability of reinfection (Le Sage et al., 2020); secretory IgA antibodies on mucosal surfaces (sIgA) are likely to play a large role ((Wang et al., 2017), see Appendix section 2). But if viruses are not blocked at the point of transmission and successfully infect host cells, an antibody-mediated recall response takes multiple days to mount (Coro et al., 2006; Lam & Baumgarth, 2019) (see detailed review in the Appendix, section 2). Virus titer—and virus shedding (Lau et al., 2010)—may peak before the production of new antibodies has even begun, leaving limited opportunity for within-host immune selection. If antigenic selection pressure is strong at the point of transmission but weak during virus exponential growth, new antigenic variants could spread rapidly at the population level without being readily selected during the timecourse of a single infection. Moreover, prior work has established tight population bottlenecks at the point of influenza transmission (McCrone et al., 2018; Xue & Bloom, 2019). With a tight transmission bottleneck and weak selection during virus exponential growth, antigenic diversity generated during any particular infection will most likely be lost, slowing the accumulation of population-level antigenic diversity.

We used a mathematical model to investigate hypothesis that realistically timed antibody-mediated immune dynamics slow within-host antigenic evolution. We found three modeling results: 1. antibody neutralization at the point of inoculation can protect experienced hosts against reinfection and explain new antigenic variants’ population level selective advantage. 2. If successful reinfection occurs, the delay between virus replication and the mounting of a recall antibody response renders within-host antigenic evolution nearly neutral, even in experienced hosts. 3. It is therefore reasonable that substantial population immunity may need to accumulate before new antigenic variants are likely to observed in large numbers at the population level, whereas effective within-host selection would predict that they should be readily observable even before they proliferate.

Our modeling results suggest a plausible mechanism that can explain otherwise poorly reconciled empirical patterns, and should motivate further experimental investigation of the mechanisms of immune protection and natural selection on influenza virus antigenic phenotypes at the point of transmission.

## Model overview

Our model reflects the following biological realities: 1. Seasonal influenza virus infections of otherwise healthy individuals typically last 5–7 days (Suess et al., 2012); 2. In influenza virus-naive individuals, it can take up to 7 days for anti-influenza virus antibodies to start being produced (Wrammert et al., 2008), effectively resulting in no selection (Fig. 1A); 3. In previously infected (“experienced”) individuals, sIgA antibodies can neutralize inoculated virions before they can infect host cells; (Wang et al., 2017) 4. However, if an inoculated virion manages to cause an infection in an experienced individual, it takes 2–5 days for the infected host to mount a recall adaptive immune response, including producing new antibodies (Coro et al., 2006; Zuccarino-Catania et al., 2014) (see Appendix section 2 for further discussion of motivating immunology). Importantly, this contrasts with previous within-host models of virus evolution, which have assumed that antibody-mediated neutralization of virions during virus replication is strong from the point of inoculation onward and is the mechanism of protection against reinfection (Luo et al., 2012; Volkov et al., 2010), and reflects new animal model evidence of sterilizing antibody immunity (Le Sage et al., 2020). We discuss existing models and hypotheses for the rarity of population level influenza antigenic variation in section 7 of the Appendix.

We constructed a model of within-host influenza virus evolution that can be parameterized to reflect different hypothesized immune mechanisms, different host immune statuses, and different durations of infection. In the model, virions *V_i_* of antigenic type *i* infect target cells *C*, replicate, mutate to a new antigenic type *j* at a rate *μ_ij_*, and decay at a rate *d_v_*. We model the innate immune response implicitly as depletion of infectible cells. We model the antibody-mediated immune response as an increase *k* in the virion decay rate in the presence of well-matched antibodies. To model partial antibody cross reactivity, we scale *k* by a parameter *c_i_* ∈ [0, 1]; *c_i_* reflects the binding strength of the host’s best-matched antibodies to antigenic type *i*. So in the presence of an antibody-mediated response, virions *V_i_* of type *i* decay at a rate *d_v_* + *c_i_k*.

The model can accommodate *N_v_* antigenic variants *i* =1, 2, … *N_v_* linked by an arbitrary network of possible substitutions and corresponding mutation probabilities *μ_ij_*, but in practice we typically consider two, the new variant *m* and the old variant *w*, and neglect back-mutation from new variant to old variant (*μ_wm_* > 0, but *μ_mw_* = 0).

To assess the importance of transmission bottlenecks, initial virus diversity, and sIgA antibody neutralization in virus evolution, we model the point of transmission as a series of stochastic events which may ultimately lead to one of more virions invading cells and initiating an infection. The recipient host is inoculated with a random sample of within-host virus diversity from a transmitting host. In experienced hosts, this inoculum is probabilistically thinned by host antibodies. The founding population that initiates the infection is then randomly sampled from among any remaining virions.

Mathematically, we model the number of inoculated virions as Poisson-distributed with a mean *v*, so if variant *i* has within-host frequency *f_i_*, the number of variant *i* virions inoculated is Poisson-distributed with mean *vf_i_*. The virions then encounter antibodies, which we interpret as sIgA but can be understood to be any antigen-specific antibody-mediated protection that precedes cell infection; each virion of variant *i* is independently neutralized with a probability *κ_i_*. This probability depends upon the strength of protection against homotypic reinfection *κ* and the sIgA cross immunity between variants *σ_ij_*, (0 ≤ *σ_ij_* ≤ 1). So if a host with antibodies to variant *j* is challenged with variant *i*, those virions will be neutralized with a probability *κ_i_* = *κσ_ij_*. For simplicity, we assume the same homotypic protection level across all variants and hosts, though in practice there may be variation in the immunogenicity of individual variants and in the strength of responses generated by individual hosts. We typically fix host immune histories to test the effect of host immune history on selective dynamics. When necessary, we can model a novel (non-recall) antibody response to a strain *i* by designating the host as experienced to *i* at some time *t_N_* post-infection (see Methods).

The model is continuous-time and stochastic: cell infection, virion production, virus mutation, and virion decay are stochastic events that occur at rates governed by the current state of the system, with exponentially distributed (memoryless) waiting times. The system is approximately well-mixed: we track counts of virions and cells without an explicit account of space within the upper respiratory tract. We treat infected and dead or removed cells implicitly.

Parameterized as in Table 1, the model captures key features of influenza infections: rapid peaks approximately 2 days post infection and slower declines with clearance approximately a week post infection, with faster clearance in experienced hosts.

**Table 1:** Model parameters, default values, and sources/justifications.

| Parameter | Meaning | Units | Value | Source or justification |
| --- | --- | --- | --- | --- |
| $t_M$ | time post-infection of antibody response in experienced hosts | days | 2 | literature (see review in Section 2) |
| $t_N^w$ | time post infection of a novel immune response to the old antigenic variant | days | 6 | literature (see review in Section 2) |
| $p_C$ | per capita growth rate of target cells at low density | $\frac{1}{\text{days}}$ | 0 | ignored on the timescale of a single infection |
| $C_{\max}$ | maximum number of target cells | cells | $4 \times 10^8$ | standard in the modeling literature (Baccam et al., 2006; Hadjichrysanthou et al., 2016; Luo et al., 2012) |
| $\mathcal{R}_0$ | within-host basic reproduction number for the virus | unitless | 5 | empirical fits of target cell models (Hadjichrysanthou et al., 2016) |
| $r_w$ | average number of virions produced by a cell infected with old antigenic variant virus | virions | 100 | literature (Frensing et al., 2016) |
| $r_m$ | average number of virions produced by a cell infected with new antigenic variant virus | virions | 100 | no within-host deleteriousness for new antigenic variants |
| $z_w$ | probability of no infection given homotypic re-inoculation for a host previously exposed to the old antigenic variant | unitless | 0.95 | challenge studies (Clements et al., 1986; Memoli et al., 2019) |
| $\beta$ | rate of infectious contact between virions and target cells per cell per virion | $\frac{1}{\text{virions cells days}}$ | calculated | from $\mathcal{R}_0$ |
| $f$ | number of target cells lost per infectious contact | cells | 1 | one cell lost per cell infection |
| $k$ | additional per virion neutralization rate in the presence of an antibody response | $\frac{1}{\text{days}}$ | 6 | varied to test hypotheses |
| $\kappa_w$ | probability an individual inoculated old antigenic variant virion inoculated into an experienced host is neutralized in the respiratory tract mucosa | unitless | set from $z_w$ | calculated from equation 38 |
| $\kappa_m$ | probability an individual inoculated old antigenic variant virion inoculated into an experienced host is neutralized in the respiratory tract mucosa | unitless | $\sigma \kappa_w$ | reduced relative to $\kappa_m$ by immune escape |
| $c_w$ | percentage cross reactivity during viral replication between host antibodies and old antigenic variant | unitless | 0 or 1 | naive or homotypically reinfected hosts |
| $c_m$ | percentage cross reactivity during viral replication between host antibodies and new antigenic variant | unitless | 0 | full escape variant |
| $\sigma$ | percentage cross immunity at sIgA bottleneck between old antigenic variant and new antigenic variant | unitless | 0 | full escape variant |
| $b$ | size of final / cell infection bottleneck | virions | 1 | NGS studies (McCrone et al., 2018; Xue & Bloom, 2019) |
| $v$ | number of virions encountering sIgA | virions | $10 \times b$ | |
| $V_{50}$ | viral load at which there is a fifty-percent transmission probability | virions | $10^8$ | chosen to give realistic transmission window (Tsang et al., 2015) and based on prior modeling studies (Russell et al., 2012) |
| $\theta$ | transmission threshold for threshold model | virions | $10^7$ | chosen to be consistent with $V_{50}$ |

We give a full mathematical description of the model in the Methods.

In addition to analyzing this within-host model, we explored the between-host and population-level implications of these within-host dynamics using a simple transmission chain model and an SIR-like population-level model, which we describe in the Methods.

## Results

### Realistically-timed immune kinetics limit otherwise rapid adaptation during exponential growth

In our model, sufficiently strong antibody neutralization during the virus exponential growth period can potentially stop replication and block onward transmission, but this mechanism of protection results in detectable new antigenic variants in each homotypic reinfection, since the infection is terminated rapidly unless it generates a new antigenic variant that substantially escapes antibody neutralization (Fig. 2).

**Fig. 2.**
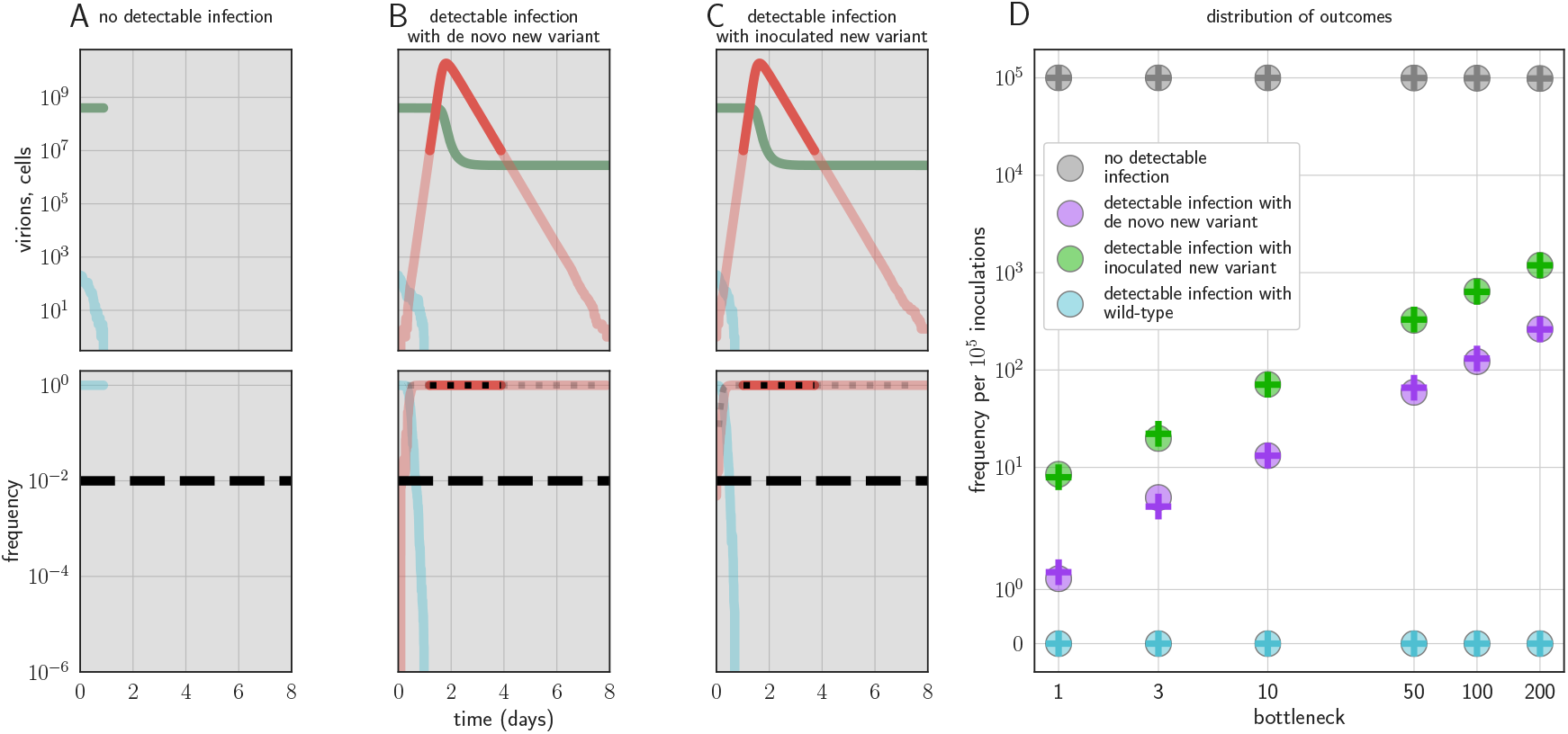
Example timecourses and distribution of outcomes when antibody immunity is active from the start of infection and sufficient to prevent detectable reinfection. *t_M_* = 0, *k* = 20, yielding 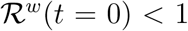 for the old antigenic variant but 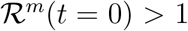 for the new antigenic variant, where 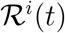 is the within-host effective reproduction number for variant *i* at time *t* (see Methods). No mucosal antibody neutralization (*z_w_* = *z_m_* = 0); protection is only via neutralization during replication Example timecourses from simulations with founding population (bottleneck) *b* = 200. Since neutralization during replication takes the place of mucosal sIgA neutralization, *b* should be understood as comparable to the sIgA bottleneck size *v* in a model with sIgA neutralization. (**A–C**) Top panels: absolute abundances of target cells (green), old antigenic variant virions (blue), and new antigenic variant virions (red). Bottom panels: frequencies of virion types. Black dotted line is an analytical prediction for the new antigenic variant frequency given the time of first appearance. Black dashed line is the threshold for NGS detection. (**D**) Probabilities of no infection, *de novo* new antigenic variant infection, inoculated new antigenic variant infection, and old antigenic variant infection per 10^5^ inoculations of an immune host by a naive host. Circles are frequencies from simulation runs (10^6^ runs for bottlenecks 1–10, 10^5^ runs for bottlenecks 50–200). Plus-signs are analytical model predictions for probabilities (see Appendix section 3.6), with *f*_mt_ set equal to the average from donor host stochastic simulations for the given bottleneck. Parameters as Table 1 unless otherwise stated.

If there is antibody neutralization throughout virus exponential growth and it is not sufficiently strong to control the infection, this facilitates the establishment of new antigenic variants: variants can be generated *de novo* and then selected to detectable and easily transmissible frequencies (Fig. 1C,D,F, sensitivity analysis in Fig. A3). We term selection on a replicating within-host virus population “replication selection”. Virus phenotypes that directly affect fitness independent of immune system interactions are likely to be subject to replication selection.

Adding a realistic delay to antibody production of two days post-infection (Lam & Baumgarth, 2019) curtails antigenic replication selection. There is no focused antibody-mediated response during the virus exponential growth phase, and so the infection is dominated by old antigenic variant virus (Fig. 1C,D,G, sensitivity analysis in Fig. A3). Antigenic variant viruses begin to be selected to higher frequencies late in infection, once a memory response produces high concentrations of cross-reactive antibodies. But by the time this happens in typical infections, both new antigenic variant and old antigenic variant populations have peaked and begun to decline due to innate immunity and depletion of infectible cells, so new antigenic variants remain too rare to be detectable with NGS (Fig. 1G).

We find that replication selection of antigenic novelty to detectable levels becomes likely only if infections are prolonged, and virus antigenic diversity and antibody selection pressure therefore coincide (Fig. 1G, see also Fig. 7). This can explain existing observations: within-host adaptive antigenic evolution can be seen in prolonged infections of immune-compromised hosts (Xue et al., 2017), and prolonged influenza infections show large within-host effective population sizes (Lumby et al., 2020).

### Neutralization of virions at the point of transmission provides host protection and population-level selection without rapid within-host adaptation

Adding antibody neutralization of virions (e.g. by mucosal sIgA) at the point of inoculation to our model produces realistic levels of protection against reinfection, and when reinfections do occur, they are overwhelmingly old antigenic variant reinfections. New antigenic variants that arise during these infections remain undetectably rare, reproducing observations from natural human infections (Fig. 1A–D, G–H, Fig. 3B–C)(Debbink et al., 2017; Dinis et al., 2016; Leonard et al., 2016; McCrone et al., 2018). The combination of mucosal sIgA protection and a realistically timed antibody recall response explains how there can exist immune protection against reinfection—and thus a population level selective advantage for new antigenic variants—without observable within-host antigenic selection in typical infections of experienced hosts.

**Fig. 3.**
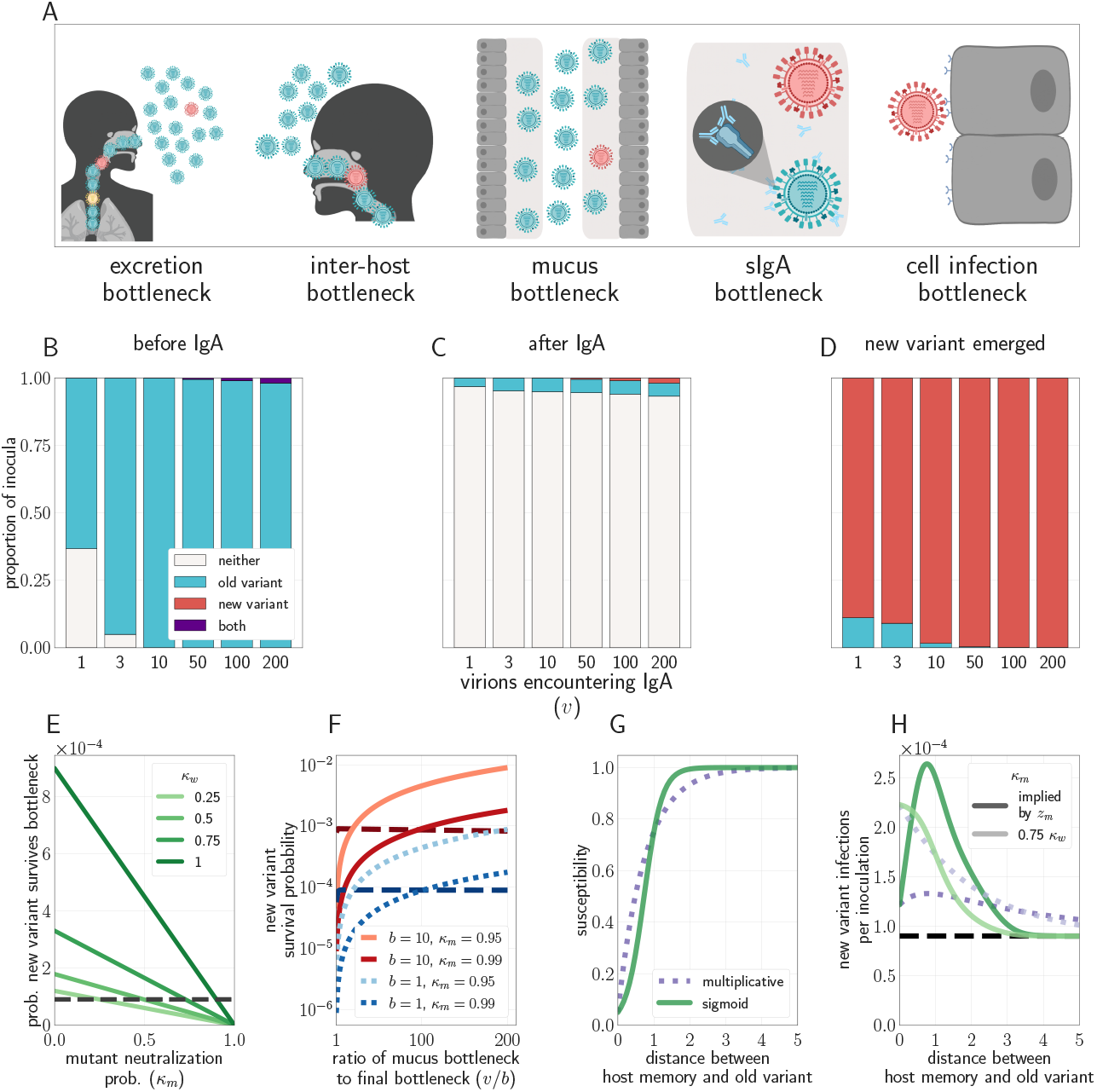
Selection for antigenic variants at the point of transmission (inoculation selection). (**A**) Schematic of bottlenecks faced by both old antigenic variant (blue) and new antigenic variant (other colors) virions at the point of virus transmission. Key parameters for inoculation selection are the mucus bottleneck *v*—the mean number of virions that encounter sIgA—and the cell infection bottleneck size *b*. (**B–D**) Effect of sIgA selection at the point of inoculation with *b* =1. (**B, C**) Analytical model distribution of virions inoculated into an immune host immediately before (**B**) and after (**C**) mucosal neutralization / the sIgA bottleneck. *f_mt_* set to mean of stochastic simulations. (**D**) Distribution of founding virion populations (after the cell infection bottleneck) among individuals who developed detectable new antigenic variant infections in stochastic simulations. (**E–F**) Analytical model probability that a variant survives the final bottleneck. Dashed horizontal lines indicate probability in naive hosts. *f*_mt_ = 9 × 10^−5^ (the approximate mean in stochastic simulations for *b* = 1). (**E**) Variant probability of surviving all bottlenecks, as a function of old antigenic variant neutralization probability *κ_w_* and new antigenic variant mucosal neutralization probability *κ_m_*. (**F**) New antigenic variant survival probability as a function of the ratio of *v* to *b*. (**G, H**) Effect of host susceptibility model on the appearance of antigenic novelty. Per inoculation rates of new variants surviving the bottleneck (**H**) depend on host immune status and on the relationship between virus antigenic phenotype and host susceptibility (1 – *z_w_*) (**G**). Plotted with *f*_mt_ = 9 × 10^−5^, and 25% susceptibility to (75% protection against, *z*_1_ = 0.75) a variant one antigenic cluster away from host memory. Unless noted, parameters for all plots as in Table 1.

### Tight bottlenecks lead to loss of generated diversity and mean new variants reach consensus through founder effects

Regardless of host immune status, an antigenic variant that has been generated *de novo* within a host must survive a series of population bottlenecks if it is to infect other individuals. To found a new infection, virions must be expelled from a currently infected host (excretion bottleneck), must enter another host (inter-host bottleneck), must escape mucus on the surface of the airway epithelium (mucus bottleneck), must avoid neutralization by sIgA antibodies on mucosal surfaces (sIgA bottleneck), and must infect a cell early enough to form a detectable fraction of the resultant infection (cell infection bottleneck) (Fig. 3A). The sum of all of these effects is the net bottleneck and typically results in infections being initiated by a single genomic variant (Ghafari et al., 2020; McCrone et al., 2018; Xue & Bloom, 2019). That said, bottlenecks resulting from direct contact transmission may be substantially wider than those associated with respiratory droplet or aerosol transmission (Varble et al., 2014) and more human studies are required to quantify these differences.

We find that because antigenic variants appear at very low within-host frequencies when generated *de novo* and undergo minimal or no replication selection, new antigenic variants most commonly reach detectable levels within hosts through founder effects at the point of inter-host transmission: a low-frequency antigenic variant generated in one host survives the net bottleneck to found the infection of a second host (Fig. 3D).

Given that influenza bottlenecks are thought to be on the order of a single virion (McCrone et al., 2018), any replication-competent mutant that founds an infection should occur at NGS-detectable levels, and likely at consensus. But for the same reason, these founder effects are rare events. These founder effects could be a purely neutral sampling process. It is likely quite close to neutral in a truly naive host who does not possess well-matched antibodies to the infecting old variant: all inoculated virions, regardless of antigenic phenotype, have an equal chance of becoming part of the new infection’s founding population. If there are *v* virions that compete to found the infection and *b* =1, then each virion founds the infection with probability 1/*v*.

In our model, new antigenic variants survive the transmission bottleneck upon inoculation into a naive host with a probability approximately equal to the donor host variant frequency *f*_mt_ times the bottleneck size *b* (see Methods). New antigenic variant infections of naive hosts should therefore occur on the order of 1 in 10^5^ or 1 in 10^4^ of such infections given biologically plausible parameters (Fig. 3E–H).

But if different virions have different chances of being neutralized at the point of transmission, the founding process may be selective. Among the virions that encounter antibodies, those that are less likely to be neutralized have a higher than average chance of undergoing stochastic promotion to consensus while those that are more likely to be neutralized have a lower than average chance. A new antigenic variant is then disproportionately likely to survive the net bottleneck (Figs. 1, 3, Appendix section 4.2). We term this potential selection on inoculated diversity “inoculation selection”. Neutralization at the point of transmission thus not only gives new antigenic variant infections their transmission advantage (population-level selection) but may also increase the rate at which these new antigenic variant infections arise (inoculation selection).

There is some suggestive evidence of differential survival of particular (not necessarily antigenic) influenza genetic variants at the point of transmission from experiments in ferrets (Moncla et al., 2016; Wilker et al., 2013). But as Lumby and colleagues (2018) point out in a reanalysis of those experiments, it difficult empirically to distinguish selection that occurs at the point of transmission from selection that occurs during replication in the donor host because of the challenges associated with sampling the small virus populations present at the earliest stages of infection. Here we define inoculation selection as selection on the bottlenecked virus population that establishes infection in the recipient host before any virus replication has taken place in that host.

### Inoculation selection depends on degree of founding competition and degree of immune escape

The strength of inoculation selection depends on the ratio of the number of virions that compete to found an infection in the absence of well-matched sIgA antibodies (the mucus bottleneck size *v*) to the number of virions that actually found an infection (the final cell infection bottleneck size *b*). The larger this *v*/*b* ratio is, the more inoculation selection in experienced hosts facilitates the survival of new antigenic variants (Fig. 3F, Fig. A1).

When new antigenic variant immune escape is incomplete due to partial cross-reactivity with previous antigenic variants, increased antibody neutralization is a double-edged sword for new antigenic variant virions. Competition to found the infection from old antigenic variant virions is reduced, but the new antigenic variant is itself at greater risk of being neutralized. The impact of inoculation selection therefore depends on the degree of similarity between previously encountered viruses and the new antigenic variant. An experienced recipient host could facilitate the survival of large-effect antigenic variants (like those seen at antigenic cluster transitions (Smith et al., 2004)) while retarding the survival of variants that provide less substantial immune escape (Fig. 3E, Fig. A1).

Inoculation selection is limited by the low frequency *f*_mt_ of new antigenic variants in transmitting donor hosts (due to weak replication selection), the potentially small mucus bottleneck size *v*, and the fact that some hosts previously infected with the same or similar antigenic variants might not possess well-matched antibodies due to original antigenic sin (Davenport & Hennessy, 1956), antigenic seniority (Lessler et al., 2012), immune backboosting (Fonville et al., 2014), or other sources of individual-specific variation in antibody production (Lee et al., 2019). These factors combined make selection and onward transmission of new variants rare.

### Immune hosts can facilitate the appearance of new variants without producing rapid diversification

Onward transmission of new variants can be facilitated by natural selection—replication selection, inoculation selection or both. The degree of facilitation depends principally on four quantities: 1. *δτ*, the product of the replication fitness difference *δ* = *k*(*c_w_* – *c_m_*) and the time under replication selection *τ*. This determines the degree to which the new variant is promoted by replication selection prior to transmission (increasing *f*_mt_). 2. *κ_w_*, the neutralization probability for the old variant, which must be large enough to reduce competition for the final bottleneck. 3. *v/b*, the ratio of the sIgA bottleneck size *v* to the cell infection (final) bottleneck size *b*. This determines the degree of competition to found the infection, and thus sets maximum potential strength of inoculation selection when *κ_w_* is large: a 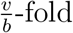 improvement over drift for small *b*. 4. 1 – *κ_m_*, how likely the new variant is to avoid neutralization at the point of transmission; this scales down the maximum replication selective strength set by *v/b* (to zero if the new variant is neutralized with certainty; *κ_m_* = 1). When *δτ* is small and *κ_w_*, *v/b*, and 1 – *κ_m_* are large, new variant survival is most facilitated by inoculation selection. When the opposite is true, replication selection is most important. And there are parameter regimes in which both replication and and inoculation selection provide a substantial improvement over drift (Fig. 4, see Appendix section 4.6 for mathematical intuition for these results).

**Fig. 4.**
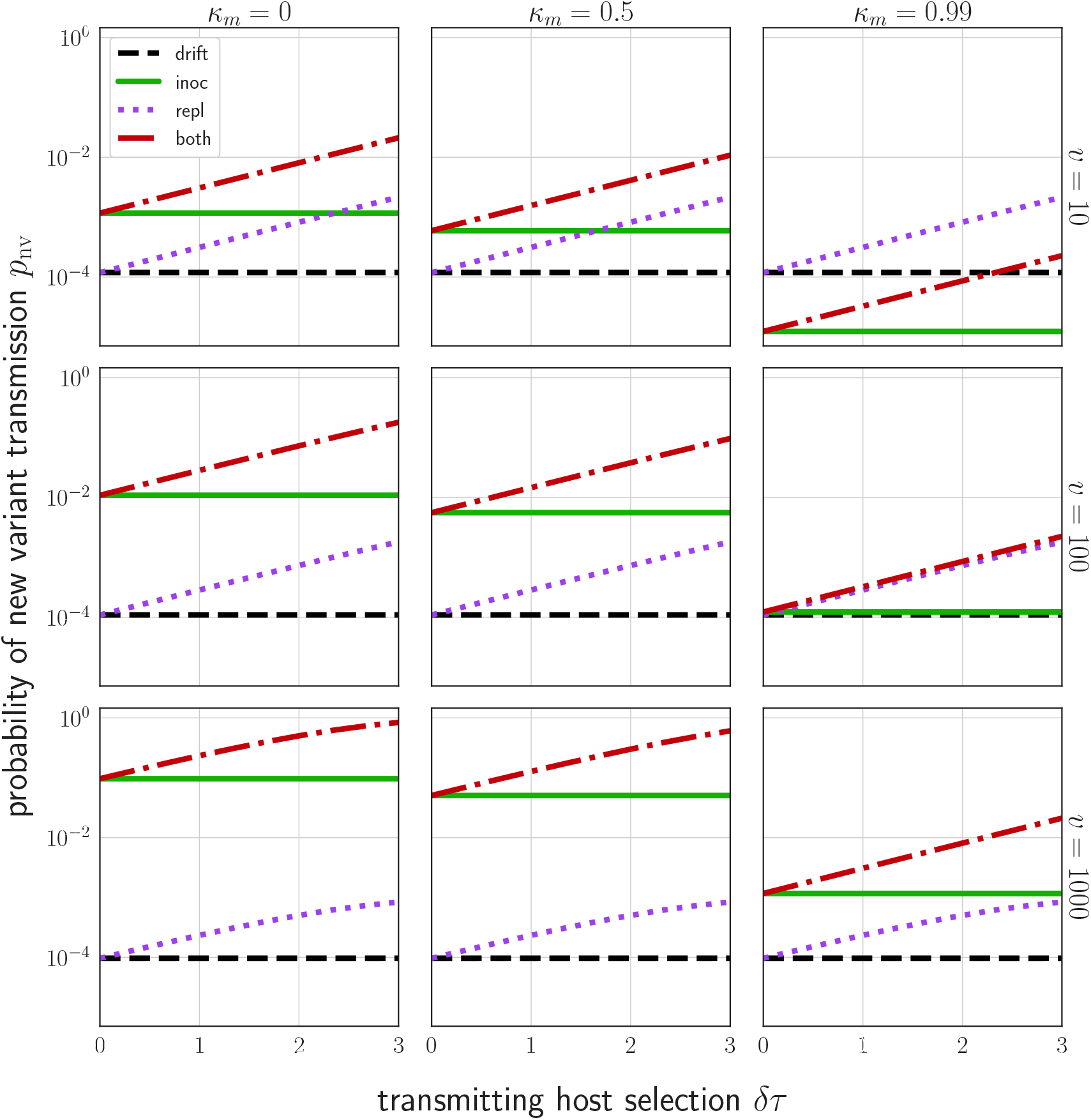
Probability *p*_nv_ of a new variant surviving the transmission bottleneck as a function of donor host replication selection and recipient host inoculation selection. Calculated according to equation 43, and plotted as a function of degree of replication selection in the donor host *δτ*, the product of the selection strength *δ* and the time duration *τ* = max 0, *t_t_* – *t_M_* between the onset of the antibody response at *t_M_* and the transmission event at *t_t_*. Black dashed line: neutral (drift) expectation, where *δ* = 0 in the donor host and the recipient host does not neutralize either the old or the new variant at the point of transmission (*κ_w_* = *κ_m_* = 0). Purple dotted line: replication selection only: *δτ* as given in the donor host, but a naive recipient host. Green solid line: inoculation selection only: *δ* = 0 in the donor host, but a recipient host with well-matched antibodies to the old variant (*κ_w_* = 1), with varying degrees of immune escape (*κ_m_* as given in the columns). Red dot-dashed line: combination of both replication selection in the donor host as before and inoculation selection in the recipient host as before. Plotted with a final bottleneck of *b* =1 and an sIgA bottleneck *v* as given in the rows. Parameters as in Table 1 unless otherwise noted.

At realistic parameter values and assuming all individuals develop well-matched antibodies to previously encountered antigenic variants, only ~1 to 2 in 10^4^ inoculations of an experienced host results in a new antigenic variant surviving the bottleneck (Fig. 3E,H, Fig. 4). This rate is likely to be an overestimate due to the factors mentioned above. Moreover, it is only about 2- to 3-fold higher than the rate of bottleneck survival in naive hosts, where new antigenic variant infections should occasionally occur via neutral stochastic founder effects. In short, even in the presence of experienced hosts, antigenic selection is inefficient and most generated antigenic diversity is lost at the point of transmission. Because of these inefficiencies, new antigenic variants can be generated in every infected host without producing explosive antigenic diversification at the population level.

### Inoculation selection produces realistically noisy between-host evolution

To investigate the between-host consequences of adaptation given weak replication selection, tight bottlenecks, and possible inoculation selection, we simulated transmission chains according to our within-host model, allowing the virus to evolve in a 1-dimensional antigenic space (Bedford et al., 2012; Smith et al., 2004) until a generated antigenic mutant becomes the majority within-host variant. When all hosts in a model transmission chain are naive, antigenic evolution is non-directional and recapitulates the distribution of within-host mutations (Fig. 5A). When antigenic selection is constant throughout infection and even a moderate fraction of hosts are experienced, antigenic evolution is unrealistically strong: the virus evolves directly away from existing immunity and large-effect antigenic changes are observed frequently (Fig. 5B,C). When the model incorporates both mucosal antibodies and realistically timed recall responses, major antigenic variants appear only rarely and the overall distribution of emerged variants better mimics empirical observations (Fig. 5D); most notably, the phenomenon of quasi-neutral diversification within an antigenic cluster seen in Figures 1 and 2 of Smith et al., 2004. A simple analytical model (see Methods) in which generated antigenic mutants fix according to their replication and inoculation selective advantages also displays this behavior (Fig. 5B–H).

**Fig. 5.**
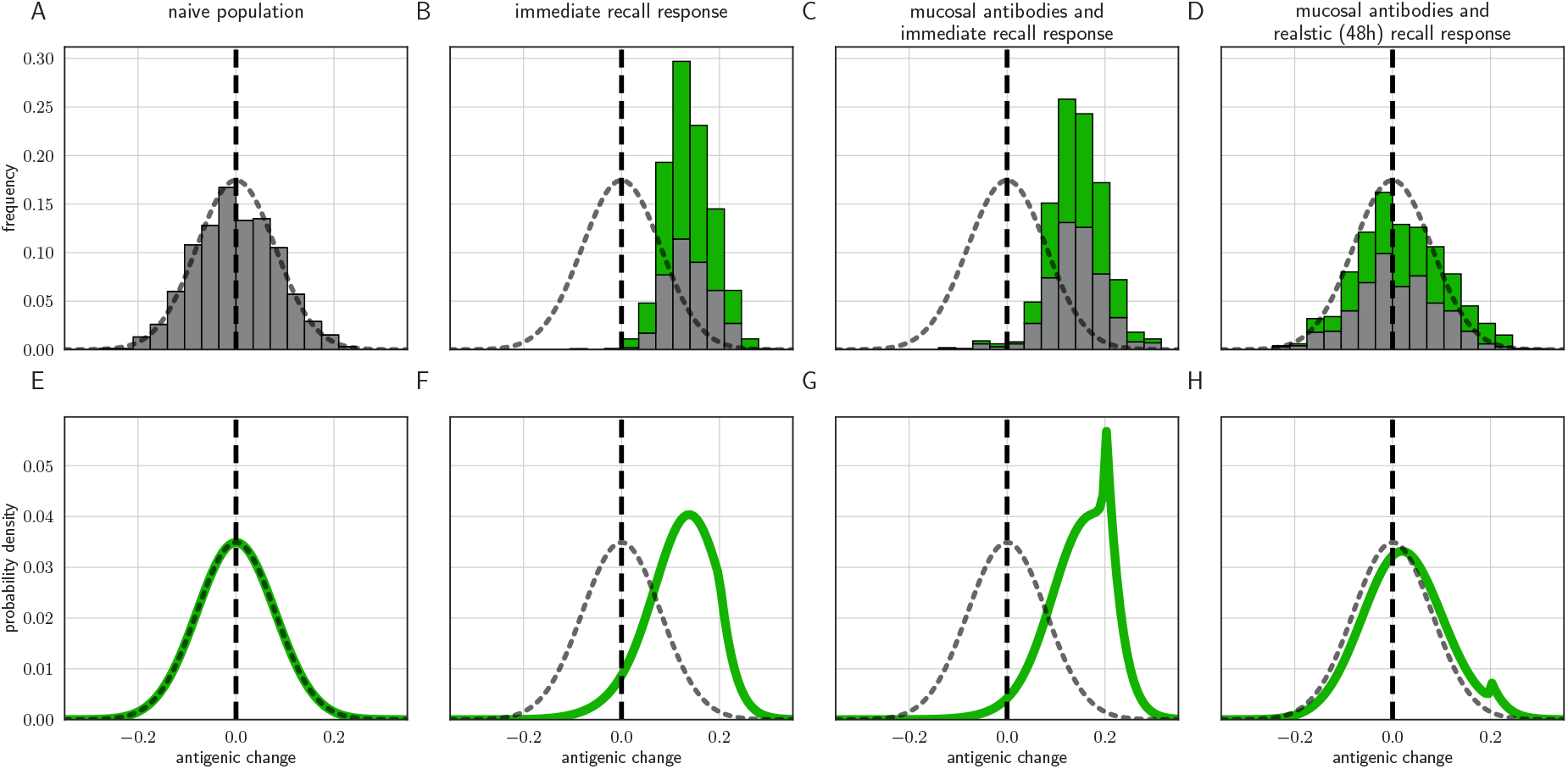
Distribution of mutant effects given replication and inoculation selection. Distribution of antigenic changes along 1000 simulated transmission chains (**A–D**) and from an analytical model (**E–H**). In (**A,E**) all naive hosts, in other panels a mix of naive hosts and experienced hosts. Antigenic phenotypes are numbers in a 1-dimensional antigenic space and govern both sIgA cross immunity *σ* and replication cross immunity *c*. A distance of ≥ 1 corresponds to no cross immunity between phenotypes and a distance of 0 to complete cross immunity. Gray line gives the shape of Gaussian within-host mutation kernel. Histograms show frequency distribution of observed antigenic change events and indicate whether the change took place in a naive (grey) or experienced (green) host. In (**B–D**) distribution of host immune histories is 20% of individuals previously exposed to phenotype −0.8, 20% to phenotype −0.5, 20% to phenotype 0 and the remaining 40% of hosts naive. In (**E**), a naive hosts inoculate naive hosts. In (**F–H**) a host with history −0.8 inoculate hosts with history −0.8. Initial variant has phenotype 0 in all sub-panels. Model parameters as in Table 1, except *k* = 25. Spikes in densities occur at 0.2 as this is the point of full escape in a host previously exposed to phenotype −0.8.

In particular, we note that whereas an immediate recall response would predict strong near-constant directed evolution of virus antigenic phenotypes away from existing immunity (Fig. 5B,C), a realistically-timed recall response predicts that small-effect, drift-like, antigenic substitutions will be observed. Even substitutions that move a virus “backward” in antigenic space and that should be more readily neutralized by existing antibodies than the ancestral variant should be observed thanks to the large role of stochasticity at the point of transmission. That said, there is a slight bias favoring forward substitutions, especially those of sufficiently large effect (Fig. 5D) to create a substantial selective advantage over the ancestral variant. Coupled with the plausible assumption that large effect substitutions are rarer than small effect substitutions (here captured qualitatively by the Gaussian mutation kernel), this predicts the pattern of quasi-neutral diversification within antigenic clusters followed by rarer directional “jumps” in phenotype. Exact rates of antigenic evolution will depend upon how these emergence processes intersect with population-level epidemic dynamics and competition among variants.

### Epidemic dynamics can alter rates of inoculation selection

We used an epidemic-level model to study the consequences of individual-level inoculation selection for population-level antigenic selection. If inoculation selection is efficient, an intermediate initial fraction of immune hosts maximizes the probability that a new antigenic variant infection is created during an epidemic (Fig. 6.) This is due to a trade-off between the frequency of naive or weakly immune “generator” hosts who can propagate the epidemic and produce new antigenic variants through *de novo* mutation, and the frequency of strongly immune “selector” hosts who, if inoculated, are unlikely to be infected, but can facilitate the survival of these new antigenic variants at the point of transmission. As selector host frequency increases, epidemics become rarer and smaller, eventually decreasing opportunities for evolution, but moderate numbers of efficient selectors can substantially increase the rate at which new antigenic variants reach within-host consensus (Fig. 6B,C).

**Fig. 6.**
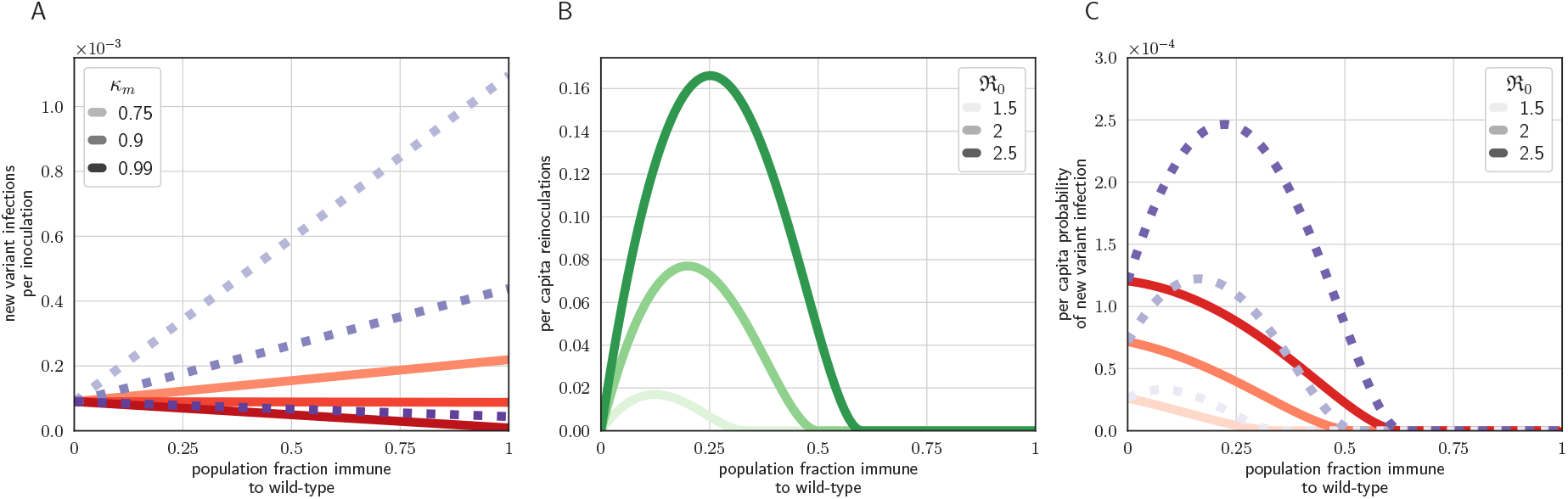
Population level antigenic dynamics resulting from inoculation selection. Analytical model results (see Methods) for population-level inoculation selection, using parameters in Table 1 and *f*_mt_ = 9 × 10^−5^ unless otherwise stated. (**A**) Probability per inoculation of a new antigenic variant founding an infection, as a function of fraction of hosts previously exposed to the infecting old antigenic variant virus, mucus bottleneck size *v* and sIgA cross immunity *σ* = *κ_m_*/*κ_w_*. Red solid lines: *v* = 10. Purple dotted lines: *v* = 50. (**B**) Expected per capita reinoculations of previously exposed hosts during an epidemic, given the fraction of previously exposed hosts in the population, if all hosts that were previously exposed to the circulating old antigenic variant virus are fully immune to that variant, for varying population level basic reproduction number 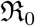. (**C**) Probability per individual host that a new antigenic variant founds an infection in that host during an epidemic, as a function of the fraction of hosts previously exposed to the old antigenic variant. Other hosts naive. *σ* = 0.75. Red solid lines: *v* = 10; purple dotted lines: *v* = 50 (as in **A**).

## Discussion

Any explanation of influenza virus antigenic evolution—and why it is not even faster—must explain why population level antigenic selection is strong, as evidenced by the typically rapid sequential population-level replacement of old antigenic variants upon the emergence of a new major antigenic variant, but within-host antigenic evolution is rarely observed.

We hypothesized that antibodies present in the repiratory tract mucosa at the time of virus exposure can effectively block transmission, but have only a small effect on viral replication once cells become productively infected. Antigenic selection after successful infection therefore begins with the mounting of a recall response 48–72 hours post infection. In this case, selection pressure can be strong at the point of transmission, but subsequently weak until after the period of virus exponential growth. This mechanistic paradigm reconciles strong but not perfect sterilizing homotypic immunity with rare observations of new antigenic variants in successfully reinfected experienced hosts.

### Alternative explanations for rare new antigenic variants

We consider several possible explanations for the observed phenomenon that new antigenic variants are rare within experienced infected hosts and at the population level prior to cluster transitions. But among these candidate hypotheses, only the mechanism of small inocula, transmission-blocking mucosal antibodies, but a slow-to-mount recall adaptive immune response can explain all the aforementioned empirical observations simultaneously.

Alternative possibilities include strong immune protection through antibody neutralization early during early viral replication (an immediate recall response), heterogeneous neutralization rates during early viral replication, new antigenic variants that are deleterious in the absence of antibody selection, and the need for new variants to emerge against a favorable genetic background.

As shown above, protection through antibody neutralization early in replication can result in rare within-host observation of new antigenic variants, but it contradicts understanding of antibody kinetics and makes other empirical predictions that are unrealistic. It implies that homotypic challenge has a binary outcome: either it results in an undetectable infection that is rapidly cleared or it results in a visible infection dominated by an escape mutant (Fig. 2). This is how influenza viruses have been hypothesized to behave in previous models of immune escape (Luo et al., 2012). But empirical work (Debbink et al., 2017; Javaid et al., 2020; McCrone et al., 2018) and human challenge studies (Clements et al., 1986; Memoli et al., 2019) have shown that detectable reinfection of experienced hosts can occur without observable immune escape. Another empirical prediction of such a model is that intermediately immune hosts should be efficient selectors, since they neutralize the old variant virus poorly enough to allow it to grow, but strongly enough to impose antigenic selection upon that growing population (Volkov et al., 2010). While such intermediately immune hosts should be present from the beginning of a new antigenic cluster’s circulation (Fonville et al., 2014), new variants are rarely observed until a new variant has circulated for multiple years (Smith et al., 2004).

Antibody neutralization during early replication could avoid binary outcomes if individual hosts are sufficiently heterogeneous in the strength of their neutralizing response, so that some individuals clear the infection rapidly while others barely exert antibody selection upon it. But while heterogeneity in immunity exists (Lee et al., 2019), this explanation requires extreme, bimodal hetegoneity to avoid the intermediately immunity regime in which replication selection is efficient (Volkov et al., 2010) and again requires an unrealistically early antibody response.

New antigenic variants could be replication-competent, but weakly deleterious within-host in the absence of immune selection and/or compensatory substitutions. Two studies (Gog, 2008; Kucharski et al., 2015) have invoked this hypothesis to explain population-level antigenic dynamics. However, a realistically strong antibody response during virus replication could still promote new variants during infections of experienced hosts, even in the absence of compensatory mutations (see Appendix section 7.4 for a model analysis). So a hypothesis to explain the relative weakness of selection during replication is still required, especially for weakly deleterious mutants that offer substantial immune escape. That said, under our hypothesis of founder effects and possible inoculation selection, weak new variant deleteriousness could further limit the rate of antigenic evolution by reducing the probability that new variants are inoculated into hosts (see Appendix section 7.4).

Finally, it has been hypothesized that antigenic mutants can only proliferate at the population level if they arise against a favorable genetic background (Koelle & Rasmussen, 2015)—in the absence of deleterious substitutions elsewhere in the genome. But antigenic cluster transitions are frequently polyphyletic (see Appendix section 6) where a new variant emerges quasi-simultaneously in multiple virus lineages. Since these lineages should have different genetic backgrounds, this suggests that favorable backgrounds are readily available, and that emergence is limited instead by the presence or absence of selection pressure.

We discuss these alternative explanations in more depth in the Appendix (sections 7 and 7.4).

### Relationship to prior influenza virus transmission bottleneck literature

Previous literature on influenza virus transmission mentions a “selective bottleneck” (Leonard et al., 2016; Moncla et al., 2016; Wilker et al., 2013), but these studies do not refer to antigenic inoculation selection. Rather, a “selective bottleneck” has been typically used to mean simply a tight neutral bottleneck (that leads to stochastic loss of diversity) or to non-antigenic factors that lead to preferential transmission of certain variants (Moncla et al., 2016). An important exception is (Lumby et al., 2018). The authors studied ferret transmission experiments and partitioned selection for adaptive mutants (not necessarily antigenic) into selection for transmissibility (acting at a potentially tight bottleneck) and selection during exponential growth. Subsequently, several studies have hypothesized that influenza virus antigenic selection might be weak in short-lived infections of individual experienced hosts and might occur at the point of transmission (Han et al., 2019; Lumby et al., 2020; Petrova & Russell, 2018).

To our knowledge, however, ours is the first study to undertake a mechanistic, model-based comparison between the role of selection at the point of transmission versus selection during replication in the promotion of antigenic mutants, to show that immunologically plausible mechanisms could make the former more salient than the latter, and to connect that finding to the rarity of observable new antigenic variants in homotypically reinfected human hosts. We discuss the relationship between this paradigm and those put forward in previous modeling studies at further length in section of the Appendix.

### Limitations and remaining uncertainties

The study presented here nonetheless has important limitations that suggest opportunities for future investigation. This is a modeling study, and a mainly theoretical one. This is out of necessity. Quality experiments of the kind that are necessary to observe selection at the point of transmission directly and measure its strength have not been published, and we were unable to find any experimental measurements of within-host competition between known antigenic variants. For additional discussion of unmodeled biological realities and mechanistic uncertainties, see sections 2 and 5 of the Appendix.

#### Within-host model

Our within-host model is a simple target cell-limited model, but the decrease in infectible cells as the infection proceeds can be qualitatively interpreted as any and all antigenicity-agnostic limiting factors that come into play as the virion and infected cell populations grow. This could include the action of innate immunity, which acts in part by killing infected cells and by rendering healthy cells difficult to infect via inflammation (see immunological review in 2 of the Appendix). The key mechanistic role played by the target cells in our model is to introduce a non-antigenic limiting factor on infections. This prevents an infection from repeatedly evolving out from under successive well-matched antibody responses, as occurs in HIV and in influenza patients with compromised immune systems (Xue et al., 2017). These factors limit the virus regardless of whether the infection remains confined to the upper respiratory tract or also infect the lower respiratory tract, thereby gaining access to additional target cells (Koel et al., 2019). The key is that non-antigenic factors prevent persistent large virus populations.

Similarly, the antibody response we introduce at 48 hours is qualitative—it is modeled simply as an increase in the virion decay rate for antibody matching virions. This response could represent IgA targeted at HA, but other antigenicity-specific modes of virus control could also be subsumed under the increased virion decay rate, for instance IgG antibodies, antibody dependent cellular cytotoxicity (ADCC), antibodies against the neuraminidase protein, or others. The key point we establish is that none of these mechanisms efficiently replication select because they all emerge once non-antigenic limiting factors have come into play. They speed clearance, but should not substantially alter virus evolution. A corollary to this point is that there are many mechanisms—and interventions—that could reduce the severity of influenza infections without substantially speeding up antigenic evolution, including universal vaccines.

#### Point of transmission

A key proposal of our study is that population-level antigenic selection and homotypic protection are mediated by antibody neutralization (likely sIgA) at the point of transmission. Currently, empirical evidence for antibody protection at the point of transmission is mostly indirect. Most of this evidence comes from human and animal challenge studies (Clements et al., 1986; Le Sage et al., 2020; Memoli et al., 2019). In these studies, individuals who are challenged with the same antigenic variant sometimes display apparent sterilizing immunity, but other times develop detectable infections. The study by (Memoli et al., 2019) is notable for having used very large inocula—10^6^ or 10^7^ TCID_50_. Despite these high doses, two of the challenge subjects had neither detectable virus nor seroconversion. Similarly, ferret experiments (Le Sage et al., 2020) have shown that many ferrets develop sufficiently sterilizing immunity to prevent the virus from ever being detected, while some ferrets showed briefly detectable infections that were then cleared. While we cannot rule out a powerful immediate cellular response that was differentially evaded in the various subjects, we believe that our model, coupled with existing understanding of the timing of cellular responses and the speed of influenza virus replication, provides a more parsimonious explanation.

Another limitation of our study is that, while we put forward mucosal sIgA as a biologically documented potential mechanism of immune protection at the point of inoculation that would not lead to strong selection during early viral replication, no modeling study can establish such a mechanism without empirical investigation. Our study reveals that such an empirical investigation would be of substantial scientific value.

We model neutralization at the point of transmission as a binomial process. Each virion is independently neutralized with a probability *κ_i_* that depends on its antigenic phenotype and the host’s immune history. As we discuss in Appendix section 5.2, this assumption of independence may be violated in practice. Careful experiments are required to develop a more realistic model of neutralization at the point of transmission.

Moreover, individual variation in immune system properties and complex effects of host immune history (Fonville et al., 2014; Lee et al., 2019) mean that even a pair of hosts who have both been previously exposed to the currently circulating variant may exert different selection pressures at the point of transmission. Modeling neutralization as a series of independent events that depend only on host history and virus phenotype is a baseline: it allows us to establish that antigenic selection at the point of transmission is possible and show what its consequences might be. But a more realistic model will be required to predict the selective pressures imposed by real hosts with real immune histories on real virions.

Finally, while we believe neutralization at the point of transmission is a crucial mechanism of protection against detectable reinfection for influenza, this may not be true for all RNA viruses. Some, such as measles and varicella, have long incubation periods even in naive hosts and induce reliable, long-lasting immune protection against detectable reinfection. This may be because they replicate slowly enough that they cannot “outrun” the adaptive response as influenza can. Neither shows influenza virus-like patterns of clocklike immune escape, suggesting that (1) escape mutants may be less available and (2) that adaptive response acts on a small population and is forceful.

#### Parameter uncertainties

We parameterized our models based on estimates from previous studies, but there are not good estimates for several important quantities. There are no high quality estimates of the rate of antibody-mediated neutralization in the presence of a homotypic antibody response or of how much this rate is reduced by particular antigenic substitutions, and there may be substantial inter-individual variation (Lee et al., 2019). Within-host timeseries data from antigenically heterogeneous infections are needed to estimate these quantities. Similarly, there are no good empirical data on the size of the bottlenecks that precede or follow the sIgA bottleneck (Fig. 3A of the main text). Better estimates of these bottlenecks, and of the probabilities of neutralization for individual virions encountering mucosal sIgA antibodies, would give more certainty about the strength of inoculation selection relative to neutral founder effects.

We also do not have a clear sense of exactly how neutralization probability in the respiratory tract mucosa (parameter *κ* in our model) and neutralization rate during replication (parameter *k* in our model) relate. We expect them to be positively related, but the exact strength and shape of this relationship is unknown. Knowing whether major antigenic changes reduce both equally or reduce one more than the other could help us better quantify the potential strength of inoculation selection and replication selection. In short, better mechanistic understanding of mucosal antibody neutralization could be extremely valuable for understanding and potentially predicting influenza virus evolution.

#### Scaling up to the population level

How readily a particular individual host or host population helps new antigenic variants reach within-host consensus depends upon several unknown quantities: 1. how host susceptibility changes with extent of antigenic dissimilarity, 2. the ratio of virions that encounter sIgA to virions that found the infection (*v/b*), 3. the probability that a single new antigenic variant virion inoculated alongside old antigenic variant virions evades neutralization 1 – *κ_m_* (Fig. 3E–H), 4. the duration *τ* from the onset of the antibody response to the time of transmission. Better empirical estimates of these quantities could shed light on how the distribution of host immunity shapes antigenic evolution. However, over a range of biologically plausible parameter values, our model contradicts the existing hypothesis that antigenic novelty appears when moderately immune hosts fail to block transmission and then select upon a growing virus population (Grenfell et al., 2004; Volkov et al., 2010). For influenza viruses, hosts whose mucosal immunity that regularly blocks old antigenic variant transmission may be crucial. Mucosal immunity not only produces a population-level advantage for new variants but may also play a role in their within-host emergence (Fig. 3E,H).

### Implications

Our study has a number of implications for the study and control of influenza viruses.

#### Importance of host heterogeneity

Experienced hosts are undoubtedly heterogeneous in their immunity to a given influenza variant (Lee et al., 2019), so the overall population average protection against homotypic reinfection with variant *i*, *z_i_*, is in fact an average over experienced hosts. Our model implies that the degree of neutralization difference between ancestral variant virions and new antigenic variant virions at the point of transmission strongly affects the probability of inoculation selection. Hosts with more focused immune responses—highly specific antibodies that neutralize old antigenic variant virions well and new antigenic variant virions poorly—could be especially good inoculation selectors and important sources of population-level antigenic selection. Hosts who develop less specific memory responses, such as very young children (Neuzil et al., 2006), could be less important. Similarly, immune-compromised hosts are excellent replication selectors (Lumby et al., 2020; Xue et al., 2017), and so their role in the generation of antigenic novelty and their impact on overall population level diversification rates both deserve further study.

#### Small-population-like evolution

Prior modeling has suggested that despite repeated tight bottlenecks at the point of transmission, evolution of influenza viruses should resemble evolution in idealized large populations (Sigal et al., 2018). In large populations, advantageous variants with small selective advantages should gradually fix and weakly deleterious variants should be purged. In these studies, diversity is rapidly generated and fit variants are selected to frequencies at which they are likely to pass through even a tight bottleneck. This is likely true of the phenotypes modeled in those studies, which include receptor binding avidity and virus budding time (Sigal et al., 2018). These phenotypes affect virus fitness throughout the timecourse of infection, so they can be efficiently replication-selected (where selection is manifest in the direct competition for infecting other cells rather than indirect competition by escaping antibodies). Indeed, next generation sequencing studies have found observable adaptative evolution of non-antigenic phenotypes in individual humans infected with avian H5N1 viruses (Welkers et al., 2019).

Seasonal influenza antigenic evolution does not resemble idealized large population evolution. Within an antigenic cluster, influenza viruses acquire substitutions that change the antigenic phenotype by small amounts. Given the large influenza virus populations within individual hosts, we might expect quasi-continuous directional pattern of evolution away from prior population immunity. Indeed, between clusters, evolution is strongly directional: only “forward” cluster transitions are observed. These are jumps—large antigenic changes. But incremental within-cluster evolution is *not* directional: the virus often evolves “backward” or “sideways” in antigenic space towards previously circulated variants (see Figs. 1 and 2 of (Smith et al., 2004)).

This noisy jump pattern is easy to explain in light of the weakness of replication selection and the importance of antigenic founder effects. Selection acts on the small sub-sample of donor-host diversity that passes through the excretion, inter-host, and mucus bottlenecks to encounter the sIgA bottleneck. Evolution via inoculation selection is therefore slower and more affected by stochasticity than evolution via replication selection (Fig. 3E–F). It resembles evolution in small populations—weakly adaptive and weakly deleterious substitutions become nearly neutral (Kimura, 1968; Ohta, 1992). If influenza virus evolution were not nearly neutral for small-effect substitutions at the within-host scale, it would be surprising to observe “backward” antigenic changes and noisy evolution at higher scales (Fig. 5A–D). In fact, there may be analogous “neutralizing” population dynamics at higher scales as well, and those may also be necessary for explaining population level noisiness. But whatever happens at higher scales, within-host replication selection would create a strongly directional bias in what population-level diversity is introduced (Fig. 5A–D), and introduces many small forward antigenic changes. Inoculation selection does not necessarily do this.

#### Population level neutralizing dynamics

Local influenza virus lineages rarely persist between epidemics (Bedford et al., 2015; Russell et al., 2008), and so new antigenic variants must establish chains of infections in other geographic locations in order to survive. new antigenic variant chains are most often founded when inoculations are common—that is, when existing variants are causing epidemics. Epidemics result in high levels of local competition between extant and new antigenic variant viruses for susceptible hosts (Hartfield & Alizon, 2015) as well as metapopulation-scale competition to found epidemics in other locations. These dynamics could create tight bottlenecks between epidemics similar to those that occur between hosts, resulting in dramatic epidemic-to-epidemic diversity losses. That said, if immune hosts are present at the start of an epidemic, there will not be asynchrony between diversity and selection pressure, so new variants may pass through between-epidemic bottlenecks more readily than through between-host bottlenecks. Further work is needed to elucidate mechanisms at the population and meta-population scales.

#### Population immunity sets the clock of antigenic evolution

Our work suggests a simple mechanism by which accumulating immunity to an antigenic variant could produce punctuated population-level antigenic evolution. Population-level modeling has shown that influenza virus global epidemiological and phylogenetic patterns can be reproduced if new antigenic variants emerge at the population level with increasing frequency the longer an old antigenic variant circulates (Koelle et al., 2009).

If immunity to the old variant only gives new variants a population-level transmission advantage (population-level selection), we anticipate a constant rate of population-level antigenic diversification, with selective sweeps once a new variant has a sufficient population-level advantage over the old variant. If population immunity also helps new variants become the dominant within-host variant through inoculation selection, increasing population immunity to an old variant can produce increasing rates of new variant emergence (Fig. 6A). Whether this occurs in practice depends on the ecology of hosts and the relative strength of inoculation selection versus drift in partially and fully immune hosts (Fig. 4).

Potential synergy between brief antigenic replication selection late in infection and subsequent inoculation selection (Fig. 4) could further promote new variant emergence as population immunity accumulates. However, it is difficult to estimate how much this synergy matters in practice without knowing more about the kinetics of homotypic and heterotypic re-inoculation and reinfection.

One suggestive population-level pattern is that new antigenic variants frequently are observed quasi-simultaneously on multiple branches of the influenza virus phylogeny shortly prior to sweeping (see Appendix section 6). This suggests that emergence rate and population-level selection pressure do increase together. That said, alternative explanations are possible, such as reduced rates of stochastic loss of new variants at higher scales (e.g. the epidemic scale) with increased population immunity.

Regardless of the strength of these effects, however, our prediction is that new antigenic variant infection foundation events should constitute a rare but non-negligible fraction of transmission events: on the order of 1 in 10^5^ or 1 in 10^4^. This parsimoniously explains both why new antigenic variants lineages are hard to observe prior to undergoing positive population level selection but are readily available to be selected upon (see Appendix section 6) once that population level selection pressure has become sufficiently strong. Prior to that point they could be rendered unobservable by population-level neutralizing dynamics or by clonal interference from non-antigenic weakly adaptive mutants (Strelkowa & Lässig, 2012).

The slow rate of antigenic evolution of A/H3N2 viruses in swine lends further support to this argument. A/H3N2 viruses accumulate genetic mutations at a similar pace in swine and humans, but antigenic evolution is much slower in swine (De Jong et al., 2007; Lewis et al., 2016). Slaughter for meat means that pig population turnover is high. It follows that the frequency of experienced hosts rarely becomes sufficient to facilitate the appearance of observable new antigenic variants.

The population-level emergence of new antigenic variants, in other words, tracks the accumulation of immunity in the population, not the accumulation of genetic diversity. This suggests that A/H3N2 evolution is indeed selection-limited, not diversity-limited. But much generated antigenic diversity is invisible to surveillance: in the absence of positive selection, it is likely to be lost at bottlenecks.

#### Implications for other pathogens

The inoculation and replication selection paradigm has implications for the understanding and management of other pathogens. For example, HIV does not readily evolve resistance to contemporary pre-exposure prophylaxis (PrEP) antiviral drugs, but it can do so when these antivirals are taken by an individual who is already infected with HIV (Weis et al., 2016). Developing resistance at the moment of exposure is a difficult problem of inoculation selection for the virus, but developing resistance during an ongoing infection is an easier problem of replication selection. Selection on small un-diverse introduced populations may also be of interest in invasion biology and island biogeography.

## Conclusion

The asynchrony between within-host virus diversity and antigenic selection pressure provides a simple mechanistic explanation for the phenomenon of weak within-host selection but strong population-level selection in seasonal influenza virus antigenic evolution. Measuring or even observing antibody selection in natural influenza virus infections is likely to be difficult because it is inefficient and consequently rare. Theoretical studies are therefore essential for understanding these phenomena and for determining which measurable quantities will facilitate influenza virus control. Our study highlights a critical need for new insights into sIgA neutralization and IgA responses to natural influenza virus infection and vaccination and shows that cross-scale dynamics can decouple selection and diversity, introducing randomness into otherwise strongly adaptive evolution.

## Methods

### Model notation

In all model descriptions, *X* += *y* and *X* – = *y* denote incrementing and decrementing the state variable *X* by the quantity *y*, respectively, and *X* = *y* denotes setting the variable *X* to the value *y*. 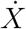 denotes the rate of event *X*.

### Within-host model overview

The within-host model is a target cell-limited model of within-host influenza virus infection with three classes of state variables:

- C: Target **C**ells available for the virus to infect. Shared across all virus variants.
- *V_i_*: **V**irions of virus antigenic variant *i*
- *E_i_*: Binary variable indicating whether the host has previously **E**xperienced infection with antigenic variant *i* (*E_i_* = 1 for experienced individuals; *E_i_* = 0 for naive individuals).

New virions are produced through infections of target cells by existing virions, at a rate *βCV_i_*. Infection eventually renders a cell unproductive, so target cells decline at a rate 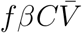, where 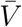 is the total number of virions of all variants. The model allows mutation: a virion of antigenic variant *i* has some probability *μ_ij_* of producing a virion of antigenic variant *j* when it reproduces.

Virions have a natural per-capita decay rate *d_v_*. A fully active specific antibody-mediated immune response to variant *i* increases the virion per-capita decay rate for variant *i* by a factor *k* (assumed equal for all variants). The degree of activation of the antibody response during an infection is given by a function *M*(*t*), where *t* is the time since inoculation.

We use a parameter *c_ij_* to denote the protective strength of antibodies raised against a strain *j* against a different strain *i*. *c_ii_* = 1 by definition and *c_ij_* = 0 indicates complete absence of cross-protection. So if host has antibodies against a strain *j* but not against the infecting strain *i*, a fully active antibody response raises the virion decay rate for strain *i* by *c_ij_k*. If there are multiple candidate forms of cross protection *c_ij_* and *c_ik_*, we choose the strongest. We typically assume that *c_ij_* = *c_ji_*.

We assume that an antibody immune response is raised whenever the host has experienced a prior infection with a partially cross-reactive strain. For notational ease, we define the host’s strongest cross-reactivity against strain *i*, *c_i_* by:

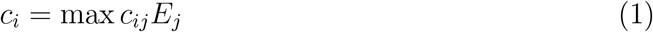

So a recall antibody response is raised during an infection with strains *i*, *j*, … whenever one of *c_i_*, *c_j_*, … > 0.

For this study, we consider a two-antigenic variant model with an ancestral “old antigenic variant” virions *V_w_* and novel “new antigenic variant” virions *V_m_*, though the model generalizes to more than two variants.

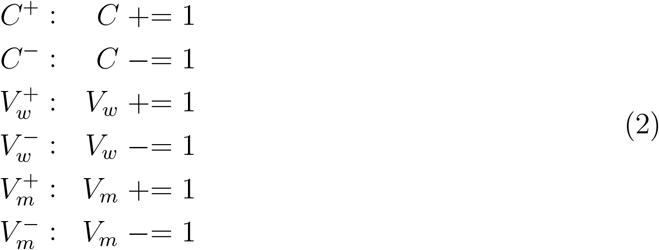

The events occur at the following rates:

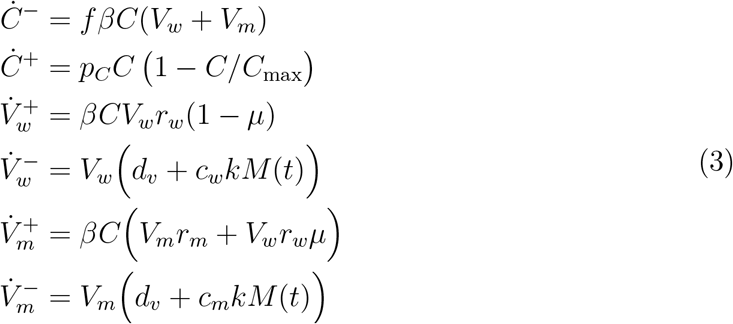

where *M*(*t*) is a minimal model of a time-varying antibody response given by:

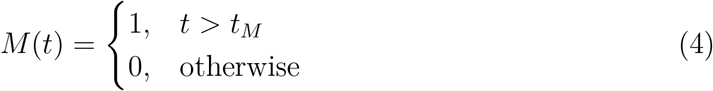

For simplicity, the equations are symmetric between old antigenic variant and new antigenic variant viruses, except that we neglect back mutation, which is expected to be rare during a single infection, particularly before mutants achieve large populations. The parameters *r_w_* and *r_m_* allow the two variants optionally to have distinct within-host replication fitnesses; for all results shown, we assumed no replication fitness difference (*r_w_* = *r_m_* = *r*), unless otherwise stated. A *de novo* antibody response raised to a not-previously-encountered variant *i* can be modeled by setting *E_i_* = 1 at a time 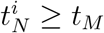 post-infection. By default, we model such a *de novo* response only for fully naive hosts, and assume that it is mounted against the majority variant at the start of infection, which is typically *w*.

We characterize a virus variant *i* by its within-host basic reproduction number 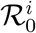, the mean number of new virions produced before decay by a single virion at the start of infection in a naive host:

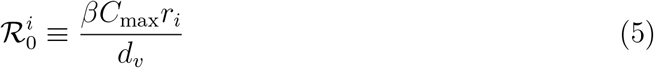

When parametrizing our model, we fixed the within-host basic reproduction number 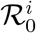, the initial target cell population *C_max_*, the virus reproduction rate *r_i_*, and the shared virion decay rate *d_v_*. We then calculated the implied *β* according to (5).

Another useful quantity is the within-host effective reproduction number 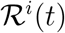 of variant *i* at time *t*: the mean number of new virions produced before decay by a single virion of variant *i* at a given time *t* post-infection.

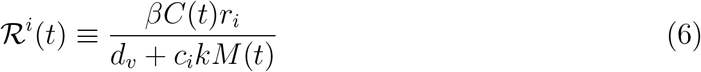

Note that 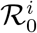 is 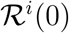 in a naive host, and that if 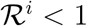, the virus population will usually decline.

The distribution of virions that encounter sIgA antibody neutralization depends on the mean mucosal bottleneck size *v* (i.e. the mean number of virions that would pass through the respiratory tract mucosa in the absence of antibodies) and on the frequency of new antigenic variant *f*_mt_ = *f*_m_ (*t*_inoc_) in the donor host at the time of inoculation *t*_inoc_. *n_w_* old antigenic variant virions and *n_m_* new antigenic variant virions encounter sIgA. The total number of virions *n_tot_* = *n_w_* + *n_m_* is Poisson distributed with mean *v* and each virion is independently a new antigenic variant with probability *f*_mt_ and otherwise an old antigenic variant. The principle of Poisson thinning then implies:

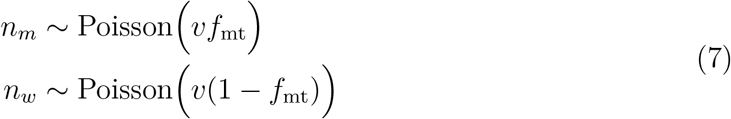

Note that since *f*_mt_ is typically small the results should also hold for a binomial model of *n_w_* and *n_m_* with a fixed total number of virions encountering sIgA antibodies: *v*_tot_ = *v*.

We then model the sIgA bottleneck—neutralization of virions by mucosal sIgA antibodies. Each virion of variant *i* is independently neutralized with a probability *κ_i_*. This probability depends upon the strength of protection against homotypic reinfection *κ* and the sIgA cross immunity between variants *σ*, (0 ≤ *σ* ≤ 1):

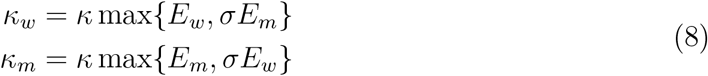

Since each virion of strain *i* in the inoculum is independently neutralized with probability *κ_i_*, then given *n_w_* and *n_m_*, the populations that compete the pass through the cell infection bottleneck *x_w_* and *x_m_* are binomially distributed:

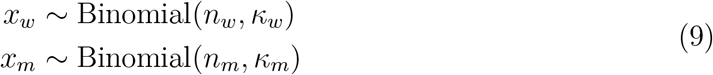

By Poisson thinning, this is equivalent to:

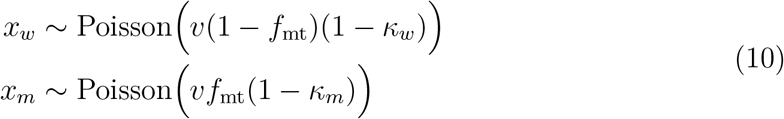

At this point, the remaining virions are sampled without replacement to determine what passes through the cell infection bottleneck, *b*, with all virions passing through if *x_w_* + *x_m_ ≤ *b**, so the final founding population is hypergeometrically distributed given *x_w_* and *x_m_*.

If *κ_w_* = *κ_m_* = 0, *f*_mt_ is small, and *v* is large, this is approximated by a binomially distributed founding population of size *b*, in which each virion is independently a new antigenic variant with (low) probability *f*_mt_ and is otherwise an old antigenic variant. Alternatively, it can be approximated by a Poisson distribution with a small mean: *f*_mt_*b* ≪ 1:

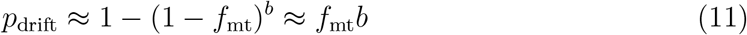

When there is mucosal antibody neutralization, the variant’s survival probability can be reduced below this (inoculation pruning) or promoted above it (inoculation promotion), depending upon parameters. There can be inoculation pruning even when the variant is more fit (neutralized with lower probability, positive inoculation selection) than the old antigenic variant (see Fig. 3E).

### Within-host model parameters

Default parameter values for the minimal model and sources for them are given in Table 1.

### Selection and drift within hosts

In this section, we derive analytical expressions for the within-host frequency of the new variant over time in an infected host (Fig. 7, the probability distribution of the time of first *de novo* mutation to produce a surviving new variant lineage, and the approximate probability of replication selection to frequency *x* by time *t* given our parameters.

**Fig. 7.**
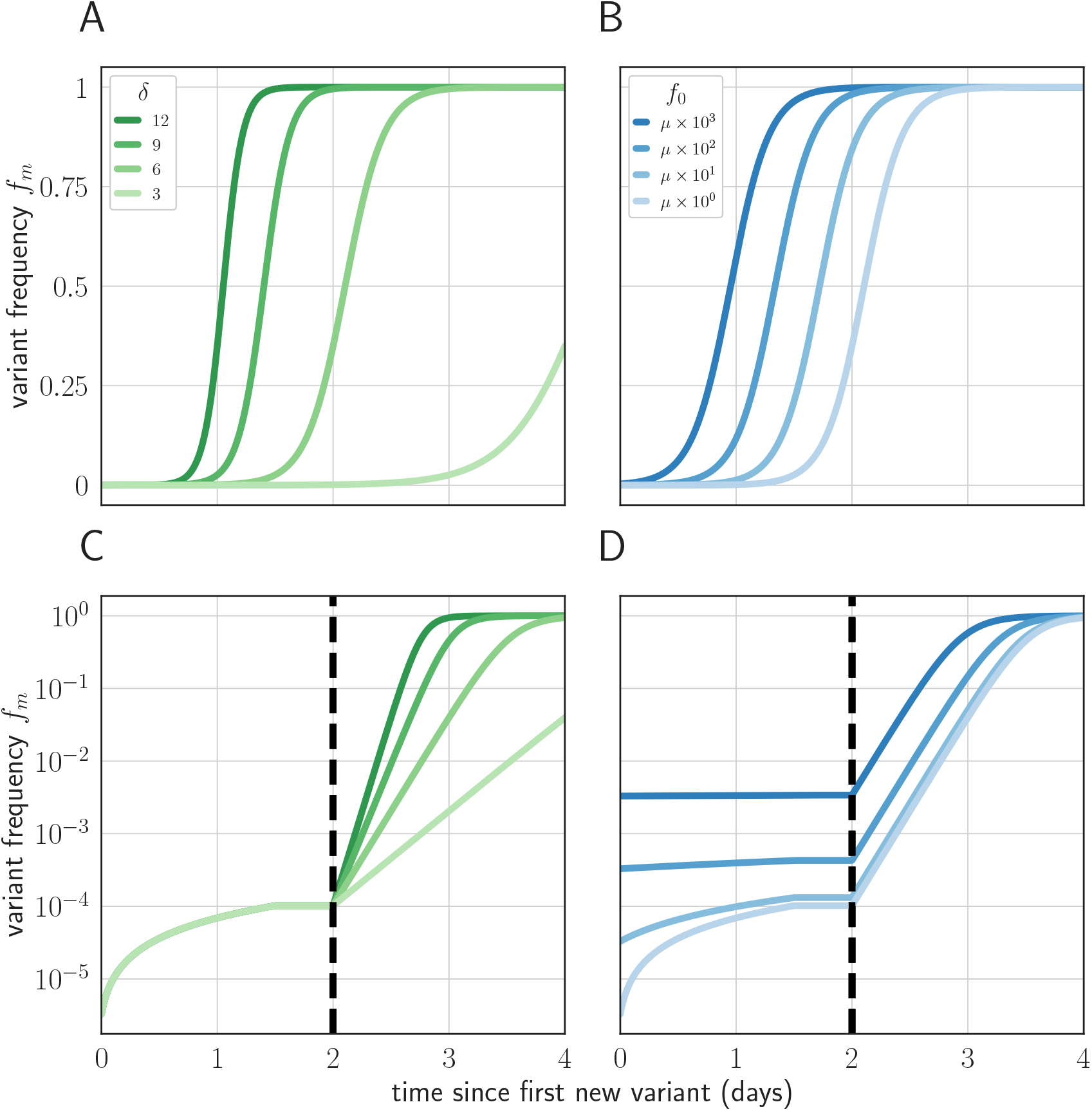
Variant within-host frequency as a function of time and initial variant frequency, according to derived replicator equation (equations 13, 15). (**A, B**) Variant frequency over time for an initially present new variant. (**A**) selection strength *δ* varied, with initial frequency *f*_0_ equal to the mutation rate *μ* = 0.33 × 10^−5^. (**B**) initial frequency *f*_0_ varied, with *δ* = 6. (**C, D**) Variant frequency over time when antigenic selections begins at *t* = 2 days after first variant emergence, with ongoing mutation prior to that point. (**C**) *δ* varied and *f*_0_ fixed as in (**A**); (**D**) *f*_0_ varied and *δ* fixed as in (**B**). Parameters as in Table 1 unless otherwise noted.

#### Within-host replicator equation

The within-host frequency of the new variant, *f*_m_, obeys a replicator equation of the form:

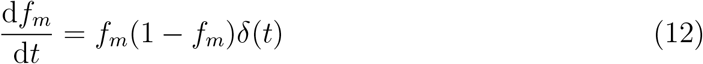

where *δ*(*t*) is the fitness advantage of the new antigenic variant over the old antigenic variant at time *t* (see Appendix section 3 for a derivation).

If the variant is neutral in the absence of antibodies, then *δ*(*t*) = *k*(*c_w_* – *c_m_*) if *t* > *t_M_* and *δ*(*t*) = 0 otherwise. If the new antigenic variant has emerged by *t_M_*, then at a time *t* > *t_M_*:

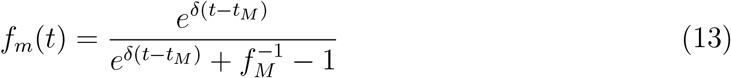

where *δ* = *k*(*c_w_* – *c_m_*) and *f_M_* = *f_m_*(*t_M_*)

When additional mutations after the first cannot be neglected, we add a correction term to 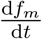 for *t_e_* < *t* < min{*t_M_*, *t*_peak_} (Fig. 7):

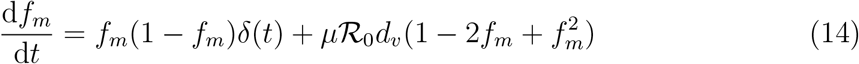

which for *δ* = 0 yields:

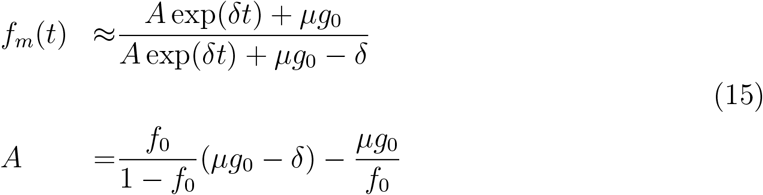

And when *δ* = 0:

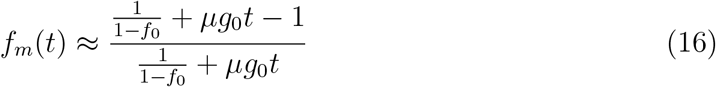

See Appendix section 3.2 for derivations and discussion.

#### Distribution of first mutation times

In our stochastic model, new variant lineages that survive stochastic extinction are produced by *de novo* mutation according to a continuous-time, variable-rate Poisson process. The cumulative distribution function for the time of the first successful mutation, *t_e_*, depends on the mutation rate *μ* and the per-capita rate at which old antigenic variant virions are produced, *g_w_*(*t*) = *r_w_βC*(*t*). It also depends on *p*_sse_, the probability that the generated new antigenic variant survives stochastic extinction. Denoting the new variant per-capita virion production rate *g_m_*(*t*) = *r_m_βC*(*t*), we calculate *p*_sse_ using a branching process approximation ((Ball et al., 2016)).

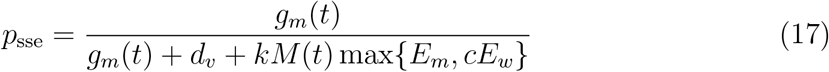

Surviving mutants therefore occur at a rate *λ_m_*(*t*):

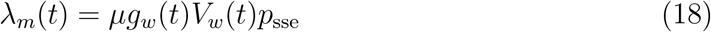

We define the cumulative rate Λ*_m_*(*x*):

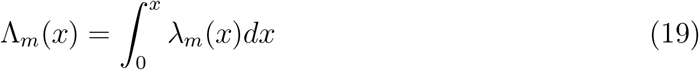

It follows that the CDF of surviving new variant mutation times is:

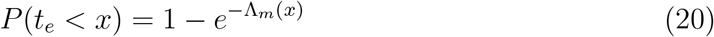

This expression is exact for any given realization of the stochastic model if the realized values of the random variables *V_w_*(*t*), *C*(*t*), and *g_w_*(*t*) are used. In practice, we mainly use it to get a closed form for the approximate new variant CDF by making the approximations that *C*(*t*) ≈ *C*_max_ early in infection. This yields approximations for *g_w_*(*t*), *g_m_*(*t*), and *V_w_*(*t*):

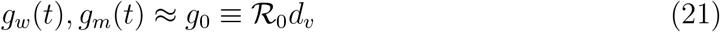

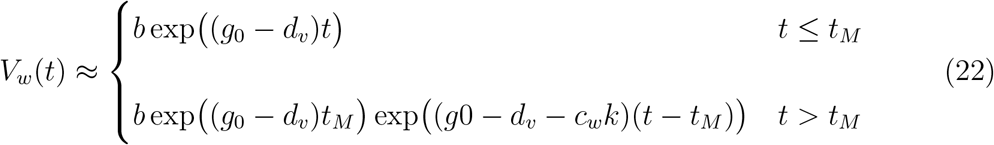

The resultant approximate solution for the CDF of new variant mutation times agrees well with simulations (Fig. 8).

**Fig. 8.**
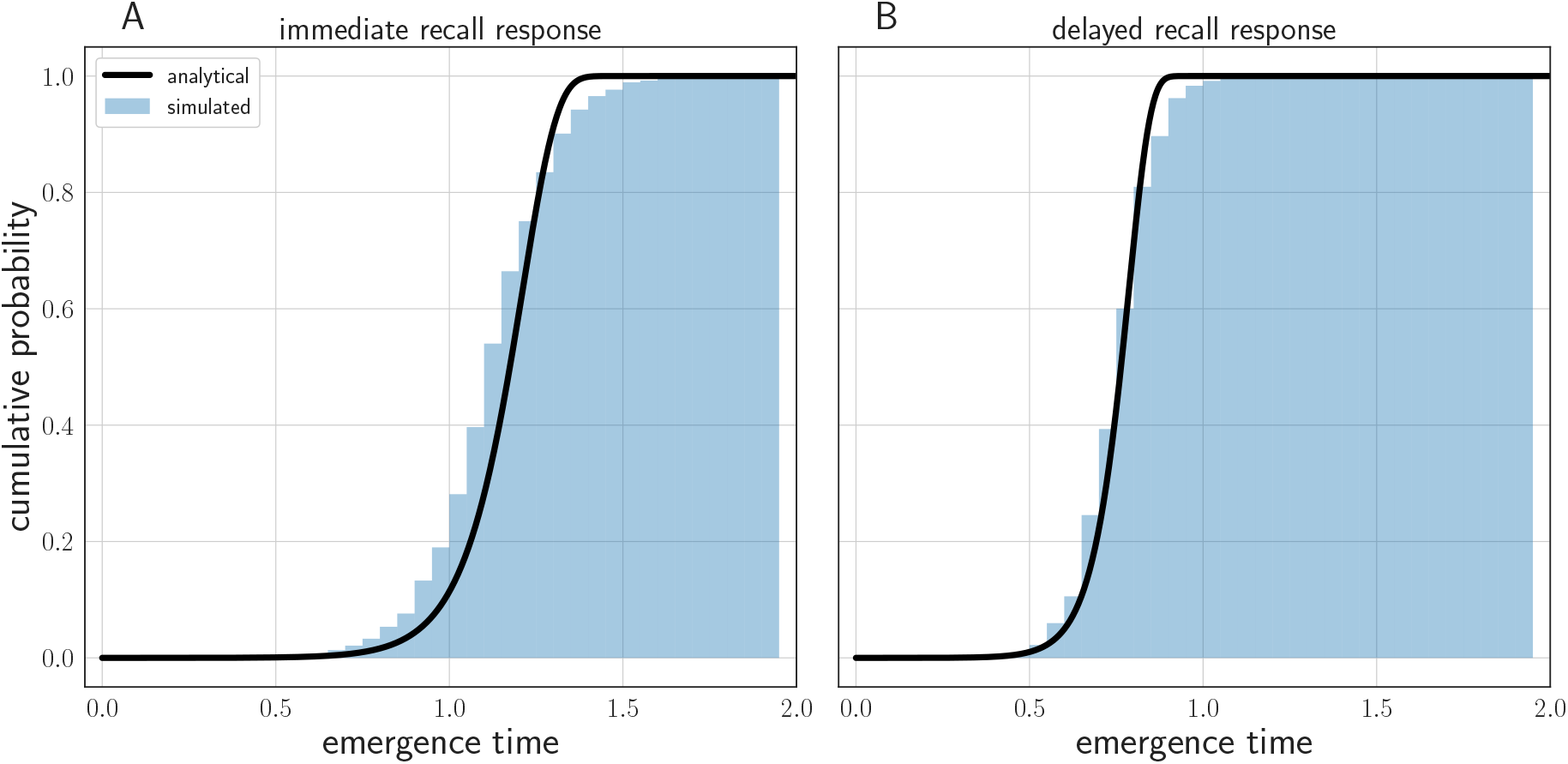
Comparison of analytically calculated cumulative distribution function (CDF) for time of first successful *de novo* mutation with simulations. Black line shows analytically calculated CDF. Blue cumulative histogram shows distribution of new variant mutation times for 250 000 simulations from the stochastic within-host model with (**A**) an immediate recall response (*t_M_* = 0) and (**B**) a realistic recall response at 48 hours post-infection (*t_M_* = 2). Other model parameters as parameters in Table 1. Note that time of first successful mutation tends to be later with an immediate recall response than with a delayed recall response. This occurs because the cumulative number of viral replication events grows more slowly in time at the start of the infection because of the strong, immediate recall response.

The slightly earlier simulated mutation times in the immediate recall response case (Fig. 8A) would only make replication selection more likely in that case than our analytical approximation suggests.

#### Required mutation time for variant to reach a given frequency

By inverting the within-host replicator equation, we can also calculate the time *t*\*(*x*, *t*) by which a new variant must emerge if it is to reach at least frequency *x* by time *t*. We show (see Appendix section 9.4 for derivation) that there are two candidate values for *t*\*, depending on whether the time that the new variant first emerges (*t_e_*) is before or after the onset of the antibody response (*t_M_*):

If *t_e_* ≤ *t_M_*:

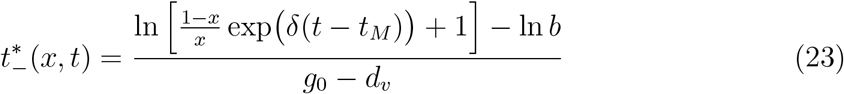

If *t_e_* > *t_M_*:

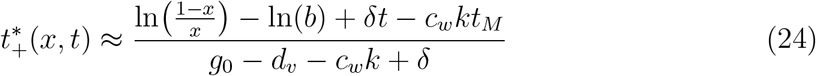

It may be that 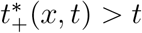 and 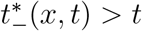. This indicates that the mutant will be at frequency *x* if it emerges at *t* itself. In that case, we therefore have *t*\*(*x*, *t*) = *t*. So combining:

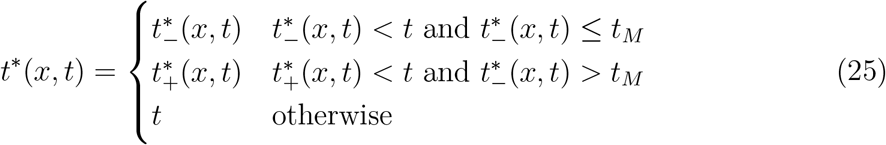

Finally, it is worth noting that in the case of a complete escape mutant (*c_m_* = 0, *δ* = *c_w_k*), the approximate expression for 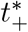 is exactly equal to the equivalent approximate expression for 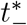:

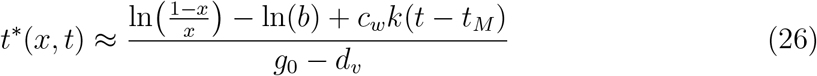

This is a linear function of *t*.

#### Probability of replication selection

Given this and the the new variant first mutation time CDF calculated in equation 20, it is straightforward to calculate the probability of replication selection to a given frequency *a* by time *t*, assuming that 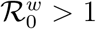 early in infection:

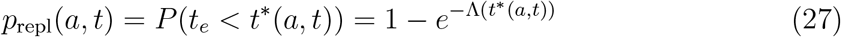

This analytical model agrees well with simulations (Fig. 9). We use it used to calculate the heatmaps shown in Fig. 1, with the *C*(*t*) ≈ *C_max_* early infection approximations that give us a closed form for Λ*_m_*(*x*).

**Fig. 9.**
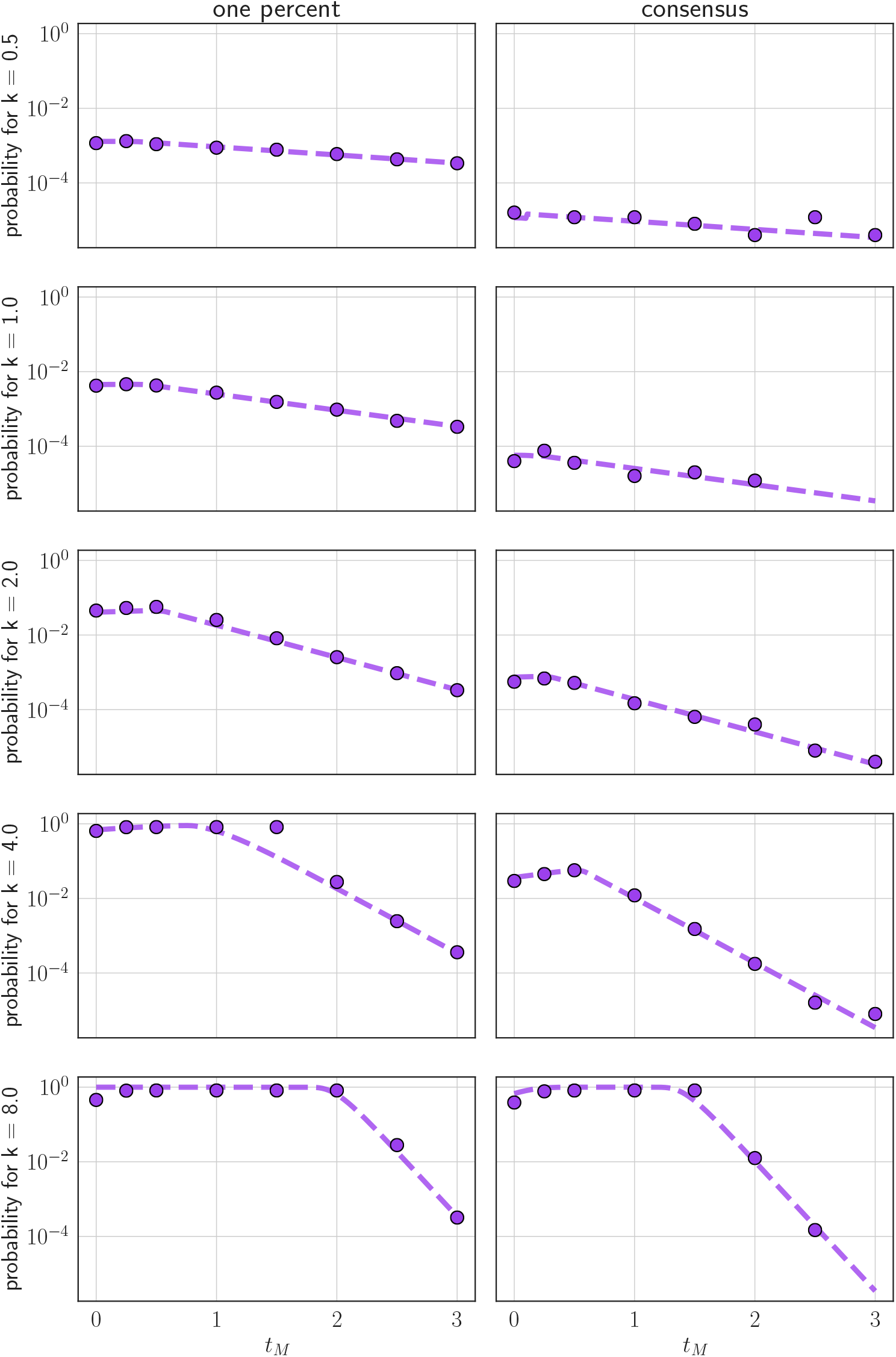
Comparison of analytically calculated probability of replication selection with stochastic simulations. Probability of replication selection to one percent (left column) or consensus (right column) by *t* = 3 days post-infection as a function of *k* and *t_M_* for 250 000 simulations from the stochastic within-host model. *c* = 0 (so the fitness difference *δ* = *k*). Other model parameters as parameters in Table 1. Dashed lines show analytical prediction and dots show simulation outcomes.

When *t*\* > *t_M_*, the integral 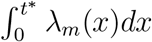 can be evaluated piecewise, first from 0 to *t_M_*, and then from *t_M_* to *t*\*. This can also be used to find evaluate the probability density function (PDF) of first mutation times, as needed.

Finally, note that for *t_e_* < *t_M_*, the expression for *p*_repl_ in practice depends only on *δ* = (*c_w_* – *c_m_*)*k*, not on *c_w_*, *c_m_* and *k* separately. In the absence of an antibody response, the mutant is generated with near certainty by 1 day post-infection (*P*(*t_e_* < 1) ≈ 1, Fig. 8B). So when *t_M_* ≥ 1, values for *p*_repl_ calculated with *c_w_* = 1 *c_m_* = 0, and *δ* = *k*—as in (Fig. 1C,D)—will in fact hold for any *c_w_*, *c_m_*, and *k* that produce that fitness difference *δ*.

When 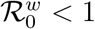 early in infection, the probability of replication selection depends on the probability of generating an escape mutant before the infection is extinguished:

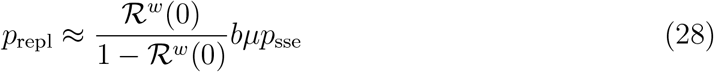

See Appendix section 3.6 for a derivation.

### Point of transmission model

In this section, we describe our model of the point of transmission, including how sIgA neutralization may impose selection pressure.

#### Transmission probability

Given contact between an infected and an uninfected host, we assume that a transmission attempt occurs with probability proportional to the current virus population size *V*_tot_ in the infected host:

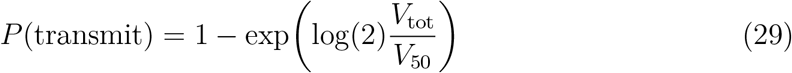

The parameter *V*_50_ sets the scaling, and reflects virus population size at which there is a 50% chance of successful transmission to a naive host. We used a default value of *V*_50_ = 10^8^ virions. In addition to this *probabilistic model*, we also consider an alternative *threshold model* in which a transmission attempt occurs with certainty if *V*_tot_ is greater than a threshold *θ* and does not occur otherwise.

#### Bottleneck survival

If there are *x_w_* old antigenic variant virions and *x_m_* variant virions, at least one variant passes through the cell infection bottleneck with probability:

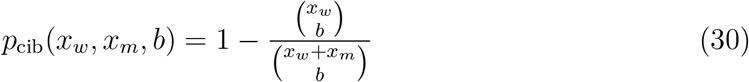

Summing *p*_cib_(*x_w_*, *x_m_*) over the possible values of *x_w_* and *x_m_* weighted by their joint probabilities yields the variant’s overall probability of successful onward transmission *p*_surv_(*f*_mt_, *v*, *b*, *κ*, *c*):

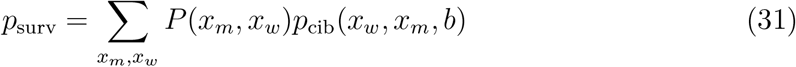

*P*(*x_w_*, *x_m_*) is the product of the probability mass functions for *x_w_* and *x_m_*:

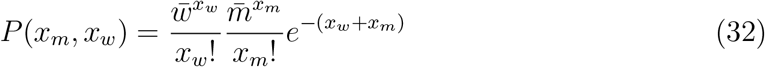

where 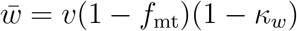 and 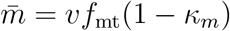

At low donor host variant frequencies *f*_mt_ ≪ 1 *p*_surv_ can be approximated using the fact that almost all probability is given to *x_m_* = 0 and *x_m_* =1 (0 or 1 new antigenic variant after the sIgA bottleneck).

At least one new antigenic variant survives the sIgA bottleneck with probability:

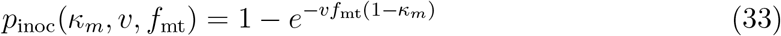

If *f*_mt_ is small, there will almost always be only one such virion (*x_m_* = 1). That new antigenic variant’s probability of surviving the cell infection bottleneck depends upon how many old antigenic variant virions are present after neutralization, *x_w_*:

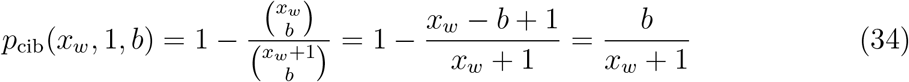

Summing over the possible *x_w_*, we obtain a closed form for the unconditional probability *p*_cib_(*κ_w_*, *v*, *b*) (see section 9.5 for a derivation):

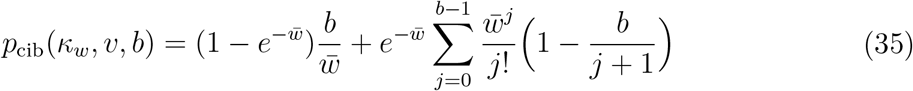

where 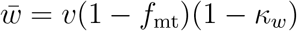 is the mean number of old antigenic variant virions present after sIgA neutralization. If *b* =1, this reduces the just the first term, and it is approximately equal to just the first term when 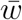 is large, since 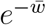 becomes small.

It follows that there is an approximate closed form for the probability that a variant survives the final bottleneck:

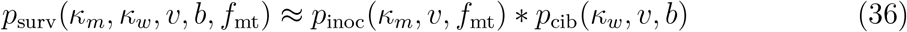

We use this expression to calculate the analytical variant survival probabilities shown in the main text, and to gain conceptual insight into the strength of inoculation selection relative to neutral processes (see Appendix section 4.5).

#### Neutralization probability and probability of no infection

A given per-virion sIgA neutralization probability *κ_i_* implies a probability *z_i_* that a transmission event involving only virions of variant *i* fails to result in an infected cell.

Some transmissions fail even without sIgA neutralization; this occurs with probability exp(–*v*). Otherwise, with probability 1 – exp(–*v*), *n_i_* virions must be neutralized by sIgA to prevent an infection. We define *z_i_* as the probability of no infection given inoculated virions in need of neutralization (i.e. given *n_i_* > 0).

The probability that there are no remaining virions of variant *i* after mucosal neutralization is exp(–*v*(1 – *κ_i_*)). So we have:

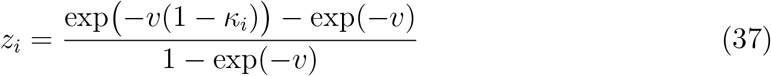

This can be solved algebraically for *κ_i_* in terms of *z_i_*, but it is more illuminating first to find the overall probability of no infection given inoculation *p*_no_ and then solve:

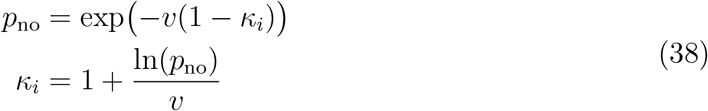

An infection occurs with probability (1 – exp(–*v*))(1 – *z_i_*) (the probability of at least one virion needing to be neutralized times the conditional probability that it is not) so:

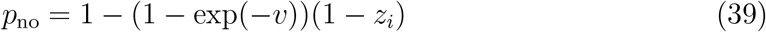

This yields the same expression for *κ_i_* in terms of *z_i_* as a direct algebraic solution of 37.

For moderate to large *v*, 1 – exp(–*v*) approaches 1 so *p*_no_ approaches *z_i_* and *κ_i_* approaches 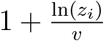. This reflects the fact that for even moderately large *v*, it is almost always the case that *n_i_* > 0: a least one virion must be neutralized to prevent infection. In those cases, *z_i_* can be interpreted as the (approximate) probability of no infection given a transmission event (i.e. as *p*_no_).

Note also that seeded infections can also go stochastically extinct; this occurs with approximate probability 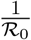. At the start of infection, if there is no antibody response, 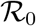 is large (5 to 10) stochastic extinction probabilities should be low (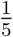 to 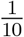), and equal in immune and naive hosts. We have therefore parametrized our model in terms of *z_i_*, the probability that no cell is ever infected, as that probability determines to leading order the frequency with which immunity protects against detectable reinfection given challenge.

#### Susceptibility models

Translating host immune histories into old antigenic variant and new antigenic variant neutralization probabilities for the analysis in Fig. 3G,H requires a model of how susceptibility decays with antigenic distance, which we measure in terms of the typical distance between two adjacent “antigenic clusters” (Smith et al., 2004). Fig. 3 shows results for two candidate models: a multiplicative model used in a prior modeling and empirical studies of influenza evolution (Asaduzzaman et al., 2018; Boni et al., 2006), and a sigmoid model as parametrized from data on empirical HI titer and protection against infection (Coudeville et al., 2010).

In the multiplicative model, the probability *z*(*i*, *x*) of no infection with variant *i* given that the nearest variant to *i* in the immune history is variant *x* is given by:

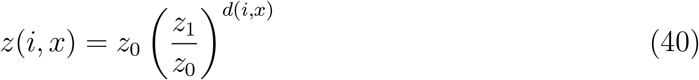

where *z*_0_ is the probability of no infection given homotypic reinfection, *d*(*i*, *x*) is the antigenic distance in antigenic clusters between *i* and *x*, and *z*_1_ is the probability of no infection given *d*(*i*, *x*) = 1.

In the sigmoid model:

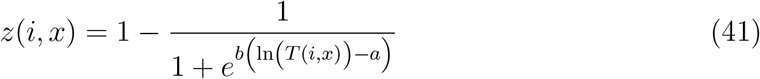

where *a* = 2.844 and *b* = 1.299 (*α* and *β* estimated in (Coudeville et al., 2010)) and *T*(*i*, *x*) is the individual’s HI titer against variant *i* (Coudeville et al., 2010). To convert this into a model in terms of *z*_0_ and *z*_1_ we calculate the typical homotypic titer *T*_0_ implied by *z*_0_ and the *n*-fold-drop *D* in titer per unit of antigenic distance implied by *z*_1_, since units in antigenic space correspond to a *n*-fold reductions in HI titer for some value *n* (Smith et al., 2004). We calculate *T*_0_ by plugging *z*_0_ and *T*_0_ into equation 41 and solving. We calculate *D* by plugging *z*_1_ and *T*_1_ = *T*_0_*D*^−1^ into equation 41 and solving. We can then calculate *T*(*i*, *x*) as:

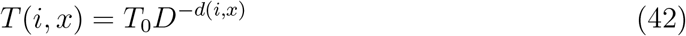

### Probability of bottleneck survival

With these analyses in hand, it is possible to combine the within-host and the point of transmission processes to calculate an overall probability that a new variant survives the transmission bottleneck. Equation 36 gives the probability of bottleneck survival given *f*_mt_ and the properties of the recpient host. And given a time of transmission *t_t_*, we can calculate the probability distribution of *f*_mt_ using our expressions for the CDF of successful mutation times *t_e_* (equation 20) and for *f_m_*(*t*) (equations 13, 15). To calculate the overall probability *p*_nv_, we average over the possible values of *f*_mt_, weighted by their probability:

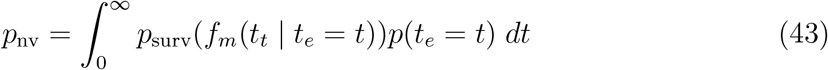

### Within-host simulations (Figs. 1, 3)

To evaluate the relative probabilities of replication selection and inoculation selection for antigenic novelty and to check the validity of the analytical results, we simulated 10^6^ transmissions from a naive host to a previously immune host. The transmitting host was simulated for 10 days, which given the selected model parameters is sufficient time for almost all infections to be cleared. Time of transmission was randomly drawn from the time period during which the infected host had a transmissible virus population and an inoculum of fixed size was drawn from the within-host virus population at that time. Variant counts in the inoculum were Poisson distributed with probability equal to the variant frequency within the transmitting host at the time of transmission. We simulated the recipient host until the clearance of infection and found the maximum frequency of transmissible variant: that is, the variant frequencies when the transmission probability was greater than 5 × 10^−2^ (probabilistic model) or when the virus population was above the transmission threshold θ (threshold model). For Fig. 3D, we defined an infection with an emerged new antigenic variant as an infection with a maximum transmissible new variant frequency of greater than 50%.

### Transmission chain model (Fig. 5)

To study evolution along transmission chains with mixed host immune statuses, we modeled the virus phenotype as existing in a 1-dimensional antigenic space. Host susceptibility *s_x_* to a given phenotype *x* was *s_x_* = min{1,min{ļ*x* – *y_i_*|}} for all phenotypes *y_i_* in the host’s immune history.

Within-host cross immunity *c_ij_* between two phenotypes *y_i_* and *y_j_* was equal to max{0, 1 – |*y_i_* – *y_j_*|}. When mucosal antibodies were present, protection against infection *z_x_* was equal to the strength of homotypic protection *z*_max_ scaled by susceptibility: *z_x_* = (1 – *s_x_*)*z_max_*. *κ_x_* was calculated from *z_x_* by exp(–*v*(1 – *κ_x_*) = *z_x_*. We used *z*_max_ = 0.95. Note that this puts us in the regime in which intermediately immune hosts are the best inoculation selectors (Fig. 3H). We set *k* = 25 so that there would be protection against reinfection in the condition with an immediate recall response but without mucosal antibodies.

We then simulated a chain of infections as follows. For each inoculated host, we tracked two virus variants: an initial majority antigenic variant and an initial minority antigenic variant. If there were no minority antigenic variants in the inoculum, a new focal minority variant (representing the first antigenic variant to emerge *de novo*) was chosen from a Gaussian distribution with mean equal to the majority variant and a given variance, which determined the width of the mutation kernel (for results shown in Fig. 5, we used a standard deviation of 0.08). We simulated within-host dynamics in the host according to our within-host stochastic model.

We founded each chain with an individual infected with all virions of phenotype 0, representing the current old antigenic variant.

We simulated contacts at a fixed, memoryless contact rate *ρ* =1 contacts per day. Given contact, a transmission attempt occurred with a probability proportional to donor host viral load, as described above. If a transmission attempt occurred, we chose a random immune history for our recipient host according to a pre-specified distribution of host immune histories. We then simulated an inoculation and, if applicable, subsequent infection, according to the within-host model described above. If the recipient host developed a transmissible infection, it became a new donor host. If not, we continued to simulate contacts and possible transmissions for the donor host until recovery. If a donor host recovered without transmitting successfully, the chain was declared extinct and a new chain was founded.

We iterated this process until the first phenotypic change event—a generated or transmitted minority phenotype becoming the new majority phenotype. We simulated 1000 such events for each model and examined the observed distribution of phenotypic changes compared to the mutation kernel.

For the results shown in Fig. 5, we set the population distribution of immune histories as follows: 20% −0.8, 20% −0.5, 20% 0.0, and the remaining 40% of hosts naive. This qualitatively models the directional pressure that is thought to canalize virus evolution (Bedford et al., 2012) once a cluster has begun to circulate.

### Analytical mutation kernel shift model (Fig. 5)

To assess the causes of the observed behavior in our transmission chain model, we also studied analytically how replication and inoculation selection determine the distribution of observed fixed antigenic changes given the mutation kernel when a host with one immune history inoculates another host with a different immune history.

We fixed a transmission time *t* = 2 days, roughly corresponding to peak within-host virus titers. For each possible new variant phenotype, we calculated *p*_nv_ according to equation 43, with parameters given by the the old variant antigenic phenotype, new variant phenotype, and host immune histories. Finally, we multiplied each phenotype’s survival probability by the same Gaussian mutation kernel used in the chain simulations (with mean 0 and s.d. 0.08), and normalized the result to determine the predicted distribution of surviving new variants given the mutation kernel and the differential survival probabilities for different phenotypes.

### Population-level model (Fig. 6)

To evaluate the probability of variants being selected and proliferating during a local influenza virus epidemic, we first noted that the per-inoculation rate of new antigenic variant chains for a population with *n_s_* susceptibility classes (which can range from full susceptibility to full immunity) is:

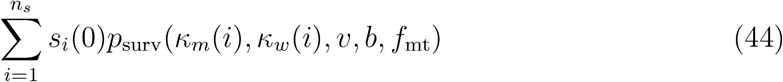

where *κ_w_*(*i*) and *κ_m_*(*i*) are the mucosal antibody neutralization probabilities for the old antigenic variant and the new antigenic variant associated with susceptibility class *i*, and 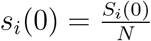 is the initial fraction of individuals in susceptibility class *i*.

We then considered a well-mixed population with frequency-dependent transmission, where infected individuals from all susceptibility classes are equally infectious if infected. Using an existing result from epidemic theory (Magal et al., 2018), we calculated *R*_∞_, the average fraction of individuals who are infected if an epidemic occurs in such a population. During such an epidemic, each individual will on average be inoculated (challenged) 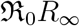 times, where 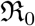 is the population-level basic reproduction number (Miller, 2012). We can then calculate the probability that a new variant transmission chain is started in an arbitrary focal individual:

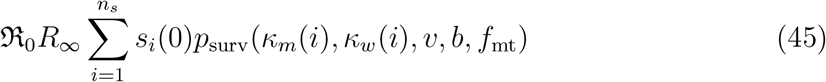

### Sensitivity analysis (Fig. A3)

We assessed the sensitivity of our results to parameter choices by re-running our simulation models with randomly generated parameter sets chosen via Latin Hypercube Sampling from across a range of biologically plausible values. Figure A3 gives a summary of the results.

We simulated 50 000 infections of experienced hosts (*c_w_* = 1) according to each of 10 random parameter sets. We selected parameter sets using Latin Hypercube sampling to maximize coverage of the range of interest without needing to study all possible permutations. We did this for the following bottleneck sizes: 1, 3, 10, 50.

We analyzed two cases: one in which the immune response is unrealistically early and one in which it is realistically timed. In the unrealistically early antibody response model, *t_M_* varied between *t_M_* = 0 and *t_M_* = 1. In the realistically timed antibody response model, *t_M_* varied between *t_M_* = 2 and *t_M_* = 4.5. Other parameter ranges were shared between the two models (Table 2).

**Table 2:** Sensitivity analysis parameter ranges shared between models.

| Parameter | Minimum value | Maximum value |
| --- | --- | --- |
| $t_N$ | 6 | 9 |
| $C_{\text{max}}$ | $10^8$ | $10^9$ |
| $\mathcal{R}_0$ | 5 | 15 |
| $r$ | 10 | 500 |
| $z_w$ | 0.70 | 0.99 |
| $z_m/z_w$ | 0.5 | 0.9 |
| $\mu$ | $0.33 \times 10^{-6}$ | $0.33 \times 10^{-4}$ |
| $d_v$ | 2 | 8 |
| $k$ | 3 | 16 |
| $c_m$ | 0.5 | 1 |
| $\theta$ | $10^6$ | $10^8$ |
| $v/b$ | 1 | 50 |

Discussion of sensitivity analysis results can be found in Section 8 below.

### Meta-analysis (Fig. 1)

We downloaded processed variant frequencies and subject metadata from the two NGS studies of immune-competent human subjects naturally infected with A/H3N2 with known vaccination status (Debbink et al., 2017; McCrone et al., 2018) from the study Github repositories and journal websites. We independently verified that reported antigenic site and antigenic ridge amino acid substitutions were correctly indicated, determined the number of subjects with no NGS-detectable antigenic amino acid substitutions, and produced figures from the aggregated data.

### Computational methods

For within-host model stochastic simulations, we used the Poisson Random Corrections (PRC) tau-leaping algorithm (Hu & Li, 2009). We used an adaptive step size; we chose step sizes according to the algorithm of Hu and Li (Hu & Li, 2009) to ensure that estimated next-step mean values were non-negative, with a maximum step size of 0.01 days. Variables were set to zero if events performed during a timestep would have reduced the variable to a negative value. For the sterilizing immunity simulations in Figure 2, we used a smaller maximum step size of 0.001 days in recipient hosts to better handle mutation dynamics involving very small numbers of replicating virions.

We obtained numerical solutions of equations, including systems of differential equations and final size equations, in Python using solvers provided with SciPy (Jones et al., 2001).

## Acknowledgments

We thank Daniel B. Cooney, Andrea L. Graham, Joseph Gibson, Katelyn M. Gostic, Alvin X. Han, Chadi M. Saad-Roy, and Edward C. Schrom for helpful discussions. We thank Christopher J. Illingworth, Daniel B. Weissman, and an anonymous reviewer for helpful comments on a prior version of this manuscript.

We thank the GISAID Initiative and the influenza surveillance and research groups who openly shared the genetic sequence data used in this work (a table with accession numbers and associated labs is available online in the project materials and data repositories on Github and OSF, see below for links).

## Author contributions

CAR, DHM, and VNP conceived and designed the research. DHM created and analyzed the mathematical models, with support from FWR, RAN, and SAL. DHM analyzed the within-host data. EP conducted phylogenetic analyses. DHM produced the figures. All authors contributed substantially to the interpretation of results and the writing and revising of the manuscript.

## Data and materials availability

All code, data, and other materials needed to reproduce the analysis in this paper are provided online on the project Github repository: https://github.com/dylanhmorris/asynchrony-influenza

They are also available on OSF: https://doi.org/10.17605/OSF.IO/jdsbp/

Output data generated by stochastic simulations, within-host NGS meta-analysis, and phylogenetic analyses are archived on OSF: https://doi.org/10.17605/OSF.IO/jdsbp/

