## Appendix for "Asynchrony between virus diversity and antibody selection limits influenza virus evolution"

October 28, 2020

### Contents

**1 Summary** 2

**2 Immunological underpinnings of the inoculation selection model** 2

**3 Detailed within-host model analysis** 5

**4 Point of transmission analysis** 11

**5 Parameter uncertainties and sensitivity analysis** 16

**6 Polyphyletic antigenicity altering substitutions** 28

**7 Prior theoretical studies** 36

**8 Further implications** 42

**9 Mathematical derivations in full** 46

### 1 Summary

This appendix contains additional information for “Asynchrony between virus diversity and antibody selection limits influenza virus evolution”.

We provide a detailed review of virological and immunological evidence supporting our model of the interaction between influenza viruses and the (human) host immune system (Section 2). We give a more detailed mathematical analysis of our within-host model, deriving the approximate and exact analytical results that are used in the main text and provide intuition to explain model behavior and limitations (Section 3). We do the same for our model of the point of transmission, and assess which hosts are most likely to act as inoculation selectors and which mutants are most likely to be inoculation selected (Section 4).

We next discuss parameter uncertainties and report the results of a sensitivity analysis for our within-host models (Section 5). We report an additional empirical analysis showing that new variants often emerge polyphyletically: they appear simultaneously on multiple branches of the influenza virus phylogeny. We discuss how this is more easily explained in light of our model than in light of previous models (Section 6).

We then review these prior studies and models of within-host influenza virus antigenic evolution and general pathogen immune escape in depth. We argue that they are insufficient to explain within-host influenza virus antigenic evolution and that inoculation selection accompanied by a realistically-timed antibody response is the most likely competing hypothesis (Section 7). We elaborate on the conclusions that can be drawn from our study and its implications for influenza virus control and for future basic research (Section 8). We conclude by providing complete step-by-step mathematical derivations of lemmas and approximations from elsewhere in the text (Section 9).

### 2 Immunological underpinnings of the inoculation selection model

In this section, we review relevant immunology to establish the biological realities that we built our model to capture. We also discuss some unmodeled biological complexities, and why they are unlikely to change our qualitative conclusions.

#### 2.1 Strong selection at the point of inoculation

The establishment of successful influenza virus infection requires access to respiratory epithelial cells covered by a mucosal layer, which acts as a protective barrier for virus entry. Sialic acid residues present in the mucus act as decoys for virions and the enzymatic activity of the influenza virus neuraminidase (NA) protein is essential for penetration of the mucosal layer (Matrosovich et al., 2004). The mucosal layer does not act only as a mechanical barrier. Secretory IgA (sIgA) mucosal antibodies specific to influenza virus proteins can apply virus-specific neutralization by immobilizing virions and slowing down the process of virus infection (Wang et al., 2017). Depending on their specificity, sIgA can either play a role in reduction of total virus population size or in providing specific competitive advantage to mutant viruses. Antibodies against the influenza virus NA protein will slow down the virus passage through the mucosal barrier irrespective of the HA antigenic phenotype of the newly infecting virus. By contrast, anti-HA antibodies are essential for inoculation selection because they recognise previously encountered antigenic variants. This provides a competitive advantage to new, distinct antigenic variants. The new variants with strongest competitive advantage are those with large antigenic effects; they have the lowest probability to be neutralized by any cross-reactive anti-HA sIgAs.

#### 2.2 Weak selection during exponential virus growth

Virions that successfully cross the mucosal barrier can infect epithelial cells in the respiratory tract. Upon initiation of replication in host cells, newly generated virions can be subjected to replication selection via virus-specific antibodies. Replication selection can take place only if influenza virus-specific antibodies are present in mucosa-associated lymphoid tissue (MALT) at the time of virus replication.

Influenza virus-specific antibodies in the MALT are secreted by local plasmablast populations, generated following activation of B naive cells or B memory cells. The exposure history of the host determines the timing and the immunological trajectory for the generation of influenza virus-specific antibodies. In individuals without previous exposure to influenza virus, newly generated virions are recognised by naive B cells which undergo rounds of affinity maturation and somatic hypermutation to produce highly-specific antibodies to the virus. This process typically occurs in germinal centres, requires T-cell help, and takes around 7 days for production of highly-specific antibodies (Wrammert et al., 2008).

In individuals with previous exposure history to influenza viruses, where the newly generated influenza virions are antigenically matched or have some degree of cross-reactivity to previously encountered variants, the generation of influenza virus-specific antibodies mainly results from B memory recall response. The activation of pre-existing memory pools enables the immune system to respond quicker to previously encountered antigens without the need to recruit naive B cell populations. The generation of antigen-specific antibodies from recalled B memory pools does not typically start until 3–5 days post infection (Coro et al., 2006; Lam & Baumgarth, 2019).

The B naive and B memory cells generating serological responses to influenza viruses are referred to as B-2 cells. The activation of these cell types depends on the recognition of virus protein via the B-cell receptor and sufficient affinity of binding to trigger stimulatory signal. Depending on their specificity to the recognised virus antigen, the antibodies produced by these cell types have the potential to apply replication selection to any generated new antigenic variant viruses and to old variant viruses during the course of infection. Antibodies against influenza viruses can also be generated from the so called B-1 cells, which do not require specific recognition of virus antigen and are activated by innate mechanisms (Lam & Baumgarth, 2019).

B-1 cells produce natural IgM antibodies with low affinity which constitute the earliest humoral response generated in the first 48–72h of infection (Lam & Baumgarth, 2019). As the activation of B-1 cells is independent of the specific recognition of influenza virus virions, the natural IgM antibodies act as a non-specific limiting factor for virus population sizes and cannot apply replication selection to particular virus antigenic variants.

The peak of influenza virus replication occurs in the first 24–48 hours post infection (Hadjichrysanthou et al., 2016). Despite the presence of several immunological routes for generation of influenza virus-specific antibodies, the timing of the serological immune response is always later than the time of peak of virus titer (at least in typical infections of otherwise healthy individuals), thus limiting the opportunity for replication selection during the course of infection.

#### 2.3 Selection during exit of an infected host

The design of the inoculation selection model focuses on selection applied by the mucosal layer upon the entry of influenza virus in naïve or previously exposed hosts. We do not take into account the potential bottleneck applied during exit of viruses from the respiratory tract of an infected host. Selection of newly generated virions at the point of exit of an infected host is technically possible as the virions released from infected epithelial cells need to cross the mucosal barrier again to reach the lumen of the respiratory tract. Such selection upon virus exit would have the same directional effect as inoculation selection; it would provide a competitive advantage for mutant viruses with large antigenic effects. The potential strength of such selection depends heavily the how many sIgA antibodies in the transmitting host's mucosa are free to neutralize the virions passing through; this quantity is unknown. Incorporating such mucosal selection upon virus exit thus requires better knowledge of the quantities of sIgA antibodies in given volume of mucus and what proportion of the available sIgA antibodies are engaged with antigen when an infected host transmits. Such data is not currently available and would require well-designed cellular culture models which take into account the role of mucosal layer in the process of virus infection and release of newly generated viruses.

#### 2.4 Nature and timing of previous exposure and effects on inoculation selection

According to our model, the strongest inoculation selection can take place in infection of previously exposed individuals and the highest probability of antigenic diversification occurs in populations with an intermediate proportion of immune hosts. In deriving this result, we have assumed homogeneous and static quantities of sIgA antibodies among previously exposed individuals. This is an oversimplification. There is substantial variation in level of sIgA antibodies across individuals depending on the nature of their previous exposures (through vaccination or natural infection). For example, intranasal inoculation with influenza virus via live attenuated influenza virus vaccines leads to generation of better mucosal immunity (Hoft et al., 2017) than intramuscular administration of trivalent inactivated vaccines and thus previously exposed individuals are likely to vary in the degree of inoculation selection applied during homotypic infections depending on the route of administration of previous vaccinations.

The model also does not account for waning of sIgA antibody titers in the months and years following each exposure. The waning of sIgA titers is likely to be substantial in absence of re-exposure but the waning process should only decrease the efficiency of inoculation selection and make the accumulation of population level selection pressure—and thus the appearance of observable new antigenic variants—rarer.

The dynamics of mucosal immunity with age are also likely to have impact on the selector phenotype of previously exposed hosts. In older individuals, multiple previous exposures will eventually lead to preferential activation of recall responses for antibody production to new variants as a result of original antigenic sin (Davenport & Hennessy, 1956), immune backboosting (Fonville et al., 2014), or antigenic seniority (Lessler et al., 2012).

Even if the recall responses result in backboosting and higher levels of sIgA, these antibodies are unlikely to be well-matched to the new variant or succeeding variants, as continuous re-exposure favours responses to conserved epitopes (Krammer, 2019; Krammer & Palese, 2019) rather than receptor-binding site epitopes suspected to be most important for immune escape.

As a result of the limited ability to generate novel immune responses and the phenomenon of antigenic seniority, an elderly individual who has been exposed multiple times in the past is likely to exert very weak inoculation selection to newly infecting viruses. By contrast, a young person recently exposed to influenza virus is more likely to act as a strong selector, due to the availability of mucosal sIgA antibodies and the more limited impact of original antigenic sin. However, due to the acute nature of influenza virus infection, very young children are likely to require more than one exposure before they begin to develop highly specific antibody responses to influenza virus infection able to exert inoculation selection (Neuzil et al., 2006).

Most current efforts for characterizing humoral immunity to influenza virus infections are focused on antibody responses in the serum. However, there is limited understanding of the relationship between antibodies in the serum measured via HI and the levels of mucosal antibodies, able to exert inoculation selection. Better characterisation of the dynamics of mucosal immunity with age and across individuals is needed to understand the implications of inoculation selection to influenza virus evolution depending on the host population structure.

### 3 Detailed within-host model analysis

Here we analyze the within-host model in depth, and find expressions for within-host variant frequencies over time, the probability distribution for the time of first successful mutation to a new variant, and an approximate probability of replication selection to a given frequency by a given time.

#### 3.1 Expression for the within-host mutant frequency

In our minimal model, competition between new variant and old variant viruses obeys a simple replicator equation.

The within-host frequency of new variant virus is $f_m(t) = \frac{V_m(t)}{V_m(t)+V_w(t)} = \frac{V_m(t)}{V_{\text{tot}}(t)}$. Its rate of change is $\frac{\mathrm{d}f_m}{\mathrm{d}t}$. We observe that $\frac{\mathrm{d}V_i}{\mathrm{d}t}$ is of the form $\frac{\mathrm{d}V_i}{\mathrm{d}t} = V_i\Big[g_i(t) - d_i(t)\Big]$ for both the new variant and the old variant, where, neglecting mutation, $g_i(t) = r_i\beta C(t)$ and $d_i(t)$ is a (possibly time-varying) neutralization or decay rate. Differentiating $f_m(t)$ with respect to time demonstrates that the frequency of the new variant over time is governed by the following replicator equation (see section 9 for an explicit derivation):

$$\frac{\mathrm{d}f_m}{\mathrm{d}t} = f_m(1 - f_m)\Big(\big[g_m(t) - g_w(t)\big] - \big[d_m(t) - d_w(t)\big]\Big) \tag{A1}$$

If the new variant is neutral in the absence of antibody-mediated selection, $r_w = r_m = r$, and therefore $g_m(t) = g_w(t)$. It follows that despite changing target cell populations and virion population sizes, the variants compete in a manner independent of $V_{\mathrm{tot}}$, provided each $d_i(t)$ is independent of $V_{\mathrm{tot}}$. We have competition only through the respective death rates of old variant and new variant virions. These may be affected by antibody-mediated neutralization of virions.

If the fitness difference $\delta(t) = \Big[g_w(t) - g_m(t)\Big] - \Big[d_m(t) - d_w(t)\Big]$ is a constant $\delta$, this differential equation has a closed-form solution:

$$f_m(t) = \frac{e^{\delta t}}{e^{\delta t} + f_0^{-1} - 1} \tag{A2}$$

where $f_0 = f_m(0)$

If $\delta = 0$, $f_m(t)$ is constant and equal to $f_0$, as expected. If $\delta(t)$ is piecewise-constant, we can apply the constant-$\delta$ solution iteratively to find the mutant frequency.

It is therefore straightforward to apply this expression to the within-host model with binary immunity by evaluating different immunity regimes piece-wise.

We consider the case of a host who is experienced to the old variant ($E_w = 1$) but naive to the new variant ($E_m = 0$).

If first appears at a time $t_e \geq 0$ post-infection at an initial frequency $f_m(t_e)$, then if $t_e < t_M$:

$$f_m(t) = \begin{cases} 0 & t < t_e \\ f_m(t_e) & t_e \leq t < t_M \\ \frac{e^{\delta(t-t_M)}}{e^{\delta(t-t_M)} + f_m(t_e)^{-1} - 1} & t_M \leq t \end{cases} \tag{A3}$$

if $t_e > t_M$:

$$f_m(t) = \begin{cases} 0 & t < t_e \\ \frac{e^{\delta(t-t_e)}}{e^{\delta(t-t_e)} + f_m(t_e)^{-1} - 1} & t_e \leq t \end{cases} \tag{A4}$$

where $\delta = k(c_w - c_m)$ is the fitness difference between the variants due to the recall adaptive response.

#### 3.2 Mutant frequency over time given ongoing *de novo* mutation

Thus far we have neglected mutation the first mutation of interest (i.e. the first that produces a new variant lineage that evades stochastic extinction). In fact, with symmetric mutation between the two types of interest, we have:

$$\begin{aligned}\frac{\mathrm{d}V_w}{\mathrm{d}t} &= V_w\Big[(1-\mu)g_w(t) - d_w(t)\Big] + \mu g_m(t)V_m \\ \frac{\mathrm{d}V_m}{\mathrm{d}t} &= V_m\Big[(1-\mu)g_m(t) - d_m(t)\Big] + \mu g_w(t)V_w\end{aligned} \tag{A5}$$

The rate of fitness-irrelevant mutation is assumed to be equal for the two types, and to preserve antigenic type ($w$ or $m$). The rate of lethal or deleterious mutants is assumed equal for the two types and is captured in $g$.

It can be shown (see section 9) that the equation for $\frac{\mathrm{d}f_m}{\mathrm{d}t}$ in this case is:

$$\frac{\mathrm{d}f_m}{\mathrm{d}t} = f_m(1-f_m)(\alpha_m - \alpha_w) + \mu\Big(g_w(t) - 2g_w(t)f_m + g_w(t)f_m^2 - g_m(t)f_m^2\Big) \tag{A6}$$

When $g_w(t) = g_m(t) = g(t)$:

$$\frac{\mathrm{d}f_m}{\mathrm{d}t} = f_m(1-f_m)\Big(d_m(t) - d_w(t)\Big) + \mu g(t)(1-2f_m) \tag{A7}$$

When $f_m \ll 0.5$, the net effect of symmetric mutation on $f_m$ is to increase it at rate $\approx \mu$ (note that the symmetric mutation term becomes zero at $f_m = 1/2$). In other words, it is roughly equivalent to a case with no back mutation at all:

$$\begin{aligned}\frac{\mathrm{d}V_w}{\mathrm{d}t} &= V_w\Big[(1-\mu)g_w(t) - d_w(t)\Big] \\ \frac{\mathrm{d}V_m}{\mathrm{d}t} &= V_m\Big[g_m(t) - d_m(t)\Big] + \mu g_w(t)V_w\end{aligned} \tag{A8}$$

in that case, we have:

$$\frac{\mathrm{d}f_m}{\mathrm{d}t} = f_m(1-f_m)\Big(\big[g_m(t)-(1-\mu)g_w(t)\big] - \big[d_m(t)-d_w(t)\big]\Big) + \mu g_w(t)(1-2f_m+f_m^2) \tag{A9}$$

In either case, we mainly study a situation in which $g_w(t) = g_m(t) = g(t)$ at all times (though note that $g(t)$ varies in time), $d_w(t) = d_m(t) = d(t) = d_v$ prior to the onset of the adaptive immune response, and $d_w(t) = d_v + k$, $d_m(t) = d_v + ck$ after the onset of the adaptive immune response.

If $f_m$ is large relative to $\mu$, the mutant growth term $V_m g(t)$ is large relative to the *de novo* mutation term $\mu g(t)V_w$. We can see this from the fact that $V_m = f_m V_{\text{tot}} \geq f_m V_w$, with equality if and only if $f_m = 0$. It follows that if $f_m > \mu > 0$, then:

$$g(t)V_m > g(t)f_m V_w > g(t)\mu V_w$$

But if the first mutation to produce a new variant is sufficiently late and there is little or no positive selection, we do not necessarily have $f_m \gg \mu$, so ongoing mutation cannot be ignored in the dynamics of $f_m$. In that case, we can consider either the $\mu g(t)(1 - 2f_m + f_m^2)$ term in equation A9 or the $\mu g(t)(1 - 2f_m)$ term in equation A7. We can approximate $g(t) \approx g_0 = g(0) = \mathcal{R}_0 d_v$ early in infection, since for small $t$, $C(t) \approx C_{\max}$ and therefore $g(t) = r\beta C(t) \approx g_0$.

With this approximation for $g(t)$, the one-way mutation case has an analytical solution for $\delta \neq 0$:

$$\begin{aligned} f_m(t) \approx & \quad \frac{A\exp(\delta t) + \mu g_0}{A\exp(\delta t) + \mu g_0 - \delta} \\ A = & \quad \frac{f_0}{1 - f_0}(\mu g_0 - \delta) - \frac{\mu g_0}{f_0} \end{aligned} \tag{A10}$$

And when $\delta = 0$:

$$f_m(t) \approx \frac{\frac{1}{1-f_0} + \mu g_0 t - 1}{\frac{1}{1-f_0} + \mu g_0 t} \tag{A11}$$

#### 3.3 When ongoing mutation matters

$f_m$ must be small for ongoing mutation to matter; it ceases to matter when $f_m \gg \mu g(t)$, and $\mu g(t)$ is small by assumption. It follows that in the absence of a fitness difference between the types:

$$\frac{\mathrm{d}f_m}{\mathrm{d}t} \approx \mu g_0 \tag{A12}$$

and therefore:

$$f_m(t) \approx f_0 + \mu g_0 t \tag{A13}$$

We can also see this by inspecting the other two expressions for $f_m(t)$ and seeing that they are quasi-linear for small $\mu$ and small $t$. This further confirms that it is not crucial to decide whether symmetric or one-way mutation is more realistic, as they will be functionally the same from the point of view of practical mutant frequencies in the absence of positive selection.

These approximations break down once $C(t) \ll C_{\max}$ and therefore $g_w(t) \ll g_0$; fewer wild-type replications are occurring, and rate of ongoing mutation therefore falls. So a reasonable approximation for the maximum value of $f_m(t)$ in the absence of positive selection is given by $f_m(t_{\text{peak}})$. For a mutation rate of $0.33 \times 10^{-5}$, typical values of $f_m(t_{\text{peak}})$ without selection range from $10^{-5}$ to $2 \times 10^{-4}$, depending on $\mathcal{R}_0$ and $t_{\text{peak}}$.

We use the expression in A10 to plot the analytical predicted mutant frequencies show in Fig. 1 and to calculate the transmission frequencies for the analytical model of observed phenotypes in 5.

##### 3.4 Consequences of ongoing *de novo* mutation

Ongoing mutation enhances possibilities for both replication and inoculation selection. It truncates the left tail of the distribution of $f_m(t_M)$ and the distribution of $f_{\text{mt}}$(the mutant frequency at the time of transmission). Whenever $t_M$ is sufficiently late that we have reached $V_w \ll \frac{1}{\mu}$ before $t_M$, $f_m(t_M)$ should be on the order of the inverse mutation rate, or larger.

The consequence is that probabilities of replication selection and inoculation selection should both be somewhat higher than those estimated based on the time to the first non-extinct *de novo* mutant lineage, as in Fig 1. This further strengthens the case for the importance of an adaptive response at 48 hours or later in explaining the absence of replication selection.

##### 3.5 Replication selection is helped when individual infected cells are more productive

For a fixed $\mathcal{R}_0$, replication selection becomes more likely if the virions produced per infected cell $r$ becomes large—that is, if every infected cell produces more individual virions.

For a given $d_v$, and $f$, a virus can achieve a large $\mathcal{R}_0$ either by having a high cell productivity $r$ or by having a high cell infection rate $\beta$.

The expected number of virus replications per lost target cell is given by $\frac{\dot{V}^+}{\dot{C}^-} = \frac{r\beta C\bar{V}}{f\beta C\bar{V}} = \frac{r}{f}$. The infection peaks and begins to decline when $\frac{\mathrm{d}V}{\mathrm{d}t} = 0$, which occurs when $g(t) = d(t)$, or $\frac{C_{\max}}{C} = \mathcal{R}_0$, provided $d(t)$ is fixed. It follows that $C_{\text{peak}} = \frac{C_{\max}}{\mathcal{R}_0}$. So the virus peaks once $C_{\max}\left(1 - \frac{1}{\mathcal{R}_0}\right)$ target cells have been consumed (assuming negligible target cell replenishment during the timecourse of infection), and $\frac{r}{f}C_{\max}\left(1 - \frac{1}{\mathcal{R}_0}\right)$ viral replication events have occurred (this is a slight overestimate due to target cell depletion).

Fixing $d_v$ and $\mathcal{R}_0$, as $r \to \infty$, $\beta \to 0$ and the number of viral replication events before peak viral load goes to $\infty$. Since each generated mutant survives stochastic extinction roughly proportional to $1/\mathcal{R}_0$, these earlier mutants also are lost stochastically less frequently. In other words, a mutant that survives stochastic extinction is generated with certainty well before the infection peaks and can spend arbitrarily long under selection. Conversely, as $r \to 0$, $\beta \to \infty$, the number of replication events that occur before the infection peaks goes to 0, and replication selection becomes impossible.

A virus of a given $\mathcal{R}_0$ that takes a shotgun approach of producing many not-especially-infectious virions is more likely to result in the proliferation of a mutant of interest under replication selection than one that produces a small number of highly infectious virions.

In other words, the relatively large infected cell productivity of influenza viruses—perhaps as large as hundreds or thousands of viable virions per cell (Frensing et al., 2016)—makes potential replication selection more efficient. This in turn makes the absence of observed replication selection harder to explain unless another mechanism can be invoked, such as inoculation selection with replication selection only occurring 2–3 days post-infection.

##### 3.6 Replication selection in the presence of sterilizing immunity

One special case bears mentioning, because it corresponds to multiple existing models (Kennedy & Read, 2017; Luo et al., 2012; Volkov et al., 2010) and may be relevant for other systems. If there is an immediate recall response ($t_M = 0$), a founding population of size $b$, and $k$ is large enough such that $\mathcal{R}^w(t = 0) < 1$ (the virus population is initially declining, in expectation), then the infection will go extinct after generating $q$ new virions for some finite $q$.

Each virions has a probability $\mu$ of being an escape mutant with $\mathcal{R}^m(0) > 1$ and each mutant independently survives stochastic extinction with probability $p_{\text{sse}} = (1 - 1/\mathcal{R}^m(0))$ (per the theory of supercritical branching processes). The conditional probability of replication selection given $q$ is therefore:

$$p_{\text{repl}}(q) = 1 - \Big(1 - \mu p_{\text{sse}})\Big)^q \tag{A14}$$

if $\mu p_{\text{sse}}$ is small, this is approximately equal to:

$$p_{\text{repl}}(q) \approx 1 - e^{-q\mu p_{\text{sse}}} \approx q\mu p_{\text{sse}} \tag{A15}$$

The approximations use the Poisson approximation to the binomial and the approximation $e^x \approx 1 + x$ when $|x| \ll 1$.

The unconditional probability of replication selection $p_{\text{repl}}$ is the expectation in $q$ of $p_{\text{repl}}(q)$: $E_q(p_{\text{repl}})$. But since $p_{\text{repl}}$ is approximately linear in $q$, the linearity of expectation implies:

$$p_{\text{repl}} \approx E_q(q\mu p_{\text{sse}}) = \bar{q}\mu p_{\text{sse}}$$

where $\bar{q}$ is the expected value of $q$.

Given $\mathcal{R}^w(0) < 1$, branching process theory predicts that the virus will produce on average $\bar{q} = \frac{b}{1-\mathcal{R}^w(0)} - b$ new copies before the infection is cleared (expected total length of $b$ independent subcritical branching processes, each with expected length $\frac{1}{1-\mathcal{R}^w(0)}$, minus the initial virions present (Ball et al., 2016)). This can be simplified to $\bar{q} = \frac{b\mathcal{R}^w(0)}{1-\mathcal{R}^w(0)}$.

So we have

$$p_{\text{repl}} \approx \bar{q}\mu p_{\text{sse}} = \frac{\mathcal{R}^w(0)}{1 - \mathcal{R}^w(0)} b\mu p_{\text{sse}} \tag{A16}$$

Note that (Iwasa et al., 2004) have previously derived this approximate result for the probability that a subcritical replicator mutates to a supercritical one prior to going extinct in the context of analyzing tumor cell mutations. They use a more rigorous generating function approach.

### 4 Point of transmission analysis

#### 4.1 Distinction between selection and drift at the point of transmission

In our model, mucosal antibodies acting at the point of transmission provide a mechanism for population level selection pressure: some protection of experienced hosts against infection with old variant virus, and worse protection against infection with new variant virus. In particular, it provides a mechanism that produces selection pressure for new antigenic variants while predicting that reinfections with old variant viruses—without observable new variant viruses—should still be observed. Prior models (see section 7) predict that in fully immune hosts, any observable reinfections will have new antigenic variant viruses at consensus.

Antibodies at the point of transmission appear to play a key role in population-level influenza virus evolution: they produce the selection pressure that allows new variants to spread more reliably and thus rapidly from host to host than old variants. But they may play a second role as well: promoting of new variants in frequency from the low frequencies at which they are typically generated to high frequencies at which they can be observed and reliably transmitted onward. This inoculation selection takes on the role of previously held by replication selection in explaining why inoculations of experienced hosts might produce new variant infections at a higher rate (per inoculation) than inoculations of naive hosts.

#### 4.2 The inoculation selection paradigm

As noted in the main text, new variants can reach high within-host frequencies through founder effects. These events are both possible and rare due to influenza's tight transmission bottleneck (McCrone et al., 2018; Xue & Bloom, 2019).

Whether this process is purely neutral or stochastically selective depends on whether the recipient host is naive, and if not how well they neutralize old variant versus new variant virions.

As mentioned in the Methods, in a fully naive host with large $v$ and small transmitting host new variant frequency, the sampling process approximates a low-probability binomial or a low frequency Poisson:

$$p_{\text{drift}} \approx 1 - e^{-f_{\text{mt}} b} \tag{A17}$$

When $f_{\text{mt}} \ll 1$, this is approximately equal to $b f_{\text{mt}}$

As noted in the main text , mucosal antibody neutralization can reduce the variant's survival probability relative to this neutral case (inoculation pruning) or increase it (inoculation promotion), depending upon parameters.

There can be inoculation pruning even when the new variant is more fit than the old variant (i.e. neutralized with lower probability). But sufficient subsequent bottlenecking after sIgA neutralization can create an inoculation promotion effect.

Whether inoculation selection produces pruning or promotion depends on the ratio of $v/b$, and on the mutant's probability of avoiding neutralization $1 - \kappa_m$ (main text Figs. 3F, 4, Fig. A1, and sec. 4.5 below).

##### 4.3 Expression for and properties of the new variant survival probability

As shown in the Methods, the probability that a new variant survives the cell infection bottleneck of size $b$ is well approximated for small $f_{\text{mt}}$ by:

$$p_{\text{surv}}(\kappa_m, \kappa_w, v, b, f_{\text{mt}}) \approx p_{\text{inoc}}(\kappa_m, v, f_{\text{mt}}) * p_{\text{cib}}(\kappa_w, v, b) \tag{A18}$$

$p_{\text{inoc}}$ is the probability that at least one mutant survives the sIgA bottleneck:

$$p_{\text{inoc}}(\kappa_m, v, f_{\text{mt}}) = 1 - e^{-v f_{\text{mt}}(1-\kappa_m)} \tag{A19}$$

Note that when $f_{\text{mt}} \ll 1, p_{\text{inoc}} \approx v f_{\text{mt}}(1 - \kappa_m)$

$p_{\text{cib}}(\kappa_w, v, b)$ is the probability that a mutant that survives the sIgA bottleneck is not lost at the final bottleneck (as given in equation 35 above):

$$p_{\text{cib}}(\kappa_w, v, b) = (1 - e^{-\bar{w}})\frac{b}{\bar{w}} + e^{-\bar{w}} \sum_{j=0}^{b-1} \frac{\bar{w}^j}{j!}\left(1 - \frac{b}{j+1}\right)$$

where $\bar{w} = v(1 - f_{\text{mt}})(1 - \kappa_w)$. If $b = 1$, this expression reduces to the first term.

###### 4.3.1 Properties of these probabilities

These probabilities have several intuitive and useful properties:

- $p_{\text{cib}} < 1$ if $\kappa_w < 1$ and $f_{\text{mt}} < 1$, and $p_{\text{cib}} = 1$ if $\kappa_w = 1$ or $f_{\text{mt}} = 1$. That is, surviving the cell infection bottleneck is certain only if there is no competition. See section 9.6 for derivation.

- The correction term in $p_{\text{cib}}$, $e^{-\bar{w}} \sum_{j=0}^{b-1} \frac{\bar{w}^j}{j!}\left(1 - \frac{b}{j+1}\right)$, is always negative. Each term except the last in the summation is negative, since $b > j + 1$ for $j < b - 1$. The last term is zero (the last term is included for notational reasons, so that the formula is correct with $b = 1$). This implies that the first term is always $\geq p_{\text{cib}}$, with equality if and only f $b = 1$.

- If $\kappa_m = \sigma \kappa_w$, then $p_{\text{inoc}}$ is decreasing in $\kappa_w$ for $0 < \sigma \leq 1$. This is as expected: the more likely the new variant is to be neutralized, the smaller $p_{\text{inoc}}$ becomes. It suffices to show that $\frac{\mathrm{d}p_{\text{inoc}}}{\mathrm{d}\kappa_w} < 0$ for $0 < \sigma \leq 1$. We find:

$$\frac{\mathrm{d}p_{\text{inoc}}}{\mathrm{d}\kappa_w} = -e^{-v f_{\text{mt}}(1-\sigma\kappa_w)}(v f_{\text{mt}} \sigma) \tag{A20}$$

  Since $f_{\text{mt}}, v > 0$, this is negative whenever $\sigma > 0$, and 0 if $\sigma$ is 0.

It is sometimes useful to write this as:

$$\frac{\mathrm{d}p_{\text{inoc}}}{\mathrm{d}\kappa_w} = v f_{\text{mt}} \sigma (p_{\text{inoc}} - 1) \tag{A21}$$

- $p_{\text{cib}}$ is decreasing in $\bar{w}$ (see section 9.7 for derivation). This is intuitive: the less competition in the form of (expected numbers of) un-neutralized old variant virions, the more likely a surviving mutant is to pass through the cell infection bottleneck.

- $p_{\text{cib}}$ is increasing in $\kappa_w$ and $f_{\text{mt}}$ and decreasing in $v$. Since $v$, $1 - \kappa_w$, and $1 - f_{\text{mt}}$ are all $\geq 0$, it is clear that $\bar{w}$ is decreasing in $\kappa_w$ and $f_{\text{mt}}$ and increasing in $v$. It follows that $p_{\text{cib}}$ is increasing in $\kappa_w$ and $f_{\text{mt}}$ and decreasing in $v$. In all cases, this reflects the parameters' effect on the mean amount of competition for the mutant from old variant virions for the cell infection bottleneck.

- $p_{\text{inoc}}$ is increasing in $f_{\text{mt}}$, since a larger $f_{\text{mt}}$ implies an $v f_{\text{mt}}(1 - \kappa_m)$ term that is larger in absolute value, so $e^{-v f_{\text{mt}}(1-\kappa_m)}$ becomes smaller and $p_{\text{inoc}} = 1 - e^{-v f_{\text{mt}}(1-\kappa_m)}$ becomes larger.

#### 4.4 Mutant survival probability increases as transmitted mutant frequency increases

For given values of the other parameters, larger $f_{\text{mt}}$ implies larger $p_{\text{surv}}$. Since both $p_{\text{inoc}}$ and $p_{\text{cib}}$ are increasing in $f_{\text{mt}}$ and are both always positive, it follows that $p_{\text{surv}} = p_{\text{inoc}} p_{\text{cib}}$ is also increasing in $f_{\text{mt}}$. This makes intuitive sense: all else equal, higher frequency mutants always have a better chance of surviving at the point of transmission, regardless of the effects of the sIgA bottleneck.

#### 4.5 The relative importance of selection and drift at the point of inoculation

Consider $p_{\text{surv}}/\ p_{\text{drift}}$. This ratio quantifies the degree to which sIgA neutralization at the point of transmission facilitates (or hinders) mutant survival of the final bottleneck relative to a purely neutral founder effect.

For realistically small $b$ and $f_{\text{mt}}$, we can use the approximations $p_{\text{drift}} \approx b f_{\text{mt}}$ and $p_{\text{inoc}} \approx v f_{\text{mt}}(1 - \kappa_m)$

Then we have

$$p_{\text{surv}}/p_{\text{drift}} = \frac{p_{\text{inoc}} p_{\text{cib}}}{p_{\text{drift}}} \approx \frac{v}{b}(1 - \kappa_m) p_{\text{cib}}(\kappa_w, v, b) \tag{A22}$$

Several intuitive results follow:

- If we hold $\kappa_w$ constant, increasing $\kappa_m$ makes inoculation selection less effective relative to drift, because the inoculated new variant is at greater risk of neutralization.

- Conversely, if we hold $\kappa_m$ constant, increasing $\kappa_w$ makes inoculation selection more effective relative to drift (this follows from the fact that $p_{\text{cib}}$ is increasing in $\kappa_w$, see section 4.3.1 above).
- Since $p_{\text{cib}}$ is increasing in $f_{\text{mt}}$ (section 4.3.1), $p_{\text{surv}}/p_{\text{drift}}$ is increasing in $f_{\text{mt}}$. That is, inoculation selection is more efficient relative to drift when promoting higher frequency mutants, all else equal.
- $p_{\text{surv}}/p_{\text{drift}}$ is decreasing in $b$ (see section 9.9 for a derivation). In other words, when the cell infection bottleneck is wider, inoculation selection is less important relative to drift. The large bottleneck gives the mutant a reasonably good chance of surviving even in the absence of any neutralization of competing old variant virions.

For a bottleneck of size $b = 1$, we can also see that inoculation selection becomes more efficient relative to drift as $v$ increases, since:

$$\begin{aligned} p_{\text{surv}}/p_{\text{drift}} &\approx \frac{v}{b}(1-\kappa_m)p_{\text{cib}} \\ &= v(1-\kappa_m)\frac{1-\exp\big(-v(1-f_{\text{mt}})(1-\kappa_w)\big)}{v(1-f_{\text{mt}})(1-\kappa_w)} \\ &= \frac{1}{1-f_{\text{mt}}}\frac{1-\kappa_m}{1-\kappa_w}\big[1-\exp\big(-v(1-f_{\text{mt}})(1-\kappa_w)\big)\big] \end{aligned}$$

This is increasing in $v$.

#### 4.6 Strongly immune hosts with small bottlenecks provide intuition for Fig. 4

In a host strongly immune to the old variant (large $\kappa_w$), we have $-v(1-f_{\text{mt}})(1-\kappa_w) \ll 1$. Applying the linear approximation to the exponential to equation 4.5:

$$\begin{aligned} p_{\text{surv}}/p_{\text{drift}} \approx & \quad \frac{1}{1-f_{\text{mt}}}\frac{1-\kappa_m}{1-\kappa_w}(v(1-f_{\text{mt}})(1-\kappa_w)) \\ = & \quad (1-\kappa_m)v \end{aligned}$$

This provides intuition for the effects seen in main text Fig. 4, in which inoculation selection improves survival relative to drift by a factor of $\frac{v}{b}$ for a full escape mutant, and by a factor scaled down by $1-\kappa_m$ for a partial escape mutant.

Moreover, if we let $f_{\text{mt}} = f_d$ for the drift case and $f_{\text{mt}} = f_r$ for the selective case, the approximate ratio becomes

$$p_{\text{surv}}/p_{\text{drift}} \approx \frac{f_r}{f_d}(1-\kappa_m)v$$

Our replicator equation implies that this ratio $\frac{f_r}{f_d}$ will increase in expectation when more replication selection occurs prior to transmission (larger $\delta\tau$), explaining the increasing ratio of the selective variant survival relative to drift observed in Fig. 4 when replication selection is permitted prior to transmission.

Note that we did not integrate over the emergence times to obtain $p_{\text{nv}}$; instead, we derived principles that take the initial frequency as a given.

##### 4.7 Effect of sIgA cross immunity $\sigma$ on probability of inoculation selection

We also wish to know which hosts best promote antigenic novelty when inoculated, given a level of cross-immunity $\sigma$, or a degree of escape $1-\sigma$. We will show that whereas replication selection suggests that intermediately immune hosts should be key selectors for antigenic novelty (Grenfell et al., 2004), in an inoculation selection regime, new variants may most often reach observable frequencies in fully immune hosts. We do this by analyzing an expression for $p_{\text{surv}}$, the probability that a mutant survives transmission to found an infection in a new host, and showing how it changes with old variant neutralization probability $\kappa_w$ and sIgA cross immunity $\sigma = \kappa_m/\kappa_w$. We find that transmission survival probability can be maximized in strongly immune or fully naive hosts, depending on parameters.

We optimize $p_{\text{surv}}(\kappa_w)$ given some value of $\sigma$ by differentiating.

$$\begin{aligned}\frac{\mathrm{d}p_{\text{surv}}}{\mathrm{d}\kappa_w} &= \frac{\mathrm{d}p_{\text{cib}}}{\mathrm{d}\kappa_w}p_{\text{inoc}} + p_{\text{cib}}\frac{\mathrm{d}p_{\text{inoc}}}{\mathrm{d}\kappa_w} \\ &= \frac{\mathrm{d}p_{\text{cib}}}{\mathrm{d}\kappa_w}p_{\text{inoc}} + p_{\text{cib}}(f_{\text{mt}}\sigma)(p_{\text{inoc}}-1)\end{aligned} \tag{A23}$$

Consider the endpoint at $\kappa_w = 1$. $p_{\text{cib}} = 1$, $p_{\text{inoc}} \approx f_{\text{mt}}v(1-\kappa_m) = f_{\text{mt}}v(1-\sigma)$, and $\bar{w} = v(1-f_{\text{mt}})(1-\kappa_w) = 0$.

So $\frac{\mathrm{d}p_{\text{cib}}}{\mathrm{d}\kappa_w} = -v(1-f_{\text{mt}})e^{-\bar{w}}\left[S'(\bar{w}) - S(\bar{w})\right]$ can be evaluated, and it evaluates to 0 except if $b = 1$ (when it is undefined).

**Case 1: $b > 1$.** For $b > 1$, it follows from the above that

$$\frac{\mathrm{d}p_{\text{surv}}}{\mathrm{d}\kappa_w} = 0 - p_{\text{cib}}\frac{\mathrm{d}p_{\text{inoc}}}{\mathrm{d}\kappa_w} \tag{A24}$$

Since $\frac{\mathrm{d}p_{\text{inoc}}}{\mathrm{d}\kappa_w} \geq 0$, $\frac{\mathrm{d}p_{\text{surv}}}{\mathrm{d}\kappa_w} < 0$ unless $\frac{\mathrm{d}p_{\text{inoc}}}{\mathrm{d}\kappa_w} = 0$, which occurs in cases of interest when $\sigma = 0$ (complete escape mutant).

In other words, if immune escape is incomplete for $b > 1$, there is always some less strongly immune host (though possibly very slightly less, see Fig. A1) that improves new variant survival relative to a host who neutralizes old variant virions with 100% certainty. Note, however, that hosts with $\kappa_w = 1$ are very unlikely to exist in nature. Our study is motivated precisely by the fact that even an experienced encountering homotypic reinfection neutralizes virions with probability $\kappa_w < 1$, and therefore can be productively reinfected (Clements et al., 1986; McCrone et al., 2018). It follows that in practice the most strongly immune hosts (with $\kappa_w$ large but less than 1) could still be the best selectors.

**Case 2:** $b = 1$. When the bottleneck is 1, $\lim_{\bar{w}\to 0} \frac{\mathrm{d}p_{\mathrm{cib}}}{\mathrm{d}\bar{w}} = -\frac{1}{2}$. So in this case, the derivative at $\kappa_w = 1$ may be positive or negative, depending on whether:

$$\frac{\mathrm{d}p_{\mathrm{cib}}}{\mathrm{d}\kappa_w} p_{\mathrm{inoc}} + p_{\mathrm{cib}}(v f_{\mathrm{mt}} \sigma)(p_{\mathrm{inoc}} - 1) > 0 \tag{A25}$$

substituting $\frac{\mathrm{d}p_{\mathrm{cib}}}{\mathrm{d}\bar{w}} \frac{\mathrm{d}\bar{w}}{\mathrm{d}\kappa_w} = -\frac{1}{2}(-v(1 - f_{\mathrm{mt}}))$ and $p_{\mathrm{cib}} = 1$, we have:

$$\frac{1}{2}(1 - f_{\mathrm{mt}}) p_{\mathrm{inoc}} + f_{\mathrm{mt}} \sigma p_{\mathrm{inoc}} > f_{\mathrm{mt}} \sigma$$

$$\frac{1}{2 f_{\mathrm{mt}}}(1 - f_{\mathrm{mt}}) p_{\mathrm{inoc}} + \sigma p_{\mathrm{inoc}} > \sigma$$

$$\frac{1}{2 f_{\mathrm{mt}}}(1 - f_{\mathrm{mt}}) p_{\mathrm{inoc}} > \sigma(1 - p_{\mathrm{inoc}})$$

$$\frac{1 - f_{\mathrm{mt}}}{2 f_{\mathrm{mt}}} \frac{p_{\mathrm{inoc}}}{1 - p_{\mathrm{inoc}}} > \sigma$$

When $p_{\mathrm{inoc}}$ is small, we have approximately $p_{\mathrm{inoc}} \approx f_{\mathrm{mt}} v(1 - \sigma)$ and $1 - p_{\mathrm{inoc}} \approx 1$, so:

$$\frac{1 - f_{\mathrm{mt}}}{2} v(1 - \sigma) > c$$

$$v \frac{1 - f_{\mathrm{mt}}}{2} > \frac{\sigma}{1 - \sigma} \tag{A26}$$

In other words, with extremely small cell infection bottlenecks like those observed for influenza viruses, fully immune hosts are the best selectors provided that the number of virions $v$ encountering IgA antibodies is sufficiently large given the degree of escape achieved. Less immune escape (larger sIgA cross immunity $\sigma$) necessitates a larger $v$ to make fully immune hosts the best selectors. Since $f_{\mathrm{mt}} \ll 1$, a rule of thumb is that $v > \frac{2\sigma}{1-\sigma}$.

The intuition is that larger $v$ means more competition for the cell infection bottleneck among virions that reach IgA, and thus a greater opportunity for the mutant's selective advantage to be realized, but that this only works provided that this advantage is large enough so that the mutant is not itself at too large a risk of being neutralized.

### 5 Parameter uncertainties and sensitivity analysis

Here we discuss parameter uncertainty in our models and how they affect our conclusions. We also conduct a simulation-based sensitivity analysis of the central within-host model.

#### 5.1 Strength of selection

A crucial parameter in our model is $\delta$: the magnitude of the fitness advantage of the new variant over the old variant during viral replication in an infected individual. One possible objection to our analysis here is that the antibody mediated virion neutralization rate $k$ is low enough or the antigenic similarity between the variants is great enough to make the fitness difference $\delta = k(c_w - c_m)$ small. In that case, replication selection to consensus will be rare even if $t_M$ is very small, and the adaptive response is mounted immediately (main text Fig. 1C,D). Despite uncertainty about both $k$ and $c_w - c_m$, $k$ is likely to be high, perhaps extremely so. And given sufficiently high $k$ and a homotypically reinfected host ($c_w = 1$), even moderate immune escape ($0 \gg c_m < 1$) produces a substantial fitness difference. Moreover, assuming small $k$ early in infection grants our basic hypothesis: antibodies mediated selection is weak early in infection, neutralization during viral replication is not the mechanism of protection against reinfection, and there is asynchrony between antigenic diversity and meaningful antigenic selection.

##### 5.1.1 Antibody-mediated virion neutralization rate ($k$) may be very high

Parameter estimates for antibody-mediated virion neutralization in the presence of substantial well-matched antibody correspond to values of $k$ that are extremely high. For instance, by fitting a single-variant within-host model to data, Cao and McCaw (Cao & McCaw, 2017) estimate antibody neutralization rates between 0.4 and 0.8 per virion-day per pg/mL of antibody, and antibody concentrations of over 100 pg/mL by day 6 in an infection of a naive host. This corresponds to a $k$ of 40 to 80 in our more phenomenological model.

For $t_M = 0$ (constant immunity) if $k$ is sufficiently large that the old variant virus has an initial $\mathcal{R}(0) < 1$ (which occurs if $k > d_v(\mathcal{R}_0 - 1)$), then all visible reinfections will be mutant infections. At $\mathcal{R}_0 < 10$ with $d_v = 4$, this corresponds to a $k$ of 40, the lower end of the Cao and McCaw estimates.

A $k$ of this magnitude has several implications that support our hypotheses of asynchrony between diversity and selection. Such a $k$ would suffice to drive $\mathcal{R}(0)$ below 1. A sufficiently early activation of a recall antibody response would then imply that all observable infections of experienced hosts would be mutant infections (Fig. 2). In intermediately immune hosts, there could be a substantial fitness advantage $\delta = (c_w - c_m)k$ for new variant viruses over old variant viruses (where $c_m < c_w$ are the cross immunities to the host memory variant for the old variant and the new variant, respectively).

With large $k$ and $t_M = 0$, intermediately immune hosts who cannot block transmission could easily have $\mathcal{R}(0) \approx 1$ for the old variant; this would make them excellent at generating and selecting for mutant before an infection is cleared.

A final reason why a high $k$ supports the hypotheses of this paper is that antigenic evolution is extremely rapid if a sufficiently strong recall antibody responses becomes active after viral replication begins but before the infection peaks. New antigenic variants should be generated with near-certainty by virus exponential growth well before the infection's peak—roughly when the number replications that have occurred is at or above the order of magnitude of the inverse mutation rate (Fig. 8). Suppose a strong

($\mathcal{R}(t_e) < 1$) recall response is mounted at that point $t_e$. The mutant then has target cell resources on which to grow (since $C(t_e) \gg C_{\text{peak}}$) and experiences negligible replication competition from the old variant (since the old variant population is not growing but in fact is shrinking). If not lost stochastically, it should therefore emerge to detectable and transmissible levels. We do not observe this. This suggests that if $k$ is large, not only must the antibody response not be immediate ($t_M > 0$), but it must also be late enough enough to make this effect unlikely: it must happen either just before or after the peak of infection. Evidence suggests that this is indeed the case. Infections peak by 36–48 hours post-infection, and antibody responses only begin 48–72 hours post infection.

##### 5.1.2 Small values of $k$ are a sub-hypothesis of the general model of asynchrony between diversity and selection

Small $k$ early in infection does produce weak replication selection, but it means that neutralization during viral replication cannot explain protection against reinfection. In such a scenario, it is still necessary to invoke mucosal sIgA antibodies or another mechanism of protection at the point of transmission, and so inoculation selection again comes into play.

Indeed, it is unlikely that $k$ is truly zero before the adaptive response is mounted. Occasionally, a virion may encounter residual IgA antibodies, for instance (see section 2. But the effective value of $k$ is likely to be small—too small to curtail the growing infection or produce substantial replication selection. We set it to 0 before $t_M$ for simplicity of model analysis, but our simple binary model approximates the likely scenario in which $k$ is small but non-zero before $t_M$ and then increasingly large afterwards (see 5.1.1).

#### 5.2 Relationship between $k$ and $\kappa$

As noted in the Discussion, the relationship between the mucosal antibody neutralization rate $\kappa$ and the antibody neutralization rate during replication $k$ is unknown, though we have every reason to expect it to be positive.

$\kappa$ values are particularly difficult to calculate. In our model, we have generally assumed that all virions of type $i$ inoculated into an experienced host are independently neutralized with probability $\kappa_i$, and therefore $z_i = e^{-v(1-\kappa_i)}$ for an inoculum consisting only of type $i$. But this assumption of independence may be violated in practice.

One reason statistical independence may be violated is that antibody numbers are finite, and an antibody that binds to one virion cannot bind to another. Consider a focal virion. Given that another virion has been neutralized, there are fewer antibodies remaining to neutralize our focal virion. This is particularly important for mixed inocula. Old variant virions, which have higher affinity for the inoculated host's IgA antibodies, may indirectly protect the new variant by competing with it for antibody binding. A new variant virion may have a higher individual chance of being neutralized by those same antibodies if it is part of a monomorphic inoculum composed of other, identical new variants. Such an interaction would strengthen inoculation selection relative to the independent neutralization we have modeled here.

#### 5.3 Bottleneck sizes

Inoculation selection may improve mutant bottleneck survival relative to neutral drift (see Fig. 3F, Fig. A1, and section 4). For this to be true, a non-antigenic bottlenecking must follow IgA antibody neutralization, with a ratio $v/b \gg 1$

Evidence suggests that a non-antigenic bottleneck does occur after the sIgA bottleneck because bottlenecks measured in vaccinated and non-vaccinated hosts are of comparable size (indistinguishable from one), suggesting that founding virion population sizes are cut down to small numbers even in the absence of IgA antibodies, though this would also be consistent with the case of $v = 1$ (i.e. IgA antibodies must neutralize on average a single virion to prevent infection, and this virion, if not neutralized or otherwise lost, uniquely founds the infection).

An experimental evolution study of avian influenza virus adaptation in ferrets found that transmission bottlenecks in naive hosts became tighter that as the virus adapted (Moncla et al., 2016). A re-analysis of that data found that bottlenecks were tight throughout (Lumby et al., 2018). The fact that bottlenecks do not appear to loosen (and may tighten) with adaptation for better transmission and replication is further evidence, albeit circumstantial, that when an influenza virus is well-adapted to its host and replicates rapidly, the first virion or first few virions to infect a cell will be the ancestor of the vast majority of progeny viruses.

#### 5.4 Double-peaked infections

One modeling study of influenza viruses proposes that infections should have a second peak after initial innate responses are overcome and some target cells are once again susceptible to infection (Pawelek et al., 2012). A relaxation of non-antigenic limiting factors later in infection can and should provide additional opportunities for replication selection, as occurs in immune-comprised human patients.

That said, the empirical data suggesting double-peaked kinetics comes from experimental inoculation of naive horses with a large quantity of influenza virus: $10^6$ 50% egg infectious dose (Quinlivan et al., 2007). The novel antibody response curbed the second peak of infection, which was much lower than the first.

In human kinetics data (Hadjichrysanthou et al., 2016) and animal transmission experiments in which one animal infects another (Canini et al., 2020; Le Sage et al., 2020) it is common to see clearly single-peaked infections.

Furthermore, for realistic (incomplete) degrees of antibody binding escape, we expect both old and new antigenic variant population sizes to decline even due to *antigenic* limiting factors, though of course we expect the new variant to decline more slowly.

The upshot is that virus clearance should be rapid following the mounting of the adaptive response in the experienced hosts where we expect selection to take place, and thus in immune-competent hosts we should expect a limited window for antibody-mediated selection on transmissible virus populations.

#### 5.5 Threshold versus probabilistic transmission

In the simulation results shown in the main text (Figs. 1, 2, 3, 5) we use a probabilistic model of transmission: the donor host's probability of inoculating the recipient at a given point during the infection is proportional to donor viral load in virions, $V_{\mathrm{tot}}(t)$.

Since influenza virus population sizes grow and shrink rapidly around the within-host peak, however, this model should be qualitatively similar to a threshold model in which transmission occurs with certainty if the total viral load is above a certain threshold $\theta$ and does not occur if it is below that threshold. To confirm this, we here show corresponding simulation results based on a threshold model, with the threshold set equal to $V_{50}$ from the proportional model.

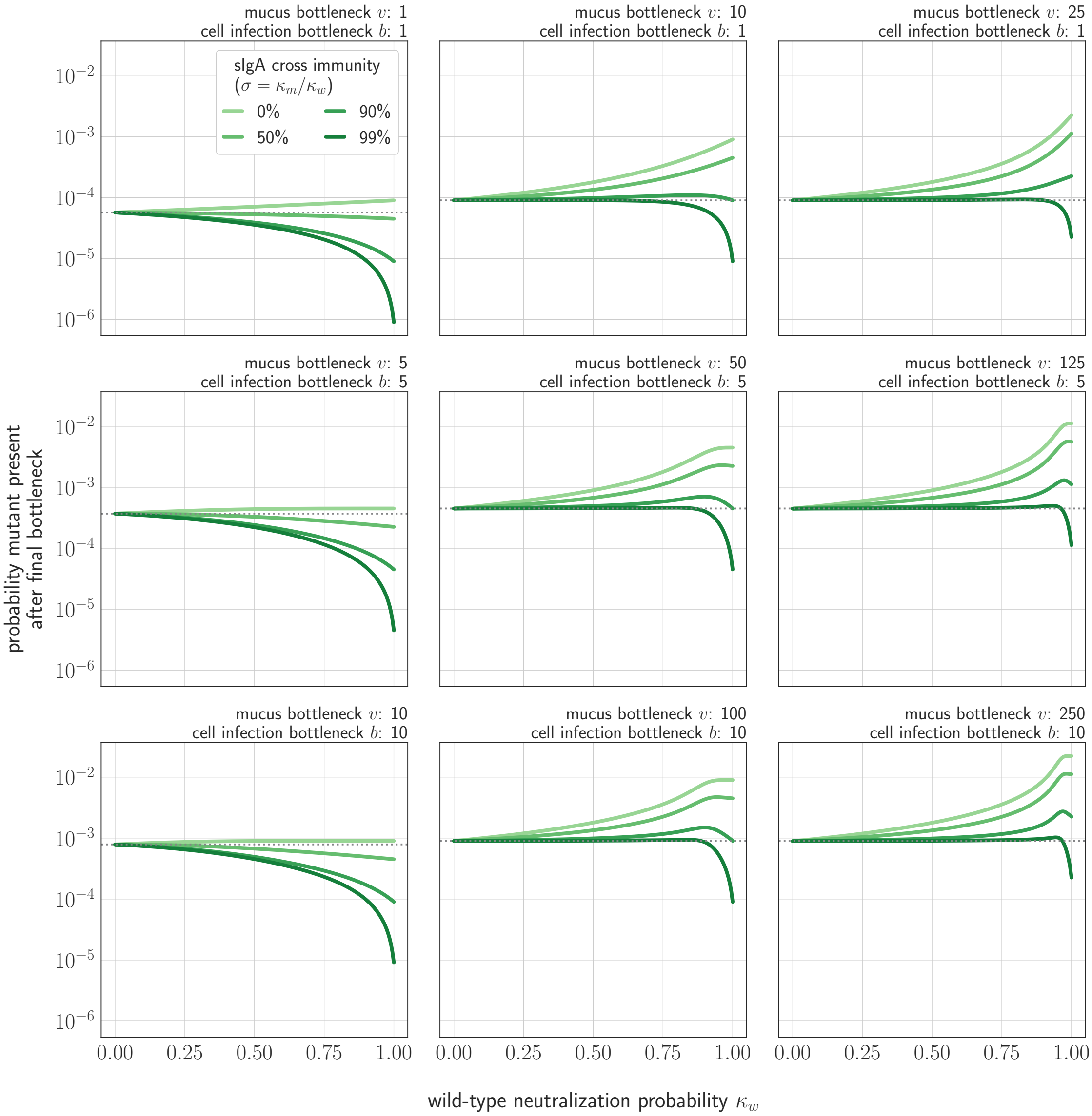


**Fig. A1. Probability that a new variant is present after cell infection (final) bottleneck as a function of cross immunity and degree of competition for the final bottleneck.** Probability shown as a function of probability of no old variant infection ($z_w$), degree of cross immunity between mutant and new variant $\sigma = \kappa_m/\kappa_w$, mucus bottleneck size $v$, and final bottleneck size $b$. Gray dotted line indicates probability that a new variant survives the transmission bottleneck in a host who is naive both to the old variant and to the new variant (i.e. drift). $f_{mt} = 9 \times 10^{-5}$, a typical value for a naive transmitting host in stochastic simulations

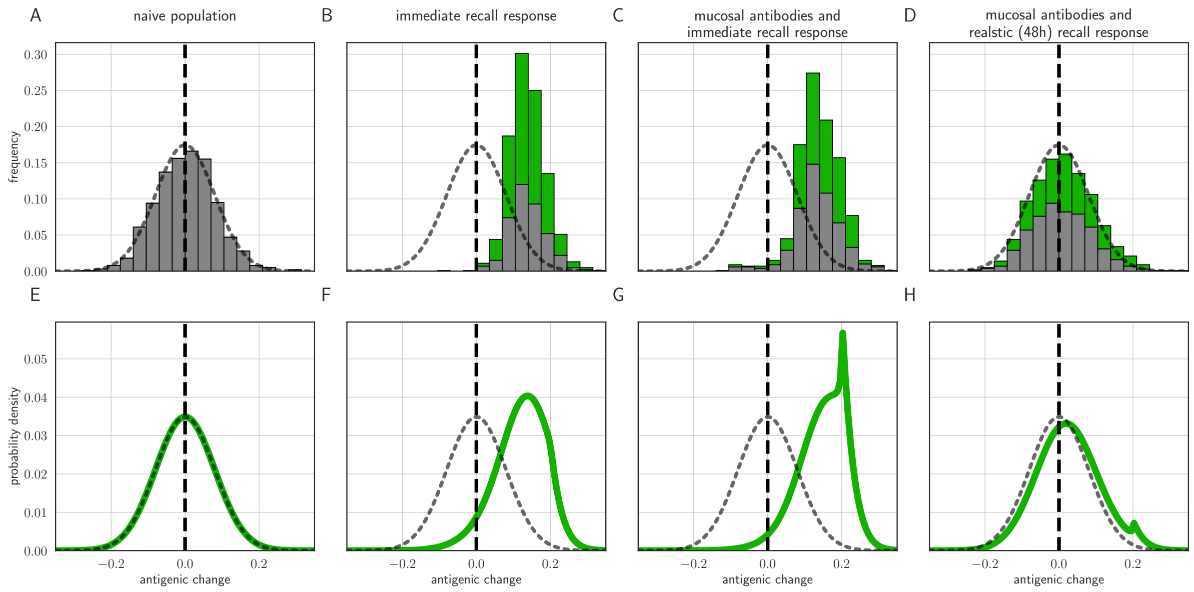


**Fig. A2. Distribution of mutant effects given replication and inoculation selection, with a transmission threshold model.** Threshold version of Fig. 5. Distribution of antigenic changes along 1000 simulated transmission chains (**A**–**D**) and from an analytical model (**E**–**H**). In (**A**,**E**) all naive hosts, in other panels a mix of naive hosts and experienced hosts. Antigenic phenotypes are numbers in a 1-dimensional antigenic space and govern both sIgA cross immunity, $\sigma$, and replication cross immunity, $c$. A distance of $\geq 1$ corresponds to no cross immunity between phenotypes and a distance of 0 to complete cross immunity. Gray line gives the shape of Gaussian within-host mutation kernel. Histograms show frequency distribution of observed antigenic change events and indicate whether the change took place in a naive (grey) or experienced (green) host. In (**B**–**D**) distribution of host immune histories is 20% of individuals previously exposed to phenotype $-0.8$, 20% to phenotype $-0.5$, 20% to phenotype 0 and the remaining 40% of hosts naive. In (**E**), a naive hosts inoculate naive hosts. In (**F**–**H**) a host with history $-0.8$ inoculate hosts with history $-0.8$. Initial variant has phenotype 0 in all sub-panels. Model parameters as in Table 1, except $k = 25$. Spikes in densities occur at 0.2 as this is the point of full escape in a host previously exposed to phenotype $-0.8$.

#### 5.6 Sensitivity analysis

As described in the Methods, we further assessed sensitivity of the within-host model to variation in key parameters by studying random parameter sets chosen from biologically plausible parameter ranges.

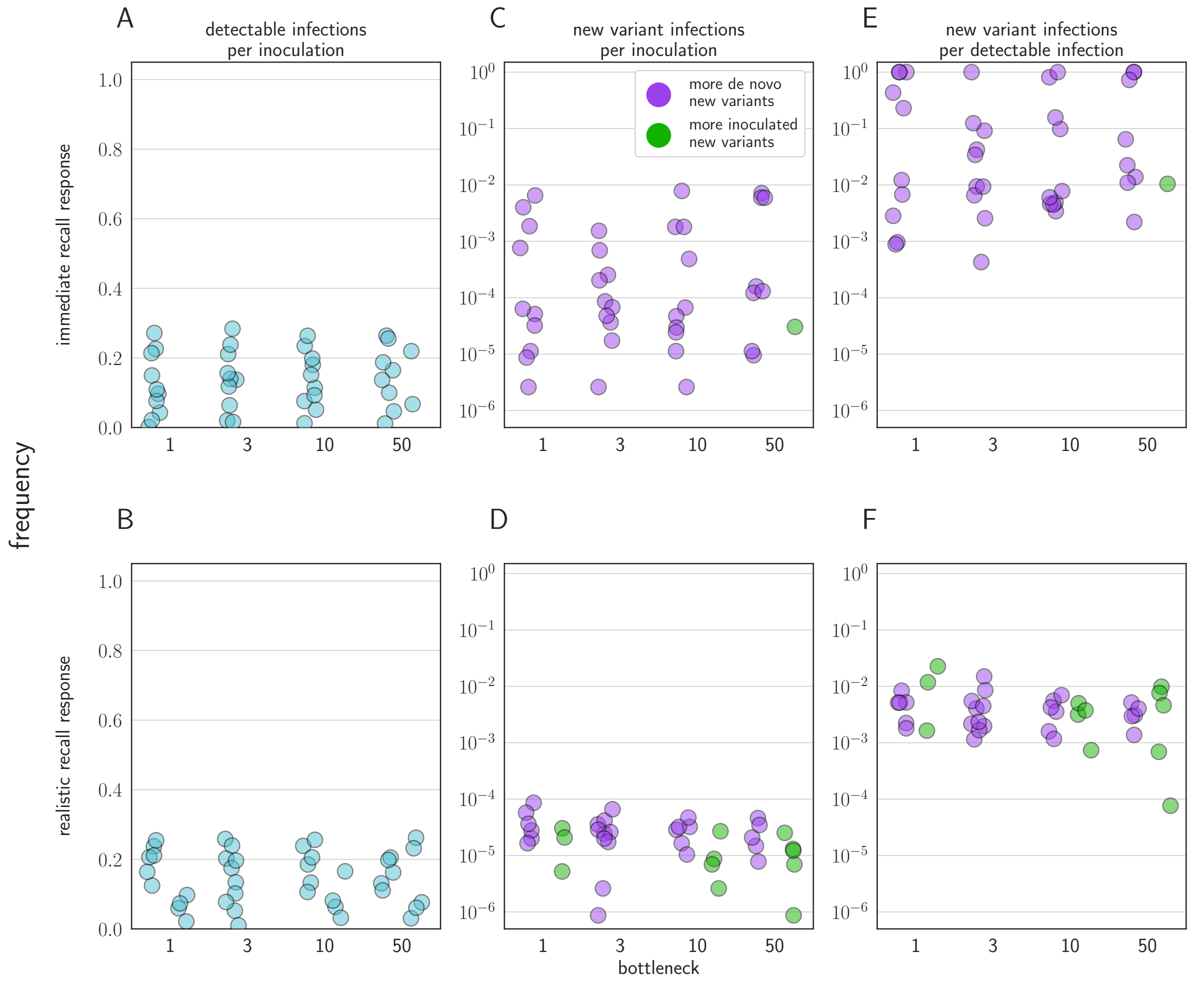


**Fig. A3. Sensitivity analysis varying model parameters across biologically-reasonable parameter ranges.** (**A, B**) Probability of a detectable infection per inoculation of an experienced host. (**C, D**) Probability of a detectable new variant infections per inoculation of an experienced host. (**E, F**) Fraction of detectable infections of experienced hosts that are new variant infections. Points colored according to whether new variant infections were more frequently caused by *de novo* generated (purple) or inoculated (green) new variant viruses. Each point represents a random parameter set; 10 random parameter sets generated for each bottleneck value shown, and 50 000 inoculations of an experienced host simulated for each parameter set. Two regimes of simulated parameter set shown: (**A, C, E**) an immediate recall response regime, in which $t_M$ varied from 0 to 1, and (**B, D, F**) a realistically timed response regime, in which $t_M$ varied from 2 to 4.5. All other parameters varied across the same ranges in both regimes (see Table 2 for ranges). Parameters Latin Hypercube sampled from within ranges for each regime and bottleneck size.

We analyzed two cases: one in which the immune response is unrealistically early ($0 \leq t_M \leq 1$, Fig. A4), and one in which it is realistically-timed ($2 \leq t_M \leq 4.5$) Fig. A5, see also Fig. A3 and other model parameters in table 2.

We found that with early immunity, regardless of particular parameter values, new variants are frequently seen when experienced hosts are detectably reinfected, and the overall probability of new variant infections is unrealistically high. These observable new variants are most often generated *de novo* and replication selected (Fig. A4, see also Fig. A3).

When immunity is realistically-timed, the pattern of much rarer replication selection is robust to variation in parameters. New variant infections compose a realistically small fraction of all detectable reinfections (Fig. A5, Fig. A3). Moreover, inoculation selection is sometimes more common than replication selection as a source of new antigenic variants (Fig. A5, see also Fig. A3).

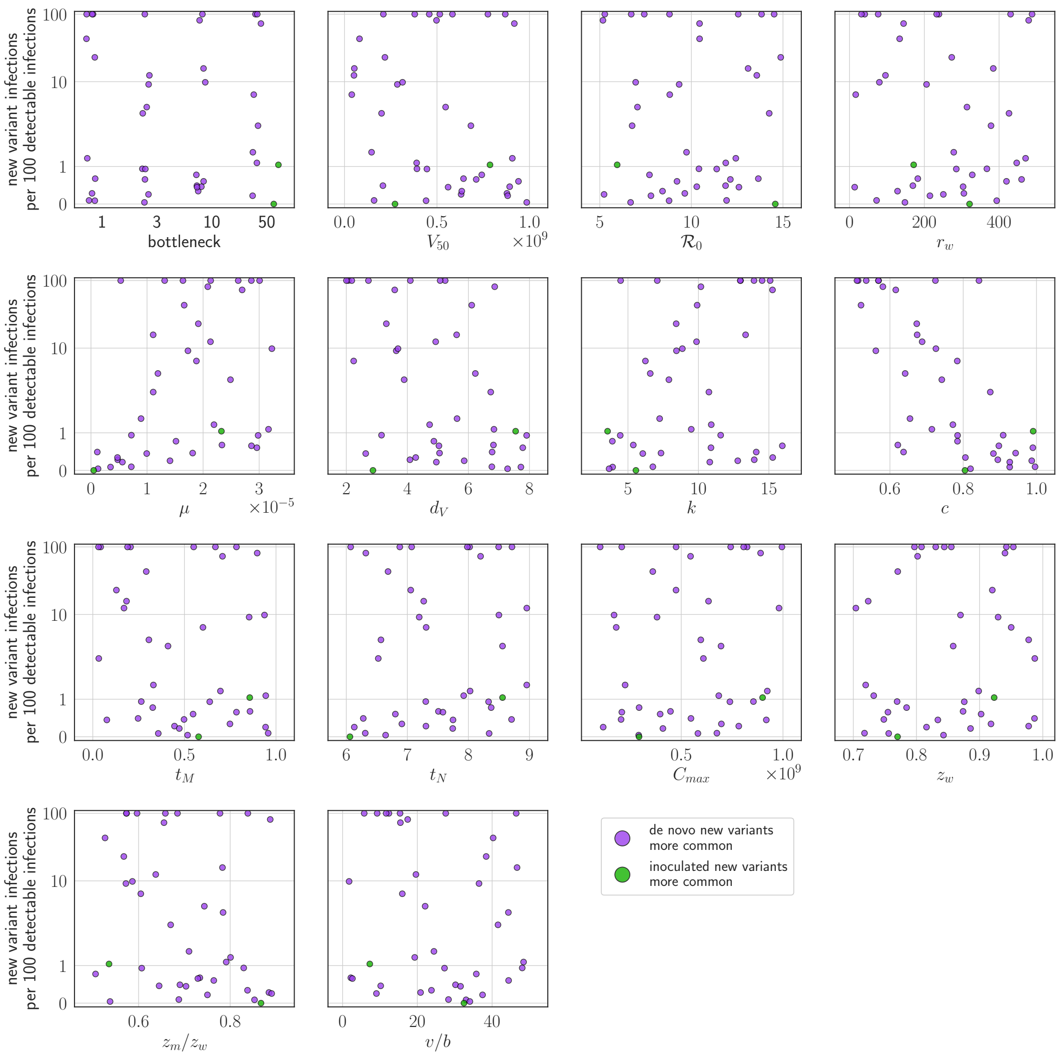


**Fig. A4. Sensitivity analysis: parameter values versus rate of new variant infections per 100 detectable infections of an experienced host, given an unrealistically early recall response.** Parameters randomly varied across the ranges given in table 2, with $t_M$ varied between 0 and 1. Each point represents a parameter set; the rate of new variant infections per hundred 100 detectable infections is estimated from 50 000 simulated inoculations of an experienced host. A new variant infection is defined as one in which the new variant reached a transmissible frequency of at least 1% at any point in the infection. Points are colored according to whether new variant infections were more frequently caused by *de novo* generated (purple) or inoculated (green) new variant viruses.

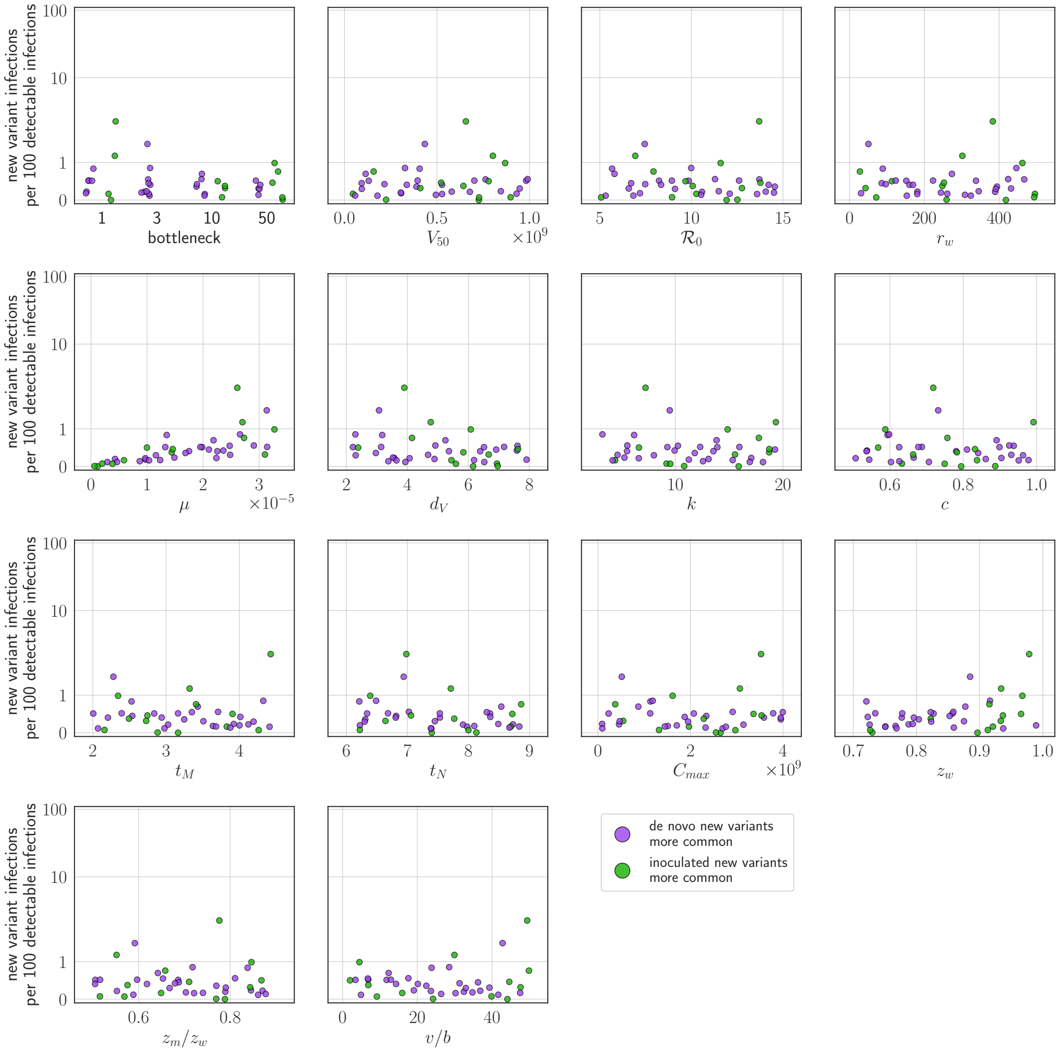


**Fig. A5. Sensitivity analysis: parameter values versus rate of new variant infections per 100 detectable infections of an experienced host given a realistic (48 hours or more post-infection) recall response.** Parameters randomly varied across the ranges given in table 2, with $t_M$ varied between 2 and 4.5. Each point represents a parameter set; the rate of new variant infections per hundred 100 detectable infections is estimated from 50 000 simulated inoculations of an experienced host. A new variant infection is defined as one in which the new variant reached a transmissible frequency of at least 1% at any point in the infection. Points are colored according to whether new variant infections were more frequently caused by *de novo* generated (purple) or inoculated (green) new variant viruses.

### 6 Polyphyletic antigenicity altering substitutions

#### 6.1 Background

Amino acid substitutions associated with antigenic "cluster transitions" have reached fixation in parallel—and near synchronously in time—in genetically distinct co-circulating lineages. This has occurred both in A/H1N1 and in A/H3N2 seasonal influenza viruses.

Existing "mutation-limited" or "diversity-limited" model explanations of why observable influenza virus antigenic novelty is rare despite continuous genetic evolution are less convincing in light of this synchronous polyphyly (see section 7). In particular, a number of models hypothesize that large effect antigenicity-altering substitutions can only emerge in specific genetic contexts (Gog, 2008; Koelle et al., 2006; Koelle & Rasmussen, 2015; Kucharski et al., 2015), and that this constraint limits the rate of population-level antigenic change.

Synchronous, polyphyletic cluster transitions cast doubt on these claims. If major antigenic changes appeared infrequently due to the rarity of "jackpot" scenarios of a large-effect mutant in a favorable genetic background (Koelle & Rasmussen, 2015), it would be surprising to see two or three distinct and synchronous jackpots after years of none. So while genetic background may play an important role in determining which lineages are fittest, synchronous polyphyly suggests that it is unlikely to set the clock of antigenic evolution. Rather, it suggests that accumulating population immunity may be crucial in setting the clock, and that there may even be a rising generation rate of observable antigenic novelty as a cluster ages, rather than a fixed rate (see section Population immunity sets the clock of antigenic evolution of the main text).

To show that large effect antigenic changes occur near-synchronously in multiple backgrounds, we assessed evolutionary relationships and built phylogenetic trees for known recent polyphyletic antigenicity-altering substitutions that have emerged in A/H1N1 and A/H3N2. Our analysis rules out reassortment as an explanation for the apparent polyphlyly, thus verifying that the branches represent distinct, independent, but simultaneous *de novo* lineages.

#### 6.2 Phylogenetic Methods

We analyzed polyphyletic antigenic changes using hemagglutinin (HA) gene nucleotide sequences deposited in the GISAID EpiFlu$^{TM}$ database.

We downloaded all seasonal A/H1N1 HA sequences from the period 1999–2008, to center on the antigenic cluster transition from the New Caledonia/1999-like phenotype to the Solomon Islands/2006-like phenotype, excluding pandemic A/H1N1 viruses. We downloaded A/H3N2 virus HA sequences for two periods: 2008 to 2011, to centre on cluster transition from Brisbane/07 to the Perth/2009-like and Victoria/2009-like phenotypic split, and 2012 to 2014, to capture the co-circulation of Clade 3C.2a and 3C.3a (table A1). We discarded all sequences with an incomplete HA1 domain or with more than 1% ambiguous nucleotides.

We aligned sequences using MAFFT v7.397 (Katoh et al., 2002) and reconstructed phylogenetic trees using RAxML 8.2.12 under the GTRGAMMA model (Stamatakis,

2014). We performed global optimization of branch length and topology on the RAxML reconstructed tree using Garli 2.01 with model parameters matching the RAxML reconstruction (500,000 generations) (Bazinet et al., 2014). It was computationally intractable to run Garli on the full phylogeny of A/H3N2 2012–2014 (n = 10107). We visualized phylogenetic trees using Figtree (http://tree.bio.ed.ac.uk/software/figtree) and ggtree (Yu et al., 2017)

We mapped amino acid substitutions onto branches using custom Python scripts available in the project Github. All numbering complies with H1/H3 numbering scheme (Burke & Smith, 2014).

Table A1: **Dataset composition**

| A/H1N1 1999–2008 | | A/H3N2 2008–2011 | | A/H3N2 2012–2014 | |
|---|---|---|---|---|---|
| Pre-filter | Final dataset | Pre-filter | Final dataset | Pre-filter | Final dataset |
| 4882 | 3514 | 7050 | 5738 | 11970 | 10107 |

## 6.3 A/H1N1

The HA K140E antigenicity-altering substitution first occurred in 2000 and was detected sporadically prior to its independent fixation in 2006–2007 in three phylogenetically distinct, geographically segregated lineages of A/H1N1 that diverged in 2004 (Fig. A6) (Bedford et al., 2015). The K140E substitution resulted in a cluster transition from the New Caledonia/1999-like antigenic phenotype to the Solomon Islands/2006-like phenotype, with lineage A emerging as the dominant lineage globally before the 2009 H1N1 pandemic (Bedford et al., 2015). Position 140 is located in the Ca2 antigenic site of the HA1 domain immediately adjacent to the receptor binding site, where amino acid composition mediates receptor binding function (Koel et al., 2013).

The three lineages have distinct HA1 mutational trajectories away from their most recent common ancestor, as defined by the set of amino acid substitutions that accumulate along the predominant trunk lineages both prior to and following the fixation of the K140E substitution (table A2, Fig. A6). All three lineages acquired amino acid substitutions at previously characterized antigenic sites, as well as substitutions with suggested functional consequences, including glycosylation gain and loss (Caton et al., 1982). The three lineages share the Y94H substitution prior to K140E fixation, with limited overlap of other substitutions: lineage A and B share changes at residue 188, whereas A189T occurs in lineage B and A prior to and post K140E emergence respectively.

Table A2: **Substitutions that characterize the co-circulating K140E-defined lineages**

| Lineage | Geographic composition in first year of co-circulation | Trunk substitutions from MRCA to K140E fixation | Trunk substitutions post-K140E fixation |
|---|---|---|---|
| 1 | South Asia | Y94H, R188K, E273K | D35N, K145R*, A189T, G185V, N183S, G185S |
| 2 | East Asia | Y94H, S36N, A189T, R188M, T193K* | N244S, K82R*, I47K, E68G |
| 3 | South-East Asia | Y94H, K73R, V128A*, A128T* | P270S |

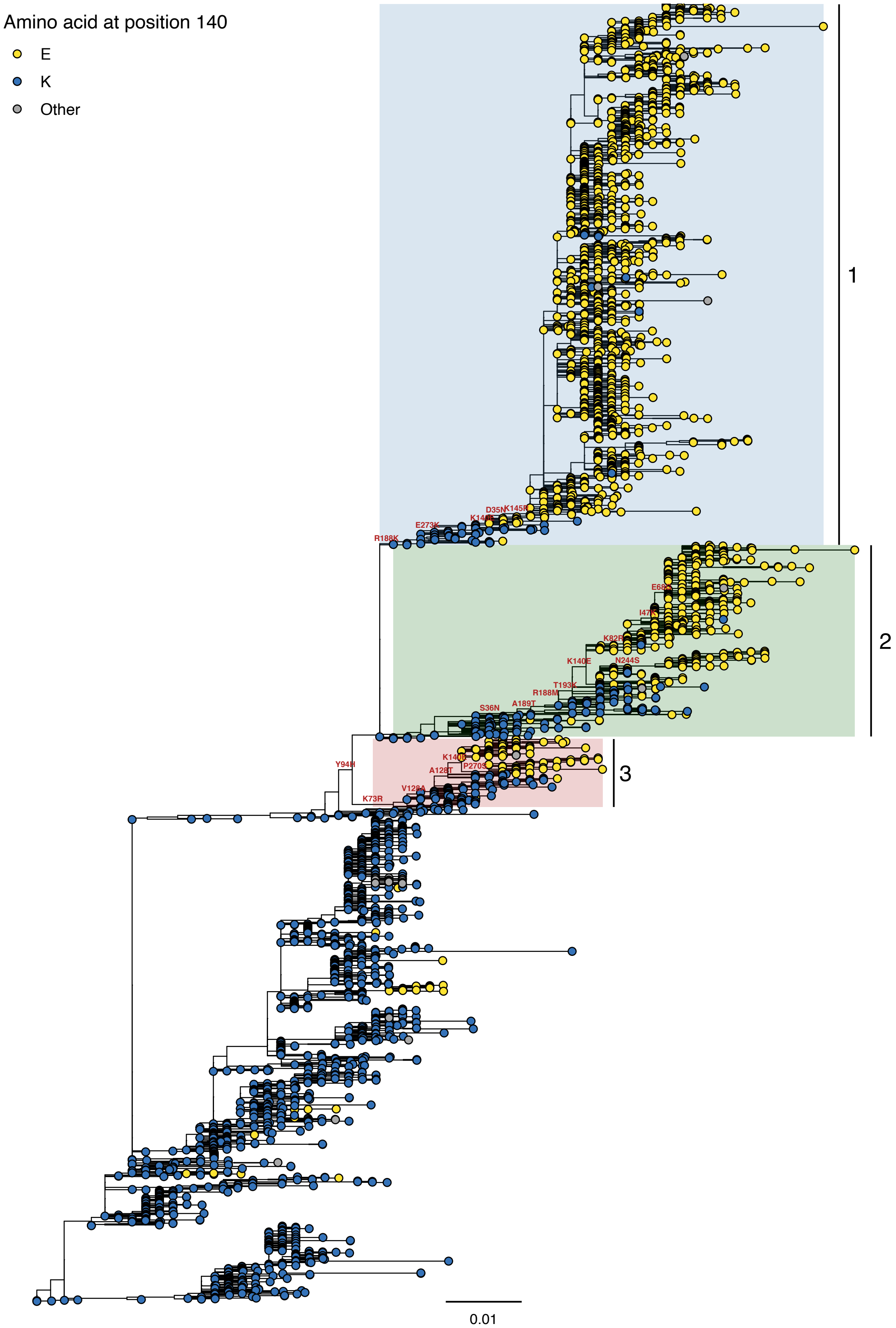


**Fig. A6. Phylogeny of A/H1N1 seasonal viruses for the period 1999 to 2008.** Branch tip colour indicates the amino acid identity at position 140. Co-circulating lineages defined by the K140E fixation are highlighted. Scale bar indicates the number of nucleotide substitutions per site. Tree rooted to A/New Caledonia/20/1999

## 6.4 A/H3N2 2008-2011

In the period 2008-2011, the K158N substitution was only detected once in 2009 before fixing in combination with N189K in the same year in two distinct co-circulating A/H3N2 lineages. N189K was not detected before fixing as a K158N/N189K double substitution. The combination of the substitutions resulted in an antigenic phenotype switch from Wisconsin/2005-like to the Perth/2009-like and Victoria/2009-like phenotypes respectively. Residues 158 and 189 are both located in antigenic site B adjacent to the receptor binding site, with substitutions at both residues characterized as cluster-transition substitutions in multiple historic A/H3N2 cluster transitions (Koel et al., 2013). The HA-defined evolutionary trajectories of viruses from the Perth/2009-like lineage and Victoria/2009-like lineages from their most recent common ancestor are characterized by distinct sets of substitutions (table A3, Fig. A7). Both lineages acquired substitutions at previously characterized antigenic sites (Koel et al., 2013), but only share the T212A substitution.

Table A3: **Substitutions that characterize the co-circulating genetic / antigenic clades defined by K158N and N189K**

| Lineage | Trunk substitutions from MRCA to K158N/N189K fixation | Trunk substitutions post- K158N/N189K fixation |
|---|---|---|
| 1 (Victoria / 2009-like) | | T212A, S45N, T48A, K92R, Q57H, A198S, V223I, N312S, N278K,Q33R, N145S, G5E, E62V, D53N, E280A, I230V, Y94H, I192T, S199A |
| 2 (Perth / 2009-like) | E62K | N144K, R261Q, I260M, P162S, E50K, V213A, N133D, T212A, R142G |

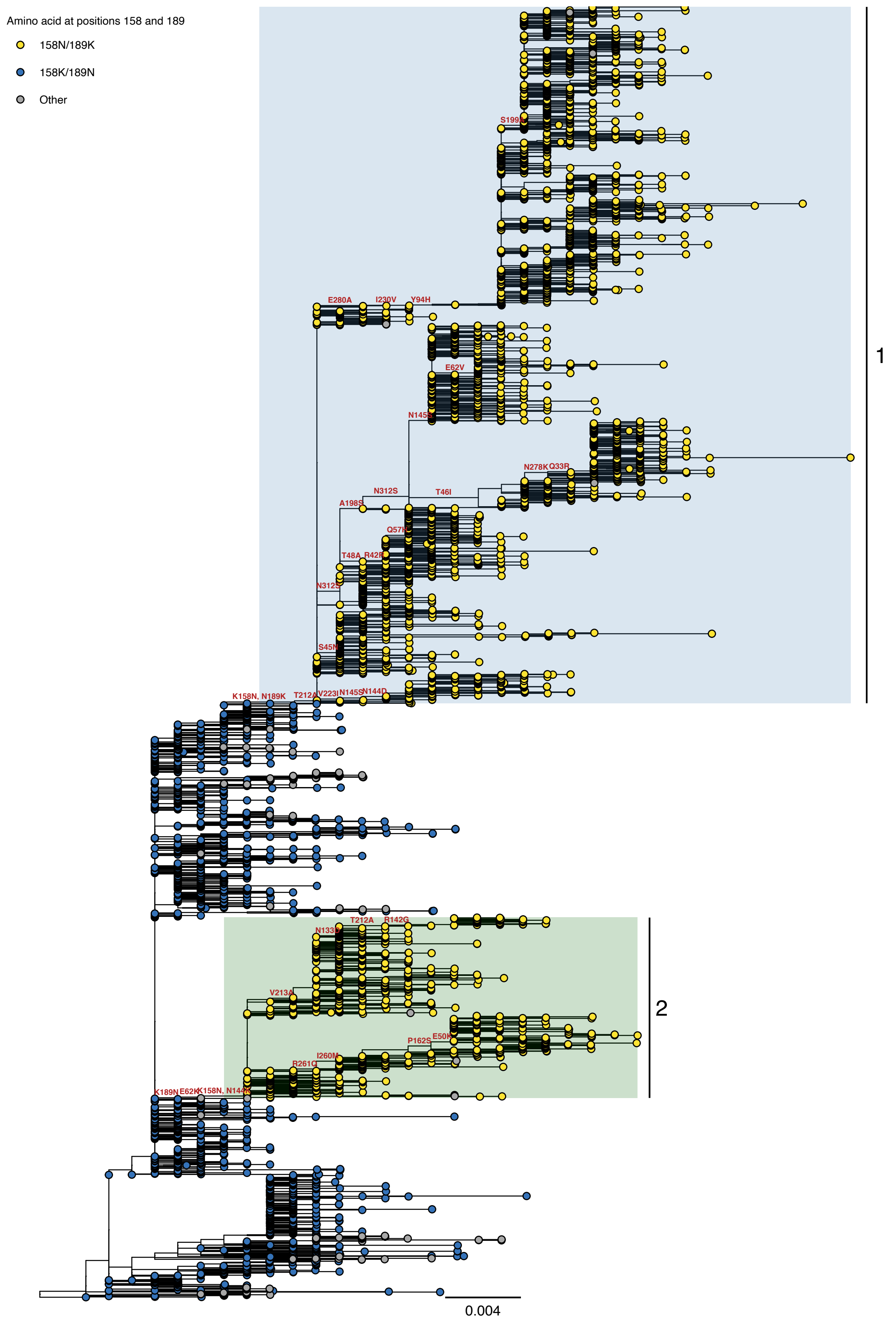


**Fig. A7. Phylogeny of A/H3N2 seasonal viruses for the period 2008 to 2011.** Branch tip colour indicates the amino acid identity at position 158 and 189. Co-circulating lineages defined by the K158N/N189K fixation are highlighted. Scale bar indicates the number of nucleotide substitutions per site. Tree rooted to A/Brisbane/10/2007.

## 6.5 A/H3N2 2012–2014

In 2014 two independent substitution at position 159, F159S and F159Y, fixed in distinct A/H3N2 lineages co-circulating globally (Fig. A8). The S159Y substitution was detected sporadically in 2012/2013 before fixing in 2014 to define the A/H3N2 clade 3C.2a, which circulated as the dominant clade globally for the next three years. The F159S substitution was not detected in 2012–2013 before reaching fixation in 2014 to define clade 3C.3a, which continued to circulate globally at low frequencies.

Residue 159 is located in antigenic site B. The S159Y substitution in combination with Y155H and K189R substitutions previously resulted in the transition of the A/H3N2 Bangkok/79-like antigenic phenotype to Sichuan/87-like phenotype (Koel et al., 2013).

The two lineages have independent mutational trajectories for their HA gene away from their most recent common ancestor, excluding both acquiring the N225D substitution prior to F159X-fixation, with changes acquired changes at the major antigenic epitopes (Table A4, Fig. A8) (Koel et al., 2013).

Table A4: **Substitutions that characterize the co-circulating genetic / antigenic clades defined by F159Y and F159S**

| Lineage | Trunk substitutions from MRCA to F159X fixation | Trunk substitutions post-F159X fixation |
|---|---|---|
| F159Y (lineage 1, Clade 3C.2a) | L3I, N225D, Q311H, N144S, K160T | R142K, R261L |
| F159S (lineage 2, Clade 3C.3a) | R142G, T128A, A138S, N225D | |

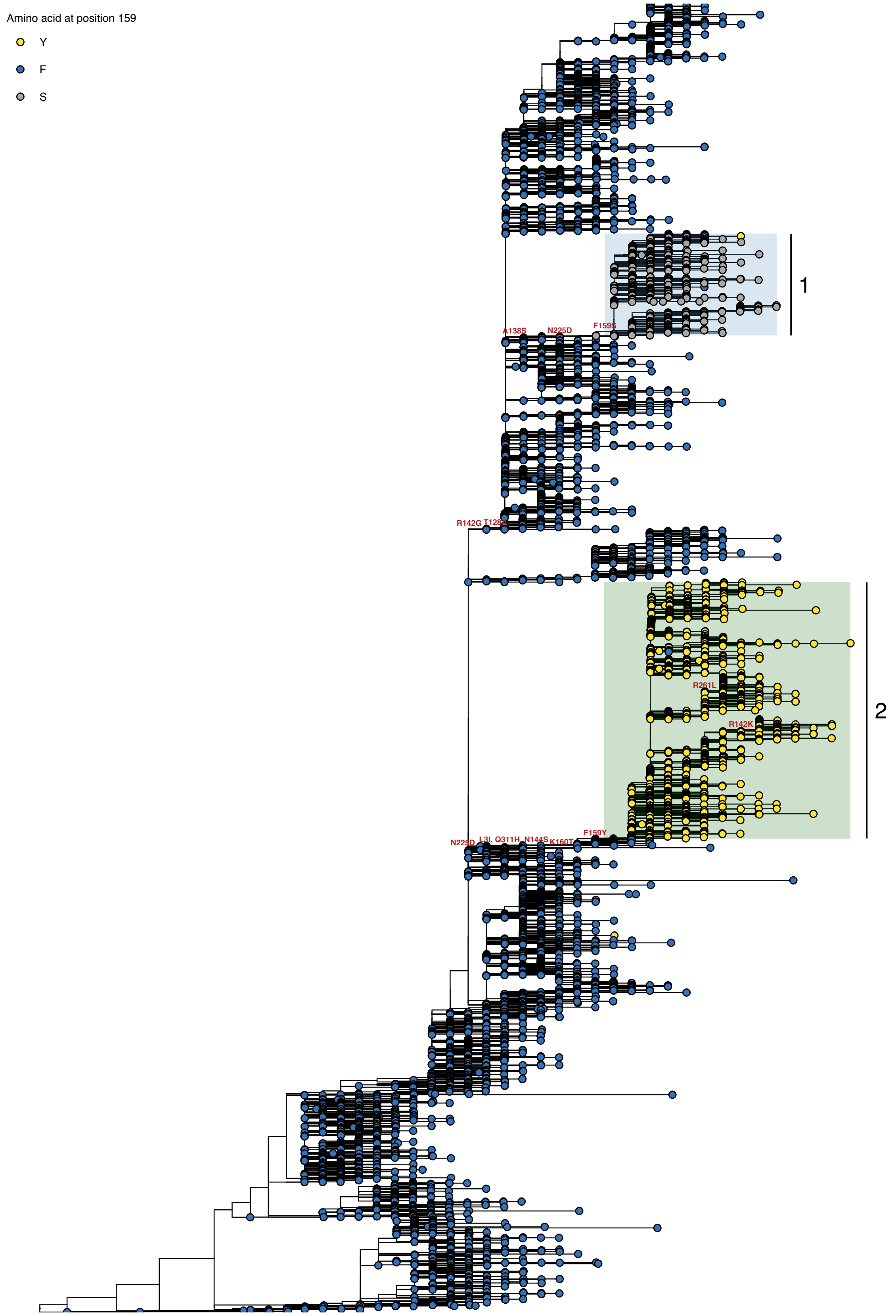


**Fig. A8. Phylogeny of A/H3N2 seasonal viruses for the period 2012 to 2014.** Branch tip colour indicates the amino acid identity at position 159. Co-circulating lineages defined by the F159X fixation are highlighted. Scale bar indicates the number of nucleotide substitutions per site. Tree is rooted to A/Perth/16/2009.

### 7 Prior theoretical studies

In this section, we discuss how our work fits into the substantial existing theoretical literature on influenza virus evolutionary dynamics.

A number of theoretical studies over the last 20 years have addressed the tempo and structure of seasonal influenza virus antigenic evolution, but only two have addressed within-host influenza virus antigenic evolution (Luo et al., 2012; Volkov et al., 2010). Most have focused on population level dynamics (Bedford et al., 2012; Ferguson et al., 2003; Gog, 2008; Koelle et al., 2006; Koelle et al., 2009; Koelle & Rasmussen, 2015; Recker et al., 2007; Strelkowa & Lässig, 2012; Wikramaratna et al., 2013; Zinder et al., 2013).

Our model suggests revisions to the theoretical understanding of within-host and point-of-transmission evolutionary dynamics. The revised paradigm has implications for the population-level models—most notably, it suggests that the rate of population-level antigenic diversification may not be constant over time.

#### 7.1 Prior within-host studies

The two within-host studies both make two key assumptions: (1) immune selection pressure is strong from the moment of inoculation in experienced hosts and (2) influenza virus transmission bottlenecks are wide ($10^5$ virions (Luo et al., 2012); the three most frequent within-host variants deterministically transmitted onward in (Volkov et al., 2010)). As discussed in section 2, antibody-mediated selection pressure is in fact likely to vary substantially over the course of infection. Recent empirical studies have found that transmission bottlenecks are in fact very small (on the order of a single virion (McCrone et al., 2018; Xue & Bloom, 2019)).

Both within-host studies find that new antigenic variants are routinely generated *de novo* during an infection and undergo substantial positive selection during that same infection. NGS studies of natural human infections suggest that this is in fact uncommon (Debbink et al., 2017; McCrone et al., 2018; Xue & Bloom, 2020).

The most crucial assumption in prior work on influenza virus immune escape within-hosts (Luo et al., 2012; Volkov et al., 2010) or within-host pathogen immune escape generally (Kennedy & Read, 2017) is that immunity protects against observable reinfection by neutralizing pathogens during the timecourse of replication. This is immunologically unrealistic for a virus that replicates as rapidly as influenza virus; it “outruns” the memory B-cell response.

What is more, our models show that it protection via neutralization produces binary outcomes: either no reinfection, or detectable reinfection with exclusively mutant viruses. This binary outcome was previously considered a feature, not a bug, because the frequency of homotypic reinfection was not yet known.

A model of immune and therapeutic escape for a generic pathogen (Kennedy & Read, 2017) studied ideas related to replication selection to argue that prophylactic anti-pathogen interventions (such as pre-exposure vaccination) could be less vulnerable to pathogen evolutionary escape than therapeutic interventions (such as post-symptomatic courses of antivirals). Therapeutic interventions occur after a number of pathogen replication events and therefore select upon on a larger, more diverse array of potential pathogen variants. Prophylactic interventions may limit the number of replications before clearance and thereby limit opportunities for diversification. Like the influenza-specific models discussed, this general model does not address the question of how evolution can be rare in symptomatic or otherwise observable infections, where substantial antigenic diversity can be generated.

Similarly, the antimicrobial resistance literature makes a distinction between "acquired" and "transmitted" (or "primary") drug resistance (Bonhoeffer et al., 1997), but this should not be confused with replication and inoculation selection. Transmitted resistance refers to the acquisition of a resistant infection from a host infected with resistant microbes, without regard for whether those microbes were a minority or majority variant in the transmitting host. Inoculation selection, in contrast, refers to natural selection acting on inoculated diversity that *favors* transmitted drug resistance variants or transmitted new antigenic variants. One variant present in a mixed infection has a higher probability of surviving the transmission bottleneck or becoming the majority variant in the new host than its frequency in the transmitting host alone would imply, simply because the recipient host is more susceptible to that variant than to any competing inoculated variants.

#### 7.2 Limits to immune escape through non-variant-specific limiting factors

Two studies (Hartfield & Alizon, 2014; Luo et al., 2012) have previously noted that the evolutionary emergence of escape mutants or otherwise fitter pathogens can be made more difficult due to ongoing competition from the old variant for a shared resource, whether susceptible cells within a host, as in (Luo et al., 2012) and in our model, or susceptible hosts within a population, as in (Hartfield & Alizon, 2014).

In our paradigm, non-variant-specific limiting factors such as target cell depletion play there roles. (1) Denying new variants the opportunity to rise in frequency once an antibody response has been mounted and fitness differences have become substantial. (2) Explaining why immune-compromised hosts, in whom these non-antigenic limiting factors are weaker, can select for antigenic mutants at the scale of a single infection. (3) Clearing infections of naive hosts well before an adaptive immune response can select an escape variant. Point (3) has been posited previously by (Luo et al., 2012), but to our knowledge (1) and (2) are novel.

Hartfield and Alizon (2014) focus on the probability that a fit mutant variant the emerges at the host population scale goes stochastically extinct before causing a large epidemic, and show that competition for hosts with the old variant raises this extinction probability. While this may be relevant to influenza viruses at the population level (see section 8.6 below), an analogous within-host argument is unlikely to be a sufficient explanation for the rarity of influenza virus antigenic mutants within human hosts. New variants are likely to be generated when an influenza virus infection is still well within its exponential growth phase, competition for susceptible cells is relatively unimportant, and probability of stochastic extinction for all incipient lineages is therefore low (due also to influenza's high $\mathcal{R}_0$). The absence of selection, rather than a high rate of stochastic loss of mutants during within-host replication, is the most important component of our explanation for the rarity of observable influenza virus antigenic evolution within individual human hosts.

Note that Hartfield and Alizon also have a model of within-host immune escape for a general pathogen, but that model is designed to estimate the contribution of an active adaptive immune response to increasing or reducing the probability that a small escape mutant population goes stochastically extinct (Hartfield & Alizon, 2015).

For influenza viruses, the *de novo* mutation that creates a stable new variant lineage of interest typically precedes time of adaptive immune proliferation, so a distinct model is required.

#### 7.3 Fitness costs for antigenic mutants

Previous population-level studies have argued that non-antigenic fitness costs associated with antigenic substitutions (Gog, 2008; Kucharski & Gog, 2011) or deleterious mutational load on the influenza virus genome (Koelle & Rasmussen, 2015) are necessary to explain the pace of influenza virus antigenic evolution in the human population. These models introduce population-level antigenic novelty at rates 5–10 times higher than predicted by inoculation selection, and at equal rates early and late during the circulation of a particular antigenic variant.

There are empirical reasons to doubt that influenza viruses are in fact antigenic diversity-limited due to fitness costs. We find multiple instances of polyphyletic cluster transitions: a cluster transition amino acid substitution arises and proliferates on two or more branches of the virus phylogeny at this same time (see section 6). This is inconsistent with the hypothesis that antigenic variants are either intrinsically unfit (deleterious substitutions) or incidentally unfit (poor background) prior to the moment that they proliferate at the population level (Gog, 2008; Koelle & Rasmussen, 2015; Kucharski et al., 2015), since the cluster transition mutant can reach surveillance-detectable levels even before substantial population immunity has accumulated (suggesting that strong antigenic selection is not required to overcome intrinsic deleteriousness), and it can fully emerge when the time is ripe against multiple independent genetic backgrounds, suggesting background may not suffice to constrain diversification earlier in a cluster's circulation.

In the absence of meaningful replication selection, the population level rate of antigenic diversification is the average rate at which new variants survive all transmission bottlenecks to found or co-found infections. In an inoculation selection regime, that rate is unlikely to surpass 2 in $10^4$ inoculations (Fig. 6A). We obtain that estimate assuming no within-host replication cost for antigenic mutants, weak cross immunity between mutant and old variant viruses, and many experienced hosts who reliably neutralize old variant virions at the point of transmission. Actual rates of new variant survival are almost certainly lower. First, we have assumed that a new antigenic variant can be produced by a single amino acid substitution (accessible via a single nucleotide substitution), however a recent detailed study of the antigenic evolution of A/H3N2 viruses shows that some new variants require more than one substitution (Koel et al., 2013). Second, some (possibly many) previously infected hosts will not possess well-matched antibodies due to original antigenic sin (Davenport & Hennessy, 1956), antigenic seniority (Lessler et al., 2012), immune backboosting (Fonville et al., 2014), or other sources individual-specific variation in antibody production (Lee et al., 2019). These factors individually, and particularly in combination, which have not been modelled in this study mean that the above 2 in $10^4$ transmission estimate is likely to be unrealistically high. By comparison, existing population level models that invoke fitness costs introduce population-level antigenic diversity at much higher rates. One study (Koelle & Rasmussen, 2015) introduces new antigenic variants at a rate of 7.5 per $10^4$ transmissions; another (Kucharski & Gog, 2011) introduces new population-level mutations at a continuous rate of $6.8 \times 10^{-4}$ mutations per infected individual per day, which should produce new variant infections by 2 days post infection in at least 1 of every $10^3$ infected hosts:

$$1 - \exp\left(-6.8 \times 10^{-4} * 2\right) \approx 0.0013$$

Prior to the accumulation of population immunity, infections dominated by new variants should be rare, and new variants should be nearly neutral relative to the old variant at the population level. This may be sufficient to explain the absence of diversification early in the circulation of a cluster without invoking non-antigenic fitness costs. That said, additional constraints on the virus, particularly those that reduce the within-host fitness of new antigenic variants, should further slow virus antigenic diversification by reducing variant frequencies in donor hosts and thereby reducing the probability that the variants survive transmission bottlenecks. Furthermore, in the absence of substantial population immunity, adaptive substitutions may be lost at the population level due to competition from lineages with non-antigenic adaptive substitutions. Our work is thus consistent with a role for clonal interference in population level influenza virus antigenic evolution (Strelkowa & Lässig, 2012).

#### 7.4 Further discussion of alternative explanations for rare within-host new antigenic variants

In this section, we expand on the discussion of alternative hypotheses from the main text (see Alternative explanations for rare new antigenic variants).

##### 7.4.1 Protection through neutralization during early viral replication, i.e. an immediate recall response

Adaptive immune pressure could be strong enough that $\mathcal{R}^w(0) < 1$. In that case, an old variant infection dies out if it does not produce a mutant virion, and it only has, on average, $\frac{b}{1-\mathcal{R}^w(0)} - b$ replication events in order to do so before the infection is cleared (average replication events in $b$ independent subcritical branching processes with branching parameter $\mathcal{R}^w(0)$, see section 3.6), so antigenic mutants can be rare in homotypically challenged hosts, depending upon $b$ and and the mutation rate $\mu$ (Luo et al., 2012).

As we note in the main text, this makes two predictions that do not match empirical reality: binary outcomes to homotypic challenge—no infection or infection with an antigenic mutant—and frequent evolution of antigenic mutants in infections of intermediately immune hosts.

Recent empirical work (Debbink et al., 2017; Javaid et al., 2020; McCrone et al., 2018) and human challenge studies (Clements et al., 1986; Memoli et al., 2019) both suggest that detectable reinfection of experienced hosts frequently occurs without observable immune escape. This contradicts the binary outcome.

A model of influenza virus evolution (Volkov et al., 2010) found that even a small number of intermediately immune hosts along a transmission chain should reliably promote the evolution of new antigenic variants. The model predicted this because it features an immediate recall response, which implies efficient replication selection in intermediately immune hosts. In that model, intermediately immune hosts can be productively infected with the old variant ($\mathcal{R}^w(0) > 1$), so they reliably generate antigenic mutants. Those mutants then immediately undergo positive selection and are often transmitted onward. We see the same phenomenon in our own transmission chain model when there is an immediate recall response (see main text Fig. 5).

Intermediate-to-strong immunity to old variant viruses is likely to be common in the human population even at the beginning of a new antigenic cluster's circulation (Fonville et al., 2014), and yet new variants are rarely observed until a new variant has circulated for multiple years (Smith et al., 2004). This suggests that intermediately immune hosts do not generate and select for new antigenic variants as readily as the model in (Volkov et al., 2010) implies.

Our proposed model of immune selection can explain why intermediately immune hosts may not reliably select for new antigenic variants. A realistically timed recall response makes replication selection unlikely. Rates of inoculation selection are limited in all hosts due to low transmitting host mutant frequencies $f_{\mathrm{mt}}$. Intermediately immune hosts who possesses sIgA that are not well-matched to the old variant may additionally have a relatively low value of $\kappa_w$ and therefore be little better than a naive host in promoting new variant survival at the point of transmission.

##### Heterogeneous neutralization rates during early viral replication

One possibility is that adaptive immune pressure could be present from the start of replication, but vary substantially among individuals, even those with the same immune history. This heterogeneity could explain why some individuals ($k$ small, $\mathcal{R}^w(0) > 1$) are productively reinfected with old variant viruses while others ($k$ large, $\mathcal{R}^w(0) < 1$) are protected. In this way one could have protection via neutralization during the timecourse of infection but still have frequent observable reinfection without immune escape.

While individual variation in immunity does exist (Lee et al., 2019), there are two reasons why this hypothesis is implausible. First, it is unrealistic given our current understanding of the adaptive immune response to influenza viruses, as discussed in section 2, since it requires a substantial antibody response before 48 hours post-infection. Second, such heterogeneous protection would be need to be extremely bimodal to completely miss the regime of $k$ values in which replication selection is efficient.

Even if this unlikely hypothesis of bimodal early adaptive responses were true, there would remain a role for sampling effects at the point of transmission. If individuals either possess sterilizing-strength immunity that acts early in the timecourse of viral replication or possess sufficiently weak immunity that replication selection is unlikely, antigenic evolution would be dominated by cases in which mutants are inoculated into strongly immune hosts, as in simpler models with $\mathcal{R}^w(0) < 1$. Inoculated mutants remain crucial in this scenario.

##### 7.4.2 Deleterious antigenic mutants

Antigenic mutants could be replication-competent, but weakly deleterious within-host in the absence of immune selection and/or compensatory substitutions. Two studies (Gog, 2008; Kucharski et al., 2015) have invoked this hypothesis to explain population-level antigenic dynamics. If neutralization is sufficiently strong during virus replication, however, replication selection can still promote mutants during infections of experienced hosts, even in the absence of compensatory mutations. Within-host replication costs act to decrease the value of the fitness difference $\delta$. Recall that:

$$\delta(t) = \left[g_m(t) - g_w(t)\right] - \left[d_m(t) - d_w(t)\right] \tag{A27}$$

Typically $g_m(t) = g_w(t)$ and so $\delta = k(c_w - c_m)$. Here, $g_m < g_w$. For instance, we could have $r_m = qr_w$, $q < 1$ and therefore $g_m = qg_w$. In that case:

$$\delta = k(c_w - c_m) - g(t)(1 - q) \tag{A28}$$

We estimate $g$ to be order 20 early in infection and declining subsequently. $k$ may well be order 10. So it is very plausible that a weakly deleterious mutant ($q = 0.95$, for instance) could still undergo substantial replication selection early in infection if it offered enough immune escape ($c = 0.70$) and antibodies were present ($t_M$ small).

The effect could be particularly extreme in intermediately immune hosts if immunity drops superlinearly with antigenic distance. If the old variant is neutralized at a rate $kc_w = 4$ but the mutant is barely neutralized at all $c_m \approx 0$, it could easily overcome even substantial within-host deleteriousness.

In the absence of replication selection, even weak within-host deleteriousness should lead mutants to be rapidly purged (purifying selection should be efficient for phenotypes subject to replication selection) (Sigal et al., 2018), reducing $f_{\text{mt}}$. Within-host deleteriousness would therefore reduce the rate at which new variants survive bottlenecks, since $p_{\text{surv}}$ and $p_{\text{drift}}$ both decrease as $f_{\text{mt}}$ decreases (see section 4.4).

A final possibility is that antigenic mutants could be extremely deleterious in the absence of compensatory mutations: $q$ small enough that $\delta < 0$ even when $t > t_M$ (i.e. the old variant is more fit within-host than the new variant even in the presence of a well-matched adaptive immune response). This scenario is unlikely given the strength of recall responses, but it would make inoculation selection and founder effects even more important for population-level antigenic diversification.

### 8 Further implications

#### 8.1 The importance of inoculated diversity

(Kennedy & Read, 2017) and (Luo et al., 2012) study immune escape with $\mathcal{R}^w(0) < 1$ but consider only diversity generated after inoculation. Our study complicates their results: we find that when $\mathcal{R}^w(0) < 1$ the dominant mode of antigenic evolution will be selection on inoculated diversity, not selection on generated diversity (Figure 2). The pace of evolution will depend on how likely an escape mutant is to be transmitted to an experienced host. If bottlenecks in experienced hosts are wide, evolution may be rapid. If these bottlenecks are tight or escape mutants are deleterious within untreated naive hosts (so $f_{\mathrm{mt}}$ is low), then $\mathcal{R}^w(0) < 1$ at the start of infection can result in slow evolution, since both replication and inoculation selection will be rare.

In the absence of mucosal neutralization with small $f_{\mathrm{mt}}$, $b \ll \frac{1}{f_{\mathrm{mt}}}$, and small $\mu$, the threshold for where inoculation selection becomes more common than replication selection with $\mathcal{R}^w(0) < 1$ is approximately

$$f_{\mathrm{mt}} b p_{\mathrm{sse}} > \mu \frac{b\mathcal{R}^w(0)}{1 - \mathcal{R}^w(0)} p_{\mathrm{sse}}$$

or simply:

$$f_{\mathrm{mt}} > \mu \frac{\mathcal{R}^w(0)}{1 - \mathcal{R}^w(0)} \tag{A29}$$

the left-hand side comes from equation A17, which gives an probability that a mutant with $\mathcal{R}^m(0) > 1$ is inoculated. The right-hand side comes from equation 28, which gives the probability that an escape mutant is generated in a declining but replicating virus population before that population goes extinct. $p_{\mathrm{sse}}$, which occurs on both sizes and cancels, is the probability that a mutant survives stochastic extinction. We have also ignored the fact the the right-hand side should have a term representing the probability that inoculation selection does not occur, but as we are considering cases when both inoculation selection and replication selection are rare, that term is approximately one.

On average, $f_{\mathrm{mt}} > \mu$ due the asymmetry between forward and back mutation rates when the mutant is rare. It follows that $f_{\mathrm{mt}} > \mu \frac{\mathcal{R}^w(0)}{1-\mathcal{R}^w(0)}$ should hold when (A) the mutant not is too deleterious in the absence of positive selection (since this reduces $f_{\mathrm{mt}}$) or (B) $\mathcal{R}^w(0)$ is not close to 1 (since then many copies are made on average before the virus goes extinct). Alternatively, if $b$ is sufficiently large—on the order of $\frac{1}{f_{\mathrm{mt}}}$—selection on inoculated diversity will be the dominant mode of evolution simply because most inocula will include mutants.

Selection on inoculated diversity may be important in many host-pathogen systems, not just in influenza virus antigenic evolution. Some anti-microbial resistance in bacteria, for instance, is acquired through the uptake of preexisting plasmids, rather than through *de novo* mutation (Perron et al., 2015). Whether drug resistance emerges in an individual treated host will depend on whether any of the bacteria inoculated bear the needed plasmid.

#### 8.2 Consequences for modeling influenza virus dynamics

Population size, population level antigenic diversification ("mutation") rate, and degree of population structure all substantially impact the proliferation of new influenza virus antigenic variants at the population level. It has been hard for population level models to distinguish plausible hypotheses regarding evolutionary constraints on the virus, because achieving simultaneous realism across in all these effects while retaining model tractability has proven extremely difficult.

For epidemiological models of influenza virus evolution overall population size matters: epidemics in larger populations involve more total inoculations, and therefore more opportunities for new variant survival at the point of transmission to occur. Large host populations with high transmission rates, strong population connectivities, and recurrent epidemics should promote antigenic evolution. This helps explain why influenza virus antigenic evolution appears to occur disproportionately in east and southeast Asia (Russell et al., 2008).

It also reveals an important consideration for interpreting results from individual-based simulation models of global influenza virus evolution. Due to computational constraints, many such models must use host population sizes that are orders of magnitude smaller than the true global human population (e.g. $N = 40$ million (Koelle & Rasmussen, 2015), $N = 90$ million (Bedford et al., 2012)). Such models will have smaller numbers of inoculations than real-world influenza virus dynamics. They therefore run the risk of overestimating rates of strain extinction or, to avoid this, of overestimating per-infection virus diversification rates.

#### 8.3 Egg and mouse passage experiments

The observation that immune escape can be reliably be selected for in egg (Davis et al., 2018) and mouse (Hensley et al., 2009) passage experiments and yet appears rare in reinfected experienced humans is unsurprising in light of our view that influenza virus evolution is limited more by the failure of antibody-mediated selection pressure and antigenic diversity to coincide than by the absence of either.

The egg experiments (Davis et al., 2018) amount to strong replication selection: the viruses were passaged in the presence of sera with highly-specific neutralizing antibodies, with these sera present at all times, including during the exponential growth phase within each egg.

The mouse passage experiments (Hensley et al., 2009) showed that serial passage in immune mice led to the appearance and fixation of antigenic escape mutants, but serial passage in naive mice did not. Several details of the experiment suggest that both replication selection and inoculation selection should have been more possible than in typical infected experienced humans.

First, the mice were intranasally inoculated with 50 $\mu l$ of homogenized lung isolate from the previous mouse in the passage chain at two days post-infection. This is likely a substantially more concentrated dose of virions than is inhaled through aerosol or even contact transmission (an inter-host bottleneck much wider than occurs in humans, to use the terminology of main text Fig. 3A). This could produce a very large value of $v$, the number of virions encountering IgA, and potentially a large value of $b$, the final (cell infection) bottleneck as well—in the presence of sufficiently many virions, early cell infections might be sufficiently simultaneous as to produce have observably diverse within-host populations. All of this should tend to facilitate new variant survival and promotion to observable levels, but also to improve chances, relative to humans, that at least some old variant virions could survive alongside the new variant.

Second, inoculations were also carried out until a successful inoculation could be achieved. This further improves the chances of observing an escape mutant selected at the point of transmission. Finally, vaccinated mice in the passage chain were inoculated 10–21 days post-vaccination, meaning antibody levels could be higher than the memory baseline. This would be expected to strengthen inoculation selection and perhaps permit replication selection.

One detail in supplementary table 1 of the mouse passage study (Hensley et al., 2009) is particularly relevant. In the passaging of virus stock #3, two mutants were observed for the first time in the second vaccinated mouse in the chain, but neither at fixation. Both remained present for 5 further passages in experienced mice without either being lost or fixing. This suggests a much wider final bottleneck than occurs in naturally-infected humans (McCrone et al., 2018). It also suggests that replication selection is unlikely to have been at all strong if present at all: over repeated exponential growth phases of two days with wide bottlenecks, even very small fitness differences can be easily amplified, so the more fit escape mutants should have fixed if they were subject to replication selection. It also suggests, as expected, that inoculation selection is weak if final bottlenecks are wide and neither antigenic variant is reliably neutralized. In that case, a long intermittent period of coexistence is possible.

An example calculation illustrates this: if $v = 1000$, $b = 10$, $f_{\text{mt}} = 0.5$, $\kappa_m/\kappa_w > 0.9$, and $\kappa_w < 0.8$ the mutant has an inoculation-selective advantage, but the expected mutant frequency at after the next transmission event approximately 0.58 or less. Letting $\bar{w} = E(x_w)$ and $\bar{m} = E(x_m)$:

$$\begin{aligned} \bar{w} > v(1 - f_{\text{mt}})(1 - \kappa_w) &= (1000)(0.5)(0.20) = 100 \\ \bar{m} > v(f_{\text{mt}})(1 - \kappa_m) &= (1000)(0.5)(0.28) = 140 \end{aligned} \tag{A30}$$

And since $\bar{w}, \bar{m} \gg b$, the hypergeometric sampling is approximately binomial, and therefore the expected frequency of mutant is just $\frac{\bar{m}}{\bar{m}+\bar{w}} \approx 0.58$

While it is difficult to compare fitness differences during replication to fitness differences in mucosal neutralization, a very small fitness difference of $\delta = 0.5$ should raise the fitter type from frequency 0.5 to frequency 0.73 over the course of two days.

#### 8.4 Magnitude of antigenic change and population level patterns

When antigenic effect size of mutations approaches zero, the effects of stochastic loss and mistimed selection pressure dominate, and inoculation and replication selection become sufficiently weak that true drift dominates as a force for introducing new antigenic variants. In such a regime, we expect the influenza viruses to evolve gradually, fail to cause large epidemics, and possibly diversify, not unlike influenza B viruses (Bedford et al., 2015). In particular, slow spread enables a kind of immunological niche partitioning: if population immune histories are not spatially uniform because epidemiological spread is slow relative to antigenic diversification, multiple variants that are each locally favored in distinct areas could co-circulate and form separate lineages, as has happened with B/Victoria and B/Yamagata.

When antigenic jump size approaches the maximum size possible, such that all population immunity disappears each time there is an antigenic cluster transition (Smith et al., 2004), we expect influenza viruses to follow a strongly clock-like pattern with high-amplitude oscillations in case numbers. They would cause massive punctuated epidemics during jump years and then in the next year either select for a new variant or go extinct. There would be huge booms followed by one or more years of bust. In between, at intermediate degrees of escape, a punctuated but somewhat less clocklike pattern—such as the pattern observed in nature for A/H3N2 viruses (Smith et al., 2004)—becomes possible.

#### 8.5 Preview substitutions and mutation limitation

"Preview" substitutions and polyphyletic cluster transitions also imply that the influenza virus is not typically mutation-limited at the population level—a substantial-effect escape mutant is usually accessible in sequence-space—but that the virus may nonetheless be highly constrained in its evolutionary trajectory. The virus repeatedly finds the same substitution as a solution to its antigenic evolutionary problem. This could occur either because only one escape mutant is available or because one of the available escape mutants provides substantially more immune escape (on average) than the others.

#### 8.6 Selection on bottlenecked diversity at higher scales

Without an explanation for why within-host dynamics do not more frequently promote antigenic mutants, it is difficult to explain the slow pace and noisy trajectory of influenza virus evolution. But such a within-host explanation, though likely necessary, may not be sufficient. It is conceivable that in a sufficiently large and well-connected global host population, population-level antigenic selection favoring mutants with higher $\mathfrak{R}_e$ would lead even small effect antigenic mutants rapidly to emerge and fix.

One important mechanism by which evolution might be also slowed at higher scales is analogous to the within-host dynamic of founder effects and inoculation selection: mutant lineages may be frequently lost (though perhaps selectively favoured) at the bottleneck that occurs between distinct influenza virus epidemics.

Prior modeling work has found that population-level host competition between pathogen variants reduces the probability that a fit mutant variant causes a large epidemic in the population in which it first emerges (Hartfield & Alizon, 2014). This is likely to be relevant in influenza virus epidemics, since the virus is thought to provoke short-term strain-transcending (i.e. variant-transcending) immunity (Ferguson et al., 2003).

We propose a previously unexplored consequence of this argument: successful establishment of a mutant lineage at the population level requires that the mutant lineage be exported to a new host subpopulation where susceptible hosts are common. This is rare but potentially selective sampling event.

When a new variant lineage emerges at the population level, it is likely to be surrounded by many propagating old variant lineages due to the generator-selector dynamic discussed in the main text (see main text Fig. 6B,C). Moreover, most mutant lineage generation events will occur as an epidemic is peaking, since that is when the most inoculations occur. The consequence is that most generated population-level mutant lineages will encounter severe competition for susceptible hosts (especially if we consider realistic, spatial host contact networks rather than well-mixed epidemic models). So a generated mutant lineage is unlikely to account for many cases—or even necessarily more than one case—in the epidemic in which it emerges.

Many local influenza virus epidemics are likely to be evolutionary dead ends with no cases exported that establish chains of transmission in other locations. Because only a minority of the human population is likely to travel to or from the site of any particular epidemic, the majority of virus diversity generated within each epidemic is likely to be lost in between-epidemic bottlenecks. Conditional on being exported, new variant lineages could have competitive advantages over old variant lineages because they spread should spread more rapidly and go stochastically extinct less easily due to their higher $\mathfrak{R}_e$. It follows that there is analogous dynamic to inoculation selection at the population level: mutant lineages are rarely exported from one sub-population (host, epidemic) to another, but conditional on being exported, they have an advantage in any potential competition with exported old variant lineages.

The rate of proliferation of mutants at the population level, then, may be limited by the rarity of early generation of mutant lineages during an epidemic (analogous to the rarity of very early within-host *de novo* generation of mutants) and by the rarity of successful mutant lineage exportation when generation is not early (analogous to the rarity of mutant virions surviving the transmission bottleneck between hosts).

We aim to explore this argument with a formal mathematical model in future work.

### 9 Mathematical derivations in full

#### 9.1 Full derivation of the within-host replicator equation

The derivation in this section establishes equation 12:

$$\frac{\mathrm{d}f_m}{\mathrm{d}t} = f_m(1-f_m)\Big(\big[g_m(t)-g_w(t)\big] - \big[d_m(t)-d_w(t)\big]\Big)$$

*Derivation.* We note that:

$$f_m(t) := \frac{V_m(t)}{V_{\text{tot}}(t)}$$

where $V_{\text{tot}}(t) = V_w(t) + V_m(t)$. Let $\dot{V_i}$ denote $\frac{\mathrm{d}V_i}{\mathrm{d}t}$. Note that $\dot{V_i} = V_i\alpha_i(t)$ where $\alpha_i(t) = g_i(t) - d_i(t)$, and that $\frac{V_w(t)}{V_{\text{tot}}(t)} = 1 - f_m(t)$. By the quotient rule:

$$\frac{\mathrm{d}f_m}{\mathrm{d}t} = \frac{V_{\text{tot}}\dot{V}_m - V_m(\dot{V}_m + \dot{V}_w)}{V_{\text{tot}}^2}$$

$$= \frac{V_{\text{tot}}(\alpha_m V_m) - V_m(\alpha_m V_m + \alpha_w V_w)}{V_{\text{tot}}^2}$$

Dividing through by $V_{\text{tot}}^2$ yields:

$$\alpha_m f_m - f_m(\alpha_m f + \alpha_w(1 - f_m))$$

$$= \alpha_m f_m - \alpha_m f_m^2 - \alpha_w f_m(1 - f_m)$$

$$= \alpha_m f_m(1 - f_m) - \alpha_w f_m(1 - f_m)$$

$$= f_m(1 - f_m)(\alpha_w - \alpha_m)$$

$$= f_m(1 - f_m)\Big(\big[g_m(t) - g_w(t)\big] - \big[d_m(t) - d_w(t)\big]\Big)$$

■

#### 9.2 Replicator equation with symmetric mutation

With symmetric mutation at a rate $\mu$, the replicator equation remains calculable.

*Derivation.* Given symmetric mutation, $\dot{V}_i = V_i\alpha_i(t) + \mu g_j(t)V_j$ where $\alpha_i(t) = (1 - \mu)g_i(t) - d_i(t)$

Substituting in to the previous derivation yields:

$$\frac{\mathrm{d}f_m}{\mathrm{d}t} = \frac{V_{\text{tot}}(\alpha_m V_m + \mu g_w(t)V_w) - V_m(\alpha_m V_m + \mu g_w(t)V_w + \alpha_w V_w + \mu g_m(t)V_m)}{V_{\text{tot}}^2}$$
$$= \alpha_m f_m + \mu g_w(t)(1 - f_m) - f_m\Big(\alpha_m f_m + \mu g_w(t)(1 - f_m) + \alpha_w(1 - f_m) + \mu g_m(t)f_m\Big)$$

$$= \alpha_m f_m + \mu g_w(t)(1 - f_m) - \alpha_m f_m^2 - \mu g_w(t)f_m(1 - f_m) - \alpha_w f_m(1 - f_m) - \mu g_m(t)f_m^2$$
$$= \alpha_m f_m(1 - f_m) - \alpha_w f_m(1 - f_m) + \mu\left(g_w(t)(1 - f_m) - g_w(t)f_m(1 - f_m) - g_m(t)f_m^2\right)$$
$$= f_m(1 - f_m)(\alpha_m - \alpha_w) + \mu\Big(g_w(t) - 2g_w(t)f_m + g_w(t)f_m^2 - g_m(t)f_m^2\Big)$$

Note that if $g_w(t) = g_m(t) = g(t)$, this simplifies to:

$$\frac{\mathrm{d}f_m}{\mathrm{d}t} = f_m(1 - f_m)\big[d_m(t) - d_w(t)\big] + \mu g(t)(1 - 2f_m)$$

■

#### 9.3 Replicator equation with one-way mutation

*Derivation.* To find the case with one-way mutation, we simply let all $\mu g_m(t)V_m$ terms from the symmetric mutation replicator equation be zero. This yields:

$$\frac{\mathrm{d}f_m}{\mathrm{d}t} = f_m(1 - f_m)(\alpha_m - \alpha_w) + \mu g_w(t)\left(1 - 2f_m + f_m^2\right) \tag{A31}$$

■

#### 9.4 Derivation of $t^*(x, t)$

This establishes the expression for the time $t^*$ by which a mutant must emerge to reach at least frequency $x$ by time $t$ given in equation 25 of the Methods.

$$t^*(x,t) = \begin{cases} t^*_-(x,t) & t^*_-(x,t) < t \text{ and } t^*_-(x,t) \leq t_M \\ t^*_+(x,t) & t^*_+(x,t) < t \text{ and } t^*_-(x,t) > t_M \\ t & \text{otherwise} \end{cases}$$

where:

$$t^*_-(x,t) = \frac{\ln\left[\frac{1-x}{x}\exp\big(\delta(t - t_M)\big) + 1\right] - \ln b}{g_0 - d_v}$$

$$t^*_+(x,t) \approx \frac{\ln\left(\frac{1-x}{x}\right) - \ln(b) + \delta t - c_w k t_M}{g_0 - d_v - c_w k + \delta}$$

*Derivation.* The frequency of the mutant at time $t$ depends on the quantity

$$\psi(t) = \exp\big(\delta(t - \max\{t_M, t_e\})\big)$$

Neglecting ongoing forward mutation the mutant must emerge at some frequency $f_e$ in order to reach frequency $x$ by time $t$:

$$\frac{\psi(t)}{\psi(t) + f_e^{-1} - 1} \geq x$$

Solving for $f_e^{-1}$ yields:

$$f_e^{-1} \leq \left(\psi(t)\frac{1-x}{x}\right) + 1 \tag{A32}$$

$f_e^{-1}$ is determined by the number of old variant virions at time $t_e$. Early in infection, the old variant population grows near-exponentially at a rate $G_0 = g_0 - d_v$. There are two cases: either time of new variant emergence (first mutation to produce a new variant lineage that evades stochastic extinction) occurs before the antibody response is present ($t_e \leq t_M$) or it emerges once the antibody response is already present ($t_e > t_M$).

##### 9.4.1 Case 1: $t_e \leq t_M$

It follows that if $t_e \leq t_M$, $f_e^{-1} \approx b\exp\big((g_0 - d_v)t_e\big) = b\exp(G_0 t_e)$.

$$be^{G_0 t_e} \leq \left(\psi(t)\frac{1-x}{x}\right) + 1$$

Taking the natural log of both sides and subtracting off $\ln b$ from both sides:

$$G_0 t_e \leq \ln\left(\left(\psi(t)\frac{1-x}{x}\right) + 1\right) - \ln b$$

$$t_e \leq \frac{\ln\left(\psi(t)\frac{1-x}{x} + 1\right) - \ln b}{G_0}$$

This establishes a value for $t^*$ when $t_e < t_M$. In that case, $\psi(t) = \exp\big(\delta(t - t_M)\big)$, and:

$$t^*_-(x,t) = \frac{\ln\left[\frac{1-x}{x}\exp\big(\delta(t - t_M)\big) + 1\right] - \ln b}{g_0 - d_v} \tag{A33}$$

##### 9.4.2 Case 2: $t > t_e > t_M$

If $t > t_e > t_M$, we must change our estimate of $f_e^{-1}$, because the old variant population grows more slowly in the presence of an antibody response. The approximate growth rate is $G_1 = G_0 - c_w k$. For $t_e > t_M$:

$$f_e^{-1} = b\exp\left[G_0 t_M\right]\exp\left[G_1(t_e - t_M)\right] = b\exp\left[c_w k t_M + G_1 t_e\right] \tag{A34}$$

The expression from A32 no longer yields a closed form solution; it is now a transcendental equation, because $\psi(t)$ now contains $t_e$ terms: $\psi(t) = \exp\big(\delta(t - t_e)\big)$.

To deal with this, we make the approximation:

$$f_m(t) = \frac{\psi(t)}{\psi(t) + f_e^{-1}}$$

This approximation is excellent unless the initial frequency $f_e$ is large, and it always yields an underestimate of $f_m(t)$, . This is desirable; we would like to be pessimistic about the probability of replication selection given early $t_M$ and early $t_e > t_M$ to avoid biasing ourselves in favor of our hypothesis (namely, that replication selection is unrealistically common if $t_M$ is early).

Letting $f_m(t) = x$, the desired frequency at $t$, we wish to solve:

$$x \leq \frac{\psi(t)}{\psi(t) + f_e^{-1}}$$

We find:

$$f_e^{-1}x + \psi(t)x \leq \psi(t)$$

$$f_e^{-1} \leq \psi(t)\frac{1-x}{x}$$

$$b\exp\left(c_w k t_M + G_1 t_e\right) \leq \exp\big(\delta(t - t_e)\big)\,\frac{1-x}{x}$$

$$\ln(b) + c_w k t_M + G_1 t_e \leq \delta t - \delta t_e + \ln\left(\frac{1-x}{x}\right)$$

$$t_e(G_1 + \delta) \leq \ln\left(\frac{1-x}{x}\right) - \ln(b) + \delta t - c_w k t_M$$

$$t_e \leq \frac{\ln\left(\frac{1-x}{x}\right) - \ln(b) + \delta t - c_w k t_M}{G_1 + \delta}$$

So when $t_e > t_M$:

$$t_+^*(x,t) \approx \frac{\ln\left(\frac{1-x}{x}\right) - \ln(b) + \delta t - c_w k t_M}{g_0 - d_v - c_w k + \delta} \tag{A35}$$

Note that since we used an underestimate of $f_m(t)$, this approximate $t^*$ is a lower bound for the true $t^*$; it may be that the new variant can actually emerge later and still successfully be replication-selected to the desired frequency $x$.

Finally, it may be that $t_+^*(x,t) > t$ and $t_-^*(x,t) > t$. This indicates that the mutant will be at at least frequency $x$ if it emerges at $t$ itself. In that case, we therefore have $t^*(x,t) = t$.

Combining:

$$t^*(x,t) = \begin{cases} t_-^*(x,t) & t_-^*(x,t) < t \text{ and } t_-^*(x,t) \leq t_M \\ t_+^*(x,t) & t_+^*(x,t) < t \text{ and } t_-^*(x,t) > t_M \\ t & \text{otherwise} \end{cases} \tag{A36}$$

■

#### 9.5 Derivation of the closed form for $p_{\mathbf{cib}}$

The derivation in this section establishes equation 35:

$$p_{\text{cib}}(\kappa_w, f_{\text{mt}}, wb) = E(p_{\text{cib}}(x_w, 1, b)) = (1 - e^{-\bar{w}})\frac{b}{\bar{w}} + e^{-\bar{w}}\sum_{j=0}^{b-1}\frac{\bar{w}^j}{j!}\left(1 - \frac{b}{j+1}\right)$$

where $\bar{w} = v(1 - f_{\text{mt}})(1 - \kappa_w)$ is the mean number of old variant virions present after IgA neutralization.

*Derivation.*

$$p_{\text{cib}}(x_w, 1, b) = \frac{b}{x_w + 1}$$

$$\begin{aligned} E(p_{\text{cib}}(x_w, 1, b)) &= b \sum_{k=0}^{\infty} \frac{\bar{w}^k}{k!} e^{-\bar{w}} \frac{1}{k+1} \\ &= \frac{b}{\bar{w}} \sum_{k=0}^{\infty} \frac{\bar{w}^{k+1}}{(k+1)!} e^{-\bar{w}} \\ &= \frac{b}{\bar{w}} \sum_{j=1}^{\infty} \frac{\bar{w}^j}{j!} e^{-\bar{w}} \end{aligned}$$

We know that:

$$e^{-\lambda} + \sum_{j=1}^{\infty} \frac{\lambda^j}{j!} e^{-\lambda} = 1$$

since the Poisson is a proper probability distribution and therefore:

$$\sum_{j=1}^{\infty} \frac{\lambda^j}{j!} e^{-\lambda} = 1 - e^{-\lambda}$$

So substituting in:

$$\frac{b}{\bar{w}} \sum_{j=1}^{\infty} \frac{\bar{w}^j}{(j)!} e^{-\bar{w}} = \frac{b}{\bar{w}} (1 - e^{-\bar{w}})$$

This is exact when $b = 1$, but in other cases $p_{\text{cib}}(x_w, 1, b)$ is more properly given by $\max\{\frac{b}{x_w+1}, 1\}$, since whenever we have $x_w + 1 < b$, a new variant survives with certainty. So for each such $x_w < b - 1$ we need to add on correction term equal to $1 - \frac{b}{x_w+1}$, weighted by the probability of that $x_w$ takes on that value: $e^{-\bar{w}} \frac{\bar{w}^{x_w}}{x_w!}$.

Summing from $x_w = 0$ to $x_w = b - 1$ and factoring out $e^{-\bar{w}}$ gives us the complete correction term:

$$e^{-\bar{w}} \sum_{j=0}^{b-1} \frac{\bar{w}^j}{j!} \left(1 - \frac{b}{j+1}\right)$$

Adding that to the expression derived above completes the derivation.

■

#### 9.6 $p_{\text{cib}} = 1$ only if there is no competition

$p_{\text{cib}} < 1$ if $\kappa_w < 1$ and $f_{\text{mt}} < 1$, and $p_{\text{cib}} = 1$ if $\kappa_w = 1$ or $f_{\text{mt}} = 1$.

*Derivation.* This is most easily seen by rewriting $p_{\text{cib}}$ as an infinite sum:

$$p_{\text{cib}}(\kappa_w, v, b) = e^{-\bar{w}}\Big[\sum_{j=0}^{b-1}\frac{\bar{w}^j}{j!}1 + \sum_{j=b}^{\infty}\frac{\bar{w}^j}{j!}\frac{b}{j+1}\Big] \tag{A37}$$

If $\bar{w} = v(1 - f_{\text{mt}})(1 - \kappa_w) = 0$, we have $p_{\text{cib}} = 1$, as expected. Since $v > 0$, this occurs either when there are no old variant virions ($f_{\text{mt}} = 1$) or when the old variant is neutralized with probability 1 ($\kappa_w = 1$). Otherwise, $\bar{w} > 0$ and we therefore have:

$$p_{\text{cib}} = e^{-\bar{w}}\Big[\sum_{j=0}^{b-1}\frac{\bar{w}^j}{j!}1 + \sum_{j=b}^{\infty}\frac{\bar{w}^j}{j!}\frac{b}{j+1}\Big] < e^{-\bar{w}}\Big[\sum_{j=0}^{b-1}\frac{\bar{w}^j}{j!}1 + \sum_{j=b}^{\infty}\frac{\bar{w}^j}{j!}1\Big] = 1 \tag{A38}$$

The last step uses the fact that the Poisson distribution is a proper probability distribution, and therefore sums to 1. ■

#### 9.7 $p_{\text{cib}}$ is decreasing in $\bar{w}$

$p_{\text{cib}}$ is decreasing in $\bar{w}$: the more competing old variant virions, the less likely the new variant is to pass through the cell infection bottleneck.

*Derivation.* For $b = 1$,

$$\frac{\mathrm{d}p_{\text{cib}}}{\mathrm{d}\bar{w}} = \frac{e^{-\bar{w}}\left(\bar{w} - e^{\bar{w}} + 1\right)}{\bar{w}^2}$$

This is negative for $\bar{w} > 0$, as $e^x > x + 1$ for $x > 0$.

That $p_{\text{cib}}$ is decreasing in $\bar{w}$ for $b > 1$ can be seen by writing:

$$\begin{aligned} p_{\text{cib}} &= e^{-\bar{w}}S(\bar{w}) \\ S(\bar{w}) &:= \sum_{j=0}^{b-1}\frac{\bar{w}^j}{j!} + \sum_{j=b}^{\infty}\frac{\bar{w}^j}{j!}\frac{b}{j+1} \end{aligned} \tag{A39}$$

This can be rewritten in two useful ways:

$$S(\bar{w}) = 1 + \sum_{j=1}^{b-1}\frac{\bar{w}^j}{j!} + \sum_{j=b}^{\infty}\frac{\bar{w}^j}{j!}\frac{b}{j+1} \tag{A40}$$

$$S(\bar{w}) = \sum_{j=0}^{b-2}\frac{\bar{w}^j}{j!} + \sum_{j=b-1}^{\infty}\frac{\bar{w}^j}{j!}\frac{b}{j+1} \tag{A41}$$

This uses the fact that $\frac{b}{j+1} = 1$ when $j = b - 1$.

We begin by showing that $S'(\bar{w}) < S(\bar{w})$. We differentiate A40 with respect to $\bar{w}$:

$$\begin{aligned} S'(\bar{w}) &= 0 + \sum_{j=1}^{b-1} \frac{\bar{w}^{j-1}}{(j-1)!} + \sum_{j=b}^{\infty} \frac{\bar{w}^{j-1}}{(j-1)!} \frac{b}{j+1} \\ &= \sum_{j=0}^{b-2} \frac{\bar{w}^{j}}{j!} + \sum_{j=b-1}^{\infty} \frac{\bar{w}^{j}}{j!} \frac{b}{j+2} \end{aligned} \tag{A42}$$

The first $b-1$ terms in $S'(\bar{w})$ are equal to the corresponding terms in equation A41 for $S(\bar{w})$. Each subsequent term is smaller in $S'(\bar{w})$ than in $S(\bar{w})$, since $\frac{b}{j+2} < \frac{b}{j+1}$ for $b, j > 0$. So we have established that $S'(\bar{w}) < S(\bar{w})$ for $b > 1$.

Now we find $\frac{\mathrm{d}p_{\mathrm{cib}}}{\mathrm{d}\bar{w}}$:

$$\frac{\mathrm{d}p_{\mathrm{cib}}}{\mathrm{d}\bar{w}} = -e^{-\bar{w}} S(\bar{w}) + e^{-\bar{w}} S'(\bar{w}) \quad = e^{-\bar{w}} \Big[ S'(\bar{w}) - S(\bar{w}) \Big] < 0 \tag{A43}$$

since $e^{-\bar{w}} > 0$ and $S'(\bar{w}) - S(\bar{w}) < 0$.

It follows that $p_{\mathrm{cib}}$ is decreasing in $\bar{w}$. ■

#### 9.8 $p_{\mathrm{cib}}$ is increasing in $b$

*Derivation.* This follows from the infinite sum expression for $p_{\mathrm{cib}}$ (equation A37). We will show that $p_{\mathrm{cib}}(b+1) > p_{\mathrm{cib}}(b)$ for integers $b \geq 1$ by showing that $p_{\mathrm{cib}}(b+1) - p_{\mathrm{cib}}(b) > 0$

$$\begin{aligned} p_{\mathrm{cib}}(\kappa_w, v, b) &= e^{-\bar{w}} \Big[ \sum_{j=0}^{b-1} \frac{\bar{w}^{j}}{j!} 1 + \sum_{j=b}^{\infty} \frac{\bar{w}^{j}}{j!} \frac{b}{j+1} \Big] \\ p_{\mathrm{cib}}(\kappa_w, v, b+1) &= e^{-\bar{w}} \Big[ \sum_{j=0}^{b} \frac{\bar{w}^{j}}{j!} 1 + \sum_{j=b+1}^{\infty} \frac{\bar{w}^{j}}{j!} \frac{b+1}{j+1} \Big] \\ &= e^{-\bar{w}} \Big[ \sum_{j=0}^{b-1} \frac{\bar{w}^{j}}{j!} 1 + \sum_{j=b}^{\infty} \frac{\bar{w}^{j}}{j!} \frac{b+1}{j+1} \Big] \end{aligned} \tag{A44}$$

This uses the fact that $\frac{b+1}{j+1} = 1$ when $j = b$.

For any $0 \leq \kappa_w \leq 1$, any $v \geq 1$, and $b \geq 1$, consider $p_{\mathrm{cib}}(b+1) - p_{\mathrm{cib}}(b)$:

$$\begin{aligned} p_{\mathrm{cib}}(b+1) - p_{\mathrm{cib}}(b+1) &= e^{-\bar{w}} \Big[ \sum_{j=b}^{\infty} \frac{\bar{w}^{j}}{j!} \frac{(b+1) - b}{j+1} \Big] \\ &= e^{-\bar{w}} \Big[ \sum_{j=b}^{\infty} \frac{\bar{w}^{j}}{j!} \frac{1}{j+1} \Big] \\ &> 0 \end{aligned} \tag{A45}$$

since this is a sum of positive terms, multiplied by a positive number.

■

#### 9.9 $p_{\text{surv}}/p_{\text{drift}}$ is decreasing in $b$

*Derivation.* We follow an approach similar to 9.8, showing that $\frac{p_{\text{surv}}(b+1)}{p_{\text{drift}}(b+1)} - \frac{p_{\text{surv}}(b)}{p_{\text{drift}}(b)} < 0$ for integer $b \geq 1$.

$$\begin{aligned}\frac{p_{\text{surv}}(b)}{p_{\text{drift}}(b)} &\approx \frac{v}{b}(1-\kappa_m)p_{\text{cib}} \\ &= \frac{v}{b}(1-\kappa_m)e^{-\bar{w}}\Big[\sum_{j=0}^{b-1}\frac{\bar{w}^j}{j!}1 + \sum_{j=b}^{\infty}\frac{\bar{w}^j}{j!}\frac{b}{j+1}\Big] \\ &= v(1-\kappa_m)e^{-\bar{w}}\Big[\sum_{j=0}^{b-1}\frac{\bar{w}^j}{j!}\frac{1}{b} + \sum_{j=b}^{\infty}\frac{\bar{w}^j}{j!}\frac{1}{j+1}\Big]\end{aligned}$$

Similarly:

$$\begin{aligned}\frac{p_{\text{surv}}(b+1)}{p_{\text{drift}}(b+1)} &\approx \frac{v}{b+1}(1-\kappa_m)e^{-\bar{w}}\Big[\sum_{j=0}^{b}\frac{\bar{w}^j}{j!}1 + \sum_{j=b+1}^{\infty}\frac{\bar{w}^j}{j!}\frac{b+1}{j+1}\Big] \\ &= \frac{v}{b+1}(1-\kappa_m)e^{-\bar{w}}\Big[\sum_{j=0}^{b-1}\frac{\bar{w}^j}{j!}1 + \sum_{j=b}^{\infty}\frac{\bar{w}^j}{j!}\frac{b+1}{j+1}\Big] \\ &= v(1-\kappa_m)e^{-\bar{w}}\sum_{j=0}^{b-1}\frac{\bar{w}^j}{j!}\frac{1}{b+1} + \sum_{j=b}^{\infty}\frac{\bar{w}^j}{j!}\frac{1}{j+1}\end{aligned}$$

We have again used the fact that $\frac{b+1}{j+1} = 1$ when $j = b$.

Taking the difference of the approximate ratios:

$$\frac{p_{\text{surv}}(b+1)}{p_{\text{drift}}(b+1)} - \frac{p_{\text{surv}}(b)}{p_{\text{drift}}(b)} \approx v(1-\kappa_m)e^{-\bar{w}}\sum_{j=0}^{b-1}\frac{\bar{w}^j}{j!}\left(\frac{1}{b+1} - \frac{1}{b}\right) < 0$$

That the expression is negative follows from the fact that all terms in the sum are negative, since $\frac{1}{b+1} < \frac{1}{b}$ for $b \geq 1$ while $\frac{\bar{w}^j}{j!}$ is always positive. The terms multiplying the sum are all positive.

It follows that $p_{\text{surv}}/p_{\text{drift}}$ is decreasing in $b$ ■

#### 9.10 In the absence of selection, the probability of surviving the bottleneck is approximately linear for large $v$ and small $f_{\text{mt}}$

*Derivation.* If $\kappa_w = \kappa_m = 0$, we have:

$$\begin{aligned}p_{\text{inoc}} &= (1 - e^{-vf_{\text{mt}}}) \\ \bar{w} &= v(1-f_{\text{mt}}) \approx v\end{aligned}$$

$$e^{-\bar{w}} \approx 0$$

$$p_{\text{cib}} \approx \frac{b}{v}$$

$$p_{\text{surv}} \approx (1 - e^{-vf_{\text{mt}}})(\frac{b}{v})$$

And since $-vf_{\text{mt}}$ is small, we can use the linear approximation to the exponential:

$$p_{\text{surv}} \approx vf_{\text{mt}}\frac{b}{v} = bf_{\text{mt}}$$

■
